# A streamlined ABC extruder-repressor module drives multi-bacteriocin resistance in streptococci

**DOI:** 10.64898/2026.02.18.706565

**Authors:** Julien Damoczi, Louise Pigeolet, Céline Cassiers, Johann Mignolet, Pascal Hols

**Affiliations:** Biochemistry and Genetics of Microorganisms, Louvain Institute of Biomolecular Science and Technology, Université catholique de Louvain, Louvain-la-Neuve, Belgium; Department of Fundamental Microbiology, Faculty of Biology and Medicine, University of Lausanne, Lausanne, Switzerland

**Author notes:** Corresponding author: Pascal Hols.

**Keywords:** antimicrobial peptide, AMP, bacteriocin defense, bacteriocin resistance, ABC exporter, GntR, YtrA, transcriptional control, sensory system

## Abstract

Bacteria inhabiting competitive microbial environments must rapidly detect and neutralize antimicrobial peptides (AMPs) produced by rivals. The activation of defense pathways relies on dedicated sensors and complex regulatory cascades. Here, we uncover a streamlined, membrane-embedded mechanism in Gram-positive bacteria that directly links detection to transcriptional control. We show that a YtrA-family transcriptional repressor is regulated through an unconventional direct physical interaction with its cognate ABC transporter. In *Streptococcus salivarius*, the MbrAB efflux pump sequesters the YtrA-like repressor MbrR through a competitive binding mechanism. Upon bacteriocin sensing, MbrR shifts from promoter-proximal DNA repression sites to membrane-associated sequestration via interaction with MbrA, thereby activating the bacteriocin defense system. ATP binding by MbrA facilitates MbrR recruitment, whereas ATP hydrolysis promotes its release, providing a dynamic, flux-responsive feedback loop finely tuned to environmental threat. Structural modeling, synteny, and conservation analyses reveal that this transporter-mediated sequestration mechanism is highly conserved across Gram-positive bacteria, suggesting a widespread and efficient strategy for AMP resistance that bypasses the need for classical sensor kinases. These findings expand the known repertoire of bacterial sensing and resistance systems and provide new insight into how Gram-positive bacteria swiftly adapt to interbacterial antagonism.

**IMPORTANCE:** Bacteria in crowded microbial communities face constant chemical warfare, yet how they sense and counteract antimicrobial peptides (AMPs) remains incompletely understood. Here we uncover a minimalist sensing-response module in streptococci in which YtrA-family transcriptional repressors are directly regulated by their partner ABC transporters. In *Streptococcus salivarius*, the MbrAB transporter detects incoming bacteriocins and physically sequesters the MbrR repressor, triggering rapid induction of a multi-gene defense program. ATP binding drives MbrR capture, while hydrolysis resets the system, providing a fast, energy-coupled switch that bypasses canonical two-component signaling. Comparative genomics shows that this transporter-repressor circuit is highly conserved across *Bacillota*, pointing to a broadly distributed and efficient strategy for AMP resistance. These findings reveal a direct physical link between membrane transport and transcriptional control, redefining how Gram-positive bacteria can sense and respond to microbial threats.

**ONE SENTENCE SUMMARY:** We identify a conserved transporter-repressor module in streptococci that directly couples bacteriocin detection to transcriptional activation, revealing a minimalist and rapid mechanism for antimicrobial peptide resistance.

## INTRODUCTION

Microbial communities are shaped by intense competition for nutrients and spatial niches [1]. Such competition is particularly prominent in the human oral cavity, one of the most densely colonized microbial ecosystems, where bacterial species continuously engage in cooperative and antagonistic interactions [2–4]. These interactions drive community structure and stability, influencing both health and disease [4]. Among the most potent weapons deployed during microbial competition are antimicrobial peptides (AMPs) known as bacteriocins, which are widely produced across bacterial taxa [5–7]. By selectively eliminating competitors while sparing the producing strain through dedicated immunity mechanisms, bacteriocins play a central role in determining microbial population dynamics [8].

To survive in bacteriocin-rich environments, bacteria have evolved multiple resistance strategies. Some species display intrinsic resistance to specific bacteriocins, whereas susceptible populations can rapidly acquire resistance following exposure [9]. Although bacteriocin resistance mechanisms often differ mechanistically from those conferring antibiotic resistance, they converge toward similar protective strategies, including remodeling of the cell envelope, antimicrobial inactivation or sequestration, and active efflux mediated by detoxification transporters [9]. In Gram-positive bacteria, resistance frequently involves modulation of cell envelope properties, such as reduced surface charge through D-alanylation of teichoic acids [10, 11], increased cell wall thickness that restricts access to the cytoplasmic membrane [10, 12], or decreased expression of membrane receptors required for bacteriocin activity [13–15]. Additional mechanisms directly target bacteriocins through proteolytic degradation [16, 17], extracellular sequestration [18, 19], or active removal from the membrane by ATP-binding cassette (ABC) transporters [20, 21].

Despite extensive characterization of these resistance mechanisms, a fundamental question remains unresolved: how bacteria rapidly detect extracellular AMPs and coordinate this detection with activation of defense responses. Most antimicrobial sensing pathways rely on multi-component regulatory cascades, particularly two-component systems (TCSs), which transduce extracellular signals into transcriptional responses through membrane-bound sensor kinases and cytoplasmic response regulators [22, 23]. Other regulatory circuits, including RNPP and GntR family regulators, also modulate bacteriocin production and resistance in response to environmental cues [24–26]. While these systems provide regulatory flexibility, they require multiple intermediate steps, potentially constraining the speed, sensitivity, and energetic efficiency of stress responses. Whether bacteria can directly couple antimicrobial detection to transcriptional control through more streamlined mechanisms remains largely unknown.

Such efficiency is particularly critical because antimicrobial production and resistance systems often impose substantial fitness costs [27–29]. Constitutive or poorly regulated activation can compromise growth and competitive fitness, especially in densely populated ecosystems where metabolic resources are limited. These constraints create strong selective pressure for regulatory circuits capable of tightly synchronizing environmental sensing with rapid and energetically economical gene activation. Identifying such mechanisms is therefore essential to understanding how bacteria balance competitive defense with metabolic economy in complex microbial communities.

*Streptococcus salivarius* is a dominant commensal of the human oral microbiota and contributes to maintaining microbial homeostasis within the oral cavity [2]. This species is notable for producing a remarkably diverse repertoire of bacteriocins [26, 30, 31]. Recent genomic and functional analyses revealed at least 21 distinct bacteriocin groups that are produced in various combinations among *S. salivarius* strains, highlighting the intense antimicrobial competition shaping its ecological niche [30]. Among these peptides, BlpK displays broad and potent activity against Gram-positive bacteria, including multidrug-resistant pathogens [30]. The widespread distribution of BlpK among *Streptococcus* species, together with the limited understanding of bacteriocin resistance mechanisms in this genus [24, 32–34], provides a unique opportunity to uncover fundamental principles governing AMP sensing and defense.

Here, we investigated resistance mechanisms to BlpK in *Streptococcus salivarius* by isolating spontaneous bacteriocin-resistant mutants and combining whole-genome sequencing with transcriptomic analyses. We identify an unconventional regulatory strategy in which a membrane-associated ABC transporter functions as both an AMP sensor and a regulatory hub. Upon bacteriocin exposure, this transporter directly interacts with and sequesters a transcriptional repressor, thereby coupling antimicrobial detection to activation of resistance genes without the involvement of canonical signal transduction pathways. These findings reveal a previously unrecognized mode of bacterial environmental sensing and suggest that transporter-mediated control of transcription may represent a widespread and efficient strategy for responding to interbacterial antagonism.

## RESULTS

### Inactivation of a YtrA-like regulator confers broad-spectrum resistance to bacteriocins

To identify genetic determinants of *S. salivarius* involved in bacteriocin resistance, we isolated spontaneous resistant mutants using agar plates supplemented with a chemically synthesized mature form of salivaricin BlpK (sBlpK) [30]. To avoid selecting mutations that increase expression of BlpK immunity proteins, we used a derivative of strain HSISS4 lacking the entire *blpK* operon, including putative immunity genes (Δ*blpK–blpI*, hereafter referred to as the wild type) [26]. From plates containing a gradient of sBlpK concentrations (0–10 µg/ml), we isolated 19 spontaneous mutants and performed whole-genome sequencing (WGS). Genomic analysis revealed multiple single nucleotide polymorphisms (SNPs) and insertions/deletions (InDels) (Table S1). Notably, approximately 75% of the mutants carried mutations in either a gene encoding a YtrA-like GntR-family transcriptional repressor, here named *mbrR* (<u>m</u>ultiple <u>b</u>acteriocin <u>r</u>esistance regulator), or in the adjacent gene *mbrB*, which encodes the membrane component of a predicted ABC transporter (named MbrAB) (Fig. 1A). Five *mbrR* missense mutations were identified: MbrR-N49I and R72P through WGS, and S25F, A54V, and T69I through targeted sequencing of *mbrR* in 21 additional spontaneous mutants (Fig. 1A and Table S1). However, most of *mbrR* mutants contained premature stop codons, suggesting partial or complete loss of MbrR function (Table S1). In addition, two distinct mutations were identified in *mbrB* at adjacent positions: an MbrB-R14P substitution and an in-frame serine insertion at position 15 (Fig. 1A).

**Fig. 1.**
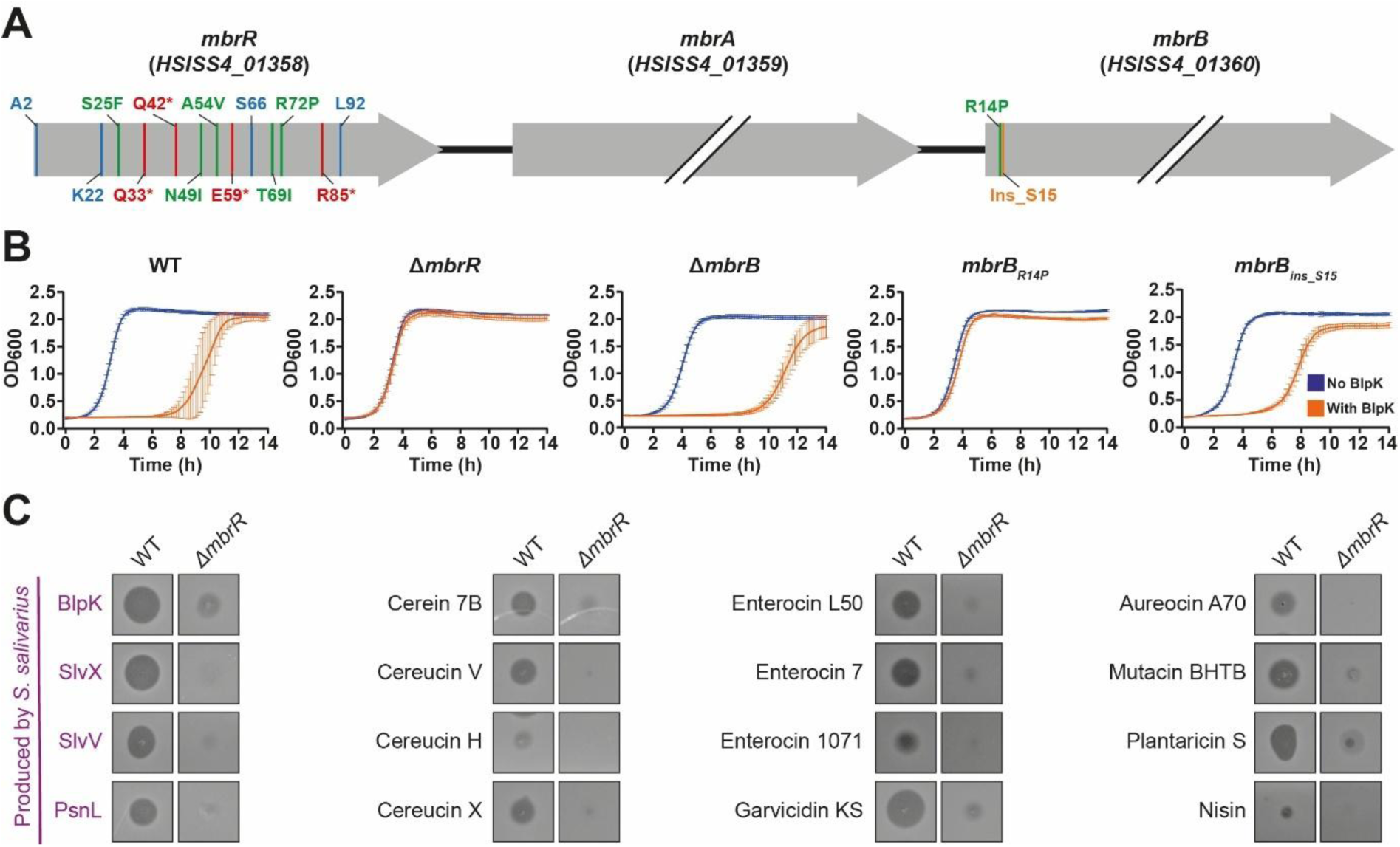
Impact of mutations in MbrR and MbrB on bacteriocin resistance. **A.** Scheme of the *mbrRAB* operon and mapping of identified mutations. Mutations are color-coded: blue for frameshifts, green for amino-acid substitution, red for premature stops (*), and orange for in-frame insertion. Except for insertion, the amino acid refers to the mutated codon. Double slashes indicate genes not drawn to scale. **B.** Impact of BlpK on the growth of the wild type and Δ*mbrR*, Δ*mbrB*, *mbrB_R14P_*, *mbrB_ins-S15_* mutant strains. Growth was performed in M17G with (orange) or without (blue) sBlpK (10 µg/ml). Each curve is a mean (± SD) of three independent biological replicates. **C.** Contribution of MbrR to general bacteriocin resistance on plates. 2 µl of each synthetic bacteriocin was spotted. For clarity, bacteriocin cocktails composed of multiple peptides are referred to the generic name of the corresponding bacteriocin. Additional details on concentration and solubilization are available in the Methods section.

To evaluate the contribution of these mutated genes to sBlpK resistance, a representative allele was individually backcrossed into the wild-type background (Fig. 1B and Fig. S1). Growth assays performed in the presence and absence of sBlpK demonstrated that deletion of *mbrR*, the *mbrB* mutations, and a mutation at genomic position 1,803,533 all conferred resistance to sBlpK (Fig. 1B and Fig. S1).

We next evaluated the resistance spectrum of the Δ*mbrR* mutant using a panel of bacteriocins (Fig. 1C) comprising 65 chemically synthesized peptides from the PARAGEN collection [35] and the five salivaricins produced by *S. salivarius* HSISS4 (Dataset S1). Of the 70 bacteriocins tested, approximately 20 displayed moderate to high activity against *S. salivarius*. In most cases, deletion of *mbrR* conferred resistance to these bacteriocins, indicating that MbrR controls a broad-spectrum bacteriocin resistance mechanism (Fig. 1C). In contrast, no significant differences in susceptibility were observed between wild-type and Δ*mbrR* strains when tested against multiple classes of antibiotics (Table S2), indicating that the MbrR-dependent resistance mechanism is largely specific to bacteriocins and does not extend to conventional antibiotics.

### MbrR is a transcriptional repressor of a multi-component bacteriocin defense system

To elucidate the regulatory mechanism underlying bacteriocin resistance, we compared the transcriptomes of wild-type and Δ*mbrR* mutant strains using RNA sequencing (RNA-seq). This analysis identified a set of nine genes that were significantly upregulated in the absence of *mbrR* (Fig. 2A and B, Table S3). Among these genes, seven encode components of three putative ABC efflux transporters (operons *mbrAB*, *mbrCD*, and *mbrEFG*), one encodes a flotillin-like protein (here named *mbrH*), and one encodes a predicted membrane-associated CaaX endoprotease (here named *mbrX*) (Fig. 2A).

**Fig. 2.**
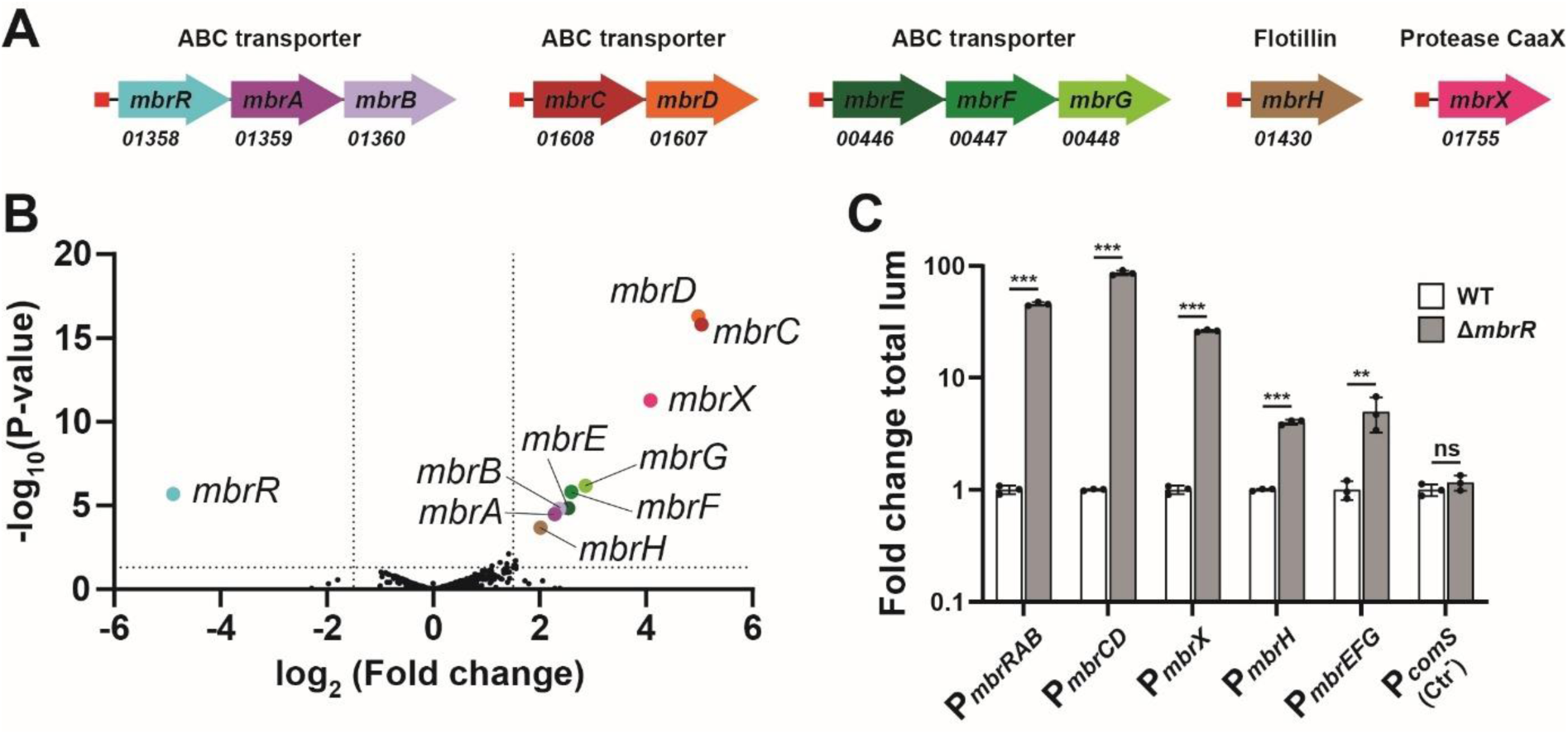
MbrR regulon. **A.** Transcriptional units (TUs) of the MbrR regulon. Red rectangles represent MbrR binding sites. TUs are color-coded: *mbrRAB* (blue to light purple arrow), *mbrCD* (red and orange arrow), *mbrEFG* (dark to light green arrow), *mbrH* (brown arrow), and *mbrX* (pink arrow). The HSISS4 locus tag is indicated below each arrow. **B.** Transcriptome (RNA-seq) comparison between wild-type (WT) and Δ*mbrR*. Black dots indicate genes for which non-significant changes were observed. Color dots indicate genes that are upregulated or downregulated. The horizontal dotted line represents a *P* value of 0.05, while the vertical dotted lines represent log_2_ fold change values of ≥1.5 and ≤−1.5. These results were obtained from two independent replicates. **C**. MbrR-dependent repression of promoters. Fold change in total luminescence (Fold change total lum) produced by each promoter fused to the *luxAB* genes in WT or Δ*mbrR* background. The promoter of *comS* (P*_comS_*) was used as negative control (Ctr^-^). Experimental values represent the average fold change (± SD) of three independent replicates; dots show the values of independent experiments (*n* = 3). Each replicate was compared to the mean value of the wild type used as reference. Statistical analysis was performed using *t* tests (ns, non-significant; **, *P* < 0.01; ***, *P* < 0.001).

To confirm the regulatory role of MbrR in controlling these transcriptional units, each promoter was fused to luciferase reporter genes (*luxAB*) and chromosomally integrated at an ectopic locus. Promoter activities were then compared in the presence or absence of MbrR. The promoter of *comS* was used as a negative control (Fig. 2C). Deletion of *mbrR* resulted in a strong increase in promoter activity, ranging from approximately 5- to 100-fold across all tested promoters, confirming that MbrR functions as a transcriptional repressor of this regulon.

### MbrR binds *in vitro* to a conserved palindromic sequence

To identify potential MbrR binding sites, we aligned the promoter regions of transcriptional units upregulated in the Δ*mbrR* mutant and identified a conserved palindromic sequence as a candidate regulatory motif (Fig. 3A and B). To further support this prediction, we used AlphaFold3 [36] to model the interaction between an MbrR dimer and the DNA sequence corresponding to the intergenic region upstream of *mbrR*. The predicted structure positioned the MbrR dimer directly over the inverted repeats identified in the promoter analysis (Fig. 3C and Fig. S2A).

**Fig. 3.**
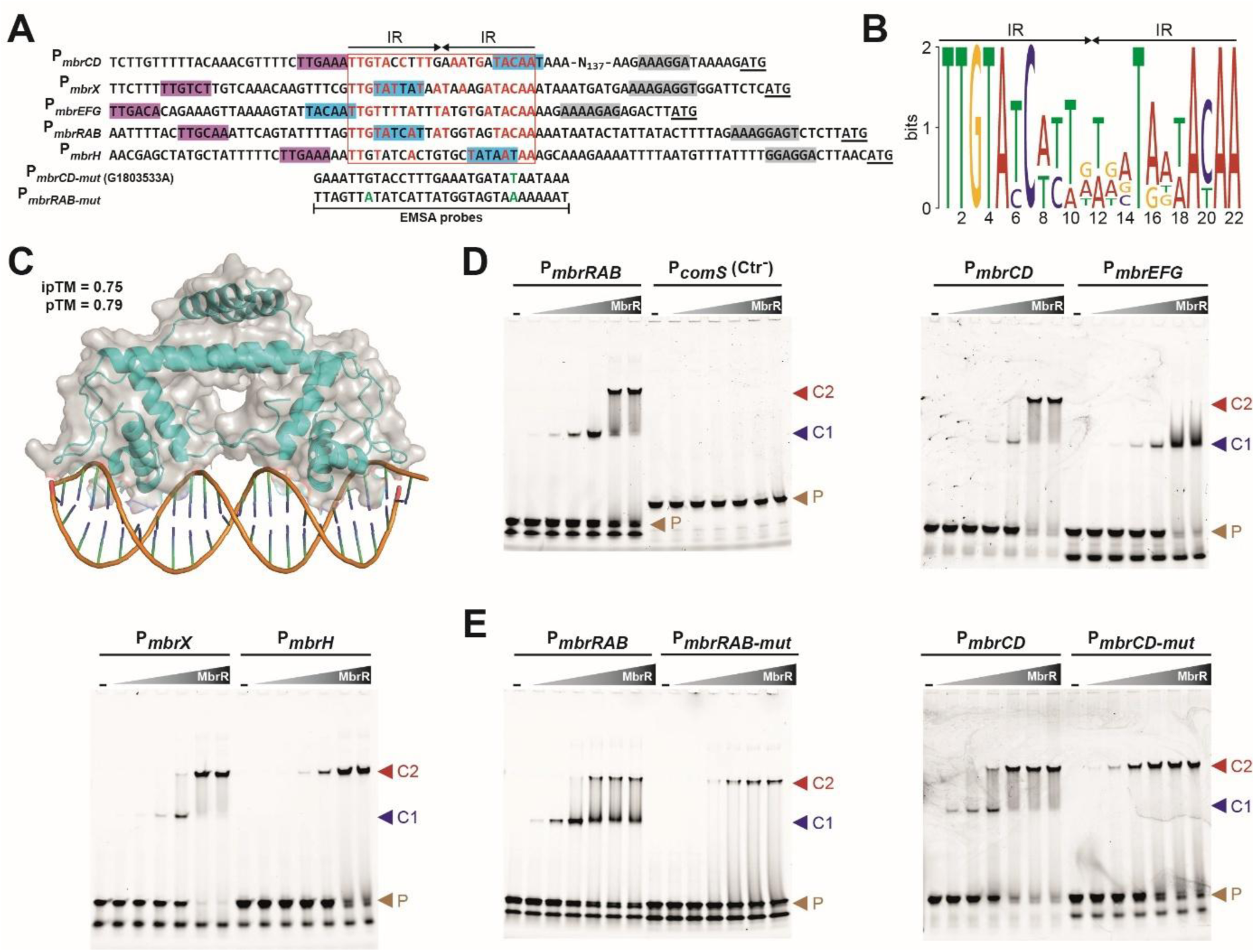
MbrR-binding site. **A.** Nucleotide alignment of P*_mbrRAB_*, P*_mbrCD_*, P*_mbrEFG_*, P*_mbrH_*, and P*_mbrX_*. Putative -35 boxes, -10 boxes, and RBS are highlighted in purple, blue, and gray, respectively. The ATG start codon is underlined and the MbrR binding site is surrounded by a red square in which matching nucleotides with the inverted repeats are in red. Black arrows indicate inverted repeat (IR) sequences. The line surrounded by vertical bars indicates probes used for electrophoretic mobility shift assays (EMSAs). **B.** Weighted consensus sequence of the MbrR-binding site. Black arrows indicate IR sequences. The logo was created by aligning the IR sequences of panel A (red rectangle) with the MEME suite online software. **C.** AlphaFold3 structure prediction of MbrR dimer (cyan) bound to the IR sequences (orange) located upstream of the *mbrRAB* operon. Scores of interface Predicted Template Modeling (iPTM) and Predicted Template Modeling (PTM) are indicated top left. **D.** *In vitro* binding of IR sequences by MbrR. EMSAs of P*_mbrRAB_*, P*_mbrCD_*, P*_mbrEFG_*, P*_mbrH_*, P*_mbrX_*, and P*_comS_* (negative control, Ctr^-^) probes conducted with increasing concentrations of purified MbrR (gray triangles; 1:4 dilutions from 4 µM). Probes of 30 bp were Cy3-conjugated and used at 40 ng. Arrowheads labeled P, C1, and C2 show the positions of the probe, C1 complex, and C2 complex, respectively. **E.** Impact of IR sequence mutations on MbrR binding. EMSAs of P*_mbrRAB_*, P*_mbrRAB-mut_*, P*_mbrCD_* and P*_mbrCD-mut_* probes conducted with increasing concentrations of purified MbrR (gray triangles; 2:2 dilutions from 2 µM). Probes of 30 bp were Cy3-conjugated and used at 40 ng. Arrowheads as in panel D.

To experimentally validate MbrR binding, we performed electrophoretic mobility shift assays (EMSAs) using a 30-bp DNA fragment encompassing the inverted repeats as a probe and increasing concentrations of purified MbrR protein (Fig. 3D and E, Fig. S2B). MbrR bound all probes containing the identified regulatory motif, whereas no binding was observed with the P*_comS_* probe used as a negative control (Fig. 3D). Depending on the promoter, two distinct complexes were detected: a high-affinity complex (C1, lower band) and/or a lower-affinity complex (C2, upper band). These results suggest that MbrR may bind DNA in multiple oligomeric states *in vitro*, consistent with observations reported for other members of the GntR family [37, 38] (Fig. 3D and Fig. S2C).

To assess binding specificity, we performed EMSAs using a mutated version of the *mbrR* promoter (P*_mbrRAB-mut_*) in which the highly conserved G3 and C20 residues within the palindromic motif were replaced by adenines (Fig. 3A and E). With the mutated probe, only the low-affinity complex C2 was detected, and no binding was observed at MbrR concentrations below 0.25 µM, whereas the wild-type probe exhibited detectable binding at concentrations as low as 4 nM (Fig. 3E). To evaluate the functional consequences of this mutation *in vivo*, we introduced a P*_mbrRAB-mut_-luxAB* reporter fusion into the wild-type background (Fig. S2D). Luciferase activity was strongly increased compared with the wild-type promoter, indicating reduced repression due to impaired MbrR binding.

In addition, WGS identified a guanine-to-adenine point mutation at genomic position 1,803,533 that conferred sBlpK resistance (Fig. S1 and Table S1). This mutation (C to T on the complementary strand) lies within the predicted MbrR binding site upstream of *mbrCD* (Fig. 3A), suggesting disruption of MbrR-mediated regulation. Consistent with this hypothesis, EMSAs performed with the corresponding mutated probe (P*_mbrCD-mut_*) showed loss of the high-affinity complex C1 and reduced overall binding affinity (Fig. 3E). Furthermore, *in vivo* luminescence measurements of the P*_mbrCD-mut_* reporter fusion in the wild-type background were significantly higher than those observed with the wild-type promoter (Fig. S2D), confirming reduced MbrR binding to the mutated regulatory sequence.

Together, these *in vitro* and *in vivo* results demonstrate that MbrR directly represses transcription of genes encoding the Mbr defense system through binding to a conserved palindromic promoter motif.

### All components of the Mbr system contribute to bacteriocin resistance

To determine the contribution of individual components of the Mbr system to BlpK resistance, we generated single operon/gene deletions in a Δ*mbrR* background, in which the entire Mbr system is constitutively expressed. The sensitivity of each double mutant to sBlpK was then assessed using spot-on-lawn assays (Fig. 4A and Dataset S1) and growth analyses in liquid medium (Fig. 4B). Deletion of *mbrCD* in the Δ*mbrR* background restored sBlpK sensitivity to a level comparable to that of the wild-type strain, whereas deletion of other Mbr components did not significantly alter resistance (Fig. 4A and B). Notably, this observation is consistent with the previously identified mutation within the MbrR-binding site upstream of *mbrCD* (genomic position 1,803,533), which also confers sBlpK resistance (Fig. S1). Together, these results demonstrate that the MbrCD transporter is essential for resistance to BlpK. We next evaluated the contribution of individual Mbr components to resistance against a broader panel of bacteriocins. Resistance to specific bacteriocins was found to be component dependent (Fig. 4C and Dataset S1). Strikingly, resistance to several bacteriocins required the coordinated action of multiple Mbr components, and in some cases involved the entire system, such as resistance to plantaricin S and mutacin BHTB (Fig. 4C).

**Fig. 4.**
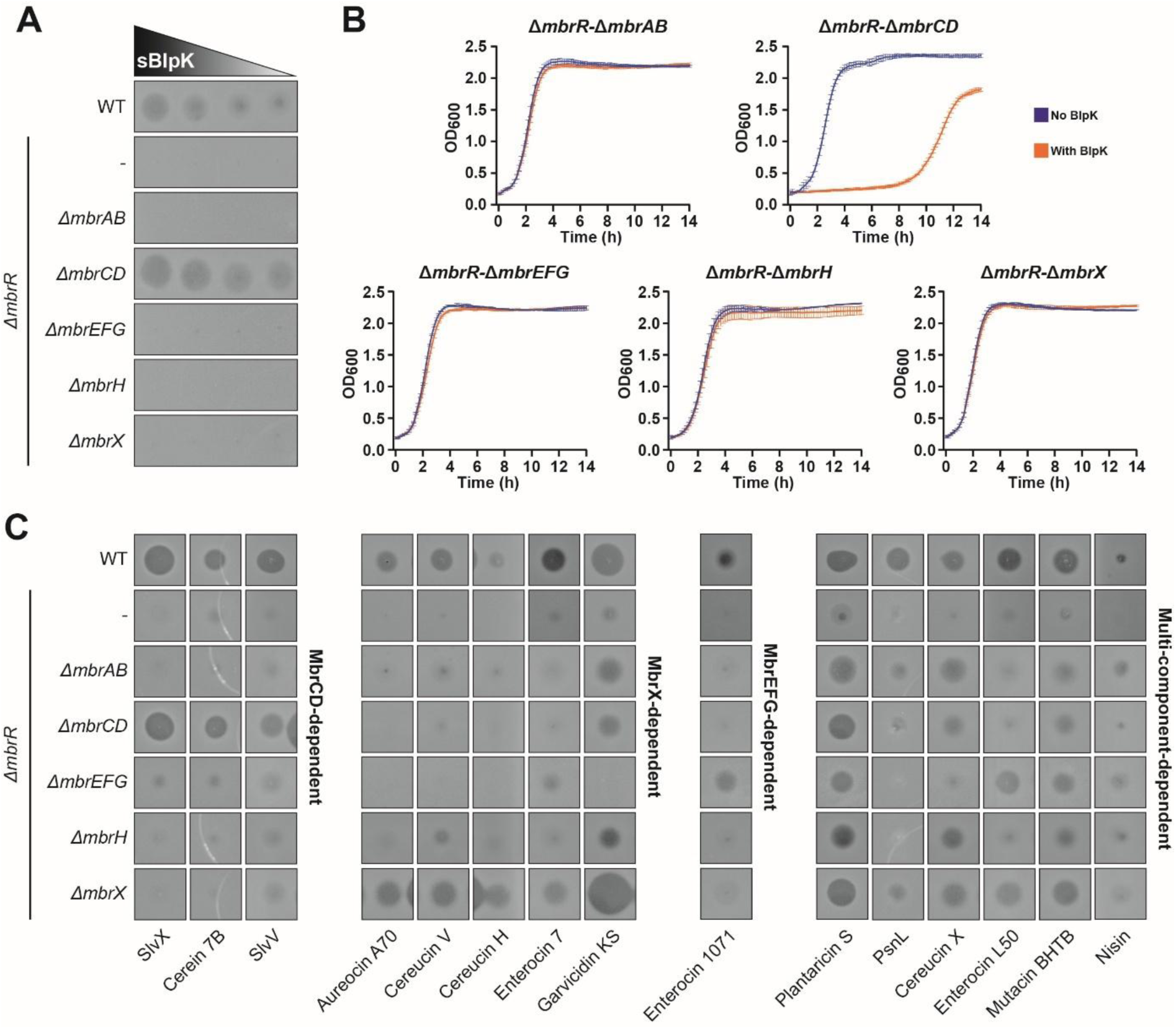
Contribution of each component of the Mbr system to bacteriocin resistance. **A.** Contribution of each Mbr component to BlpK resistance on plates. 2µl of a decreasing concentration of sBlpK (black to gray triangle; 2:2 dilutions from 500 µg/ml) were spotted. Each inhibition zone on the spot-on-lawn assay is representative of three independent experiments performed in the same conditions. **B.** Contribution of each Mbr component to BlpK resistance in liquid culture. Growth was performed in M17G liquid medium with sBlpK (10 µg/ml) (orange) or without (blue). Each curve is a mean (± SD) of three independent replicates. **C.** Contribution of Mbr components to resistance against various bacteriocins on plate. The experimental setup as described in figure 1C.

Collectively, these findings indicate that the Mbr system functions as a multi-component defense module in which each element contributes to broad-spectrum bacteriocin resistance in *S. salivarius*.

### The cytoplasmic ATPase component of MbrAB sequesters MbrR

Although inactivation of MbrB or the entire MbrAB transporter did not affect resistance to BlpK (Fig. 1B and Fig. 4A and B), two independent substitutions in the membrane component MbrB conferred resistance (Fig. 1A and B), suggesting an additional regulatory role for the MbrAB transporter. To further investigate this possibility, we assessed the bacteriocin resistance profile of the *mbrB_R14P_* mutant. This mutant displayed a resistance spectrum nearly identical to that of the Δ*mbrR* mutant, indicating that the MbrB-R14P substitution may relieve MbrR-mediated repression (Fig. S3 and Dataset S1). To test this hypothesis, we monitored expression of the P*_mbrRAB_*-*luxAB* reporter fusion in the *mbrB_R14P_* background (Fig. 5A). The R14P substitution resulted in strong derepression of the *mbrRAB* operon. Notably, in a strain lacking the cytoplasmic ATPase MbrA (Δ*mbrA*, markerless deletion) but expressing MbrB-R14P, *mbrRAB* expression returned to wild-type levels (Fig. 5A), indicating that MbrA is required for MbrR derepression.

**Fig. 5.**
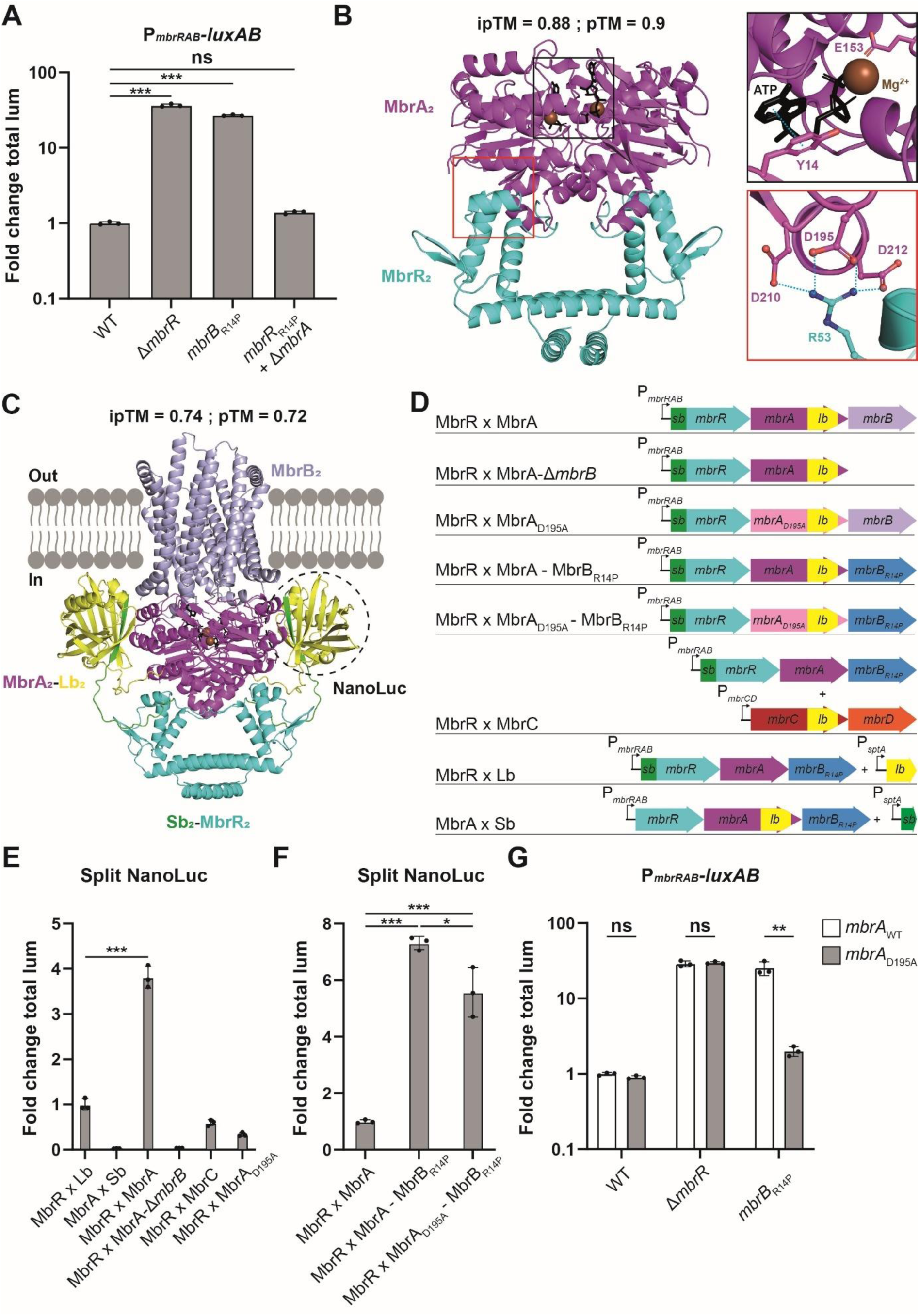
Specific interaction between MbrR and MbrB. **A.** Impact of the substitution MbrB-R14P on P*_mbrRAB_* activation. Fold change in total luminescence (Fold change total lum) produced by the P*_mbrRAB_*-*luxAB* fusion in WT (negative control used as reference), Δ*mbrR*, *mbrB_R14P_*, or *mbrB_R14P_* Δ*mbrA* background. **B.** AlphaFold3 structure prediction of the tetrameric complex MbrR_2**·**_MbrA_2_ in presence of 2 Mg^2+^ and 2 ATP. Black rectangle shows the ATP binding site and residues Y14 and E153 involved in ATP binding and hydrolysis, respectively. Red rectangle shows the network of salt bridges between MbrR-R53 and MbrA-D195-D210-D212. **C.** AlphaFold3 structure prediction of the hexameric complex MbrB_2**·**_MbrA_2_-Lb_2**·**_Sb_2_-MbrR_2_ in presence of 2 ATP and 2 Mg^2+^. The reconstitution of the NanoLuc by Sb-Lb interaction is shown. In panels B and C, scores of iPTM and PTM are indicated at the top of the structure. **D.** Schematic diagram of constructs used for split-NanoLuc assays. Insertion of fragments encoding Sb or Lb were performed at the native loci. For negative controls, the production of Sb or Lb alone under the control of P*_sptA_* (sBI7 inducible) was achieved from an ectopic chromosomal locus. In panels B, C, and D, proteins, genes and molecules are color-coded: cyan for MbrR/*mbrR,* dark purple for MbrA/*mbrA,* pink for *mbrA_D195A_,* light purple for MbrB/*mbrB,* dark blue for *mbrB_R14P_,* red for *mbrC*, and orange for *mbrD,* green for Sb/*sb*, yellow Lb/*lb*, black for ATP, and brown for Mg^2+^. **E.** Split-NanoLuc assays for the validation of MbrR-MbrA interaction in WT. Fold change in total luminescence produced by Sb-MbrR × Lb (negative control used as reference), MbrA-Lb × Sb (negative control), Sb-MbrR × MbrA-Lb, Sb-MbrR × MbrA-Lb (Δ*mbrB*), Sb-MbrR × MbrC-Lb (negative control), and Sb-MbrR × MbrA_D195A_-Lb. **F.** Split-NanoLuc assays for the validation of MbrR-MbrA interaction in MbrB_R14P_ background. Fold change in total luminescence produced by Sb-MbrR × MbrA-Lb in WT (positive control used as reference), Sb-MbrR × MbrA-Lb with MbrB_R14P_, and Sb-MbrR × MbrA_D195A_-Lb with MbrB_R14P_. **G**. Impact of the mutation MbrA-D195A on P*_mbrRAB_* activation. Fold change in total luminescence produced by the P*_mbrRAB_*-*luxAB* fusion in WT, Δ*mbrR* or *mbrB_R14P_* background. In panel A, each replicate was compared to the mean value of the wild type. Experimental values represent the average fold change (± SD) of three independent replicates. One-way ANOVA with Dunnett’s test and *t* test were performed for data in panels A/F and E/G, respectively (ns, non-significant; *, *P* < 0.05; **, *P* < 0.01; ***, *P* < 0.001).

To explore potential physical interactions among MbrR, MbrA, and MbrB, we generated structural models using AlphaFold3 [36] (Fig. S4). Among the predicted complexes, MbrR_2_**·**MbrA_2_ tetrameric and MbrR_2_**·**MbrA_2_**·**MbrB_2_ hexameric assemblies (in the presence of ATP and Mg²⁺) supported a direct MbrR-MbrA interaction with high confidence scores (ipTM and pTM: ∼0.9) (Fig. S4A and Fig. 5B). In contrast, models predicting direct interaction between MbrR and MbrB yielded low confidence score (ipTM and pTM: ∼0.3), suggesting that MbrR primarily interacts with MbrA. Structural analysis further identified a potential salt-bridge network involving MbrR residue R53 and MbrA residues D195, D210, and D212 (Fig. 5B), suggesting a stabilizing interface between the two proteins. As a specificity control, we modeled equivalent complexes between MbrR and the paralogous MbrCD transporter. These models predicted weak interactions, as supported by predicted aligned error (PAE) heatmaps (Fig. S4B), indicating that MbrR-MbrA interaction is likely specific.

To experimentally validate this interaction, we employed a split NanoLuc luciferase assay, which measures protein-protein interactions through reconstitution of luciferase activity from complementary inactive fragments, the large bit (Lb, 156 aa) and small bit (Sb, 11 aa) [39, 40] (Fig. 5C). Multiple constructs were integrated at native chromosomal loci to assess MbrR-MbrA interactions (Fig. 5D). As negative controls, free Lb or Sb fragments were co-expressed with Sb-MbrR or MbrA-Lb, respectively, resulting in minimal luminescence signals (Fig. 5E). Co-expression of Sb-MbrR and MbrA-Lb in the wild-type background produced a luminescence signal approximately fourfold higher than the negative control, demonstrating interaction between MbrR and MbrA (Fig. 5E). This signal was markedly reduced in the absence of MbrB, indicating that MbrB promotes or stabilizes the MbrR-MbrA interaction (Fig. 5E). Strikingly, the interaction signal increased approximately eightfold in the *mbrB_R14P_* background (Fig. 5F), suggesting that the R14P substitution enhances complex formation. No significant luminescence was detected between MbrR and MbrC (Fig. 5F), confirming interaction specificity.

To assess the functional relevance of the predicted salt-bridge interface, we substituted MbrA residue D195 with alanine (MbrA-D195A) (Fig. 5B). In split NanoLuc assays, this mutation significantly reduced luminescence signals in both wild-type and *mbrB_R14P_* backgrounds (Fig. 5E and F), supporting a critical role for this residue in stabilizing MbrR-MbrA interaction. Consistent with these results, P*_mbrRAB_* activation assays showed that the MbrA-D195A mutation had minimal impact under basal conditions but significantly reduced operon activation in the *mbrB_R14P_* background (Fig. 5G), where MbrR recruitment to MbrA is enhanced.

Together, these results demonstrate a specific and direct interaction between MbrR and the ATPase component MbrA and support a model in which MbrA functions as a regulatory partner that modulates MbrR activity through sequestration. This mechanism reveals an unconventional mode of transcriptional control mediated by an ABC transporter.

### MbrAB acts as a bacteriocin sensor controlling MbrR activity

Given the essential role of the MbrAB transporter in resistance to multiple bacteriocins (Fig. 4C), we hypothesized that MbrAB might directly sense antimicrobial peptides and, in response, trigger sequestration of MbrR. To test this possibility, we monitored the activity of a P*_mbrRAB_*-*luxAB* reporter fusion in wild-type and Δ*mbrR* backgrounds following exposure to increasing concentrations of four bacteriocins: BlpK, enterocin 1071, cereucin X, and PsnL (Fig. S5). Cereucin X and PsnL were selected based on the strong hypersensitivity of the Δ*mbrAB-*Δ*mbrR* mutant relative to the Δ*mbrR* mutant, suggesting that these bacteriocins may directly interact with the MbrAB transporter (Fig. 4C).

Among the tested bacteriocins, PsnL and cereucin X induced significant derepression of *mbrRAB* expression in the wild-type background (up to ∼5-fold and ∼25-fold, respectively), whereas no induction was observed in the Δ*mbrR* strain, demonstrating MbrR-dependent regulation (Fig. S5). Cereucin X is composed of three peptides (A, B, and C), and subsequent analysis identified peptide B (cereucin X-B) as the component responsible for transcriptional derepression (Fig. S6A). Consistent with these findings, both PsnL and cereucin X-B activated multiple MbrR-controlled promoters (Fig. S6B and C). Dose-response analyses revealed that derepression of *mbrRAB* occurs at subinhibitory concentrations of PsnL and cereucin X-B, with minimal impact on bacterial growth (Fig. 6A and B), indicating that MbrAB-mediated sensing precedes overt antimicrobial toxicity. Importantly, bacteriocin-dependent induction was absent or strongly reduced in Δ*mbrR*, *mbrA_D195A_*, and *mbrB_R14P_* mutants (Fig. 6A and B and Fig. S6D and E), demonstrating that each component of the MbrRAB system is required for signal transmission. Notably, the *mbrB_R14P_* variant exhibited constitutive derepression and failed to respond to bacteriocin exposure, suggesting that this mutation locks the system in an activated state. Structural modeling supports this interpretation, predicting that the proline substitution alters the conformation and flexibility of the MbrB bacteriocin-binding cavity (Fig. S7).

**Fig. 6.**
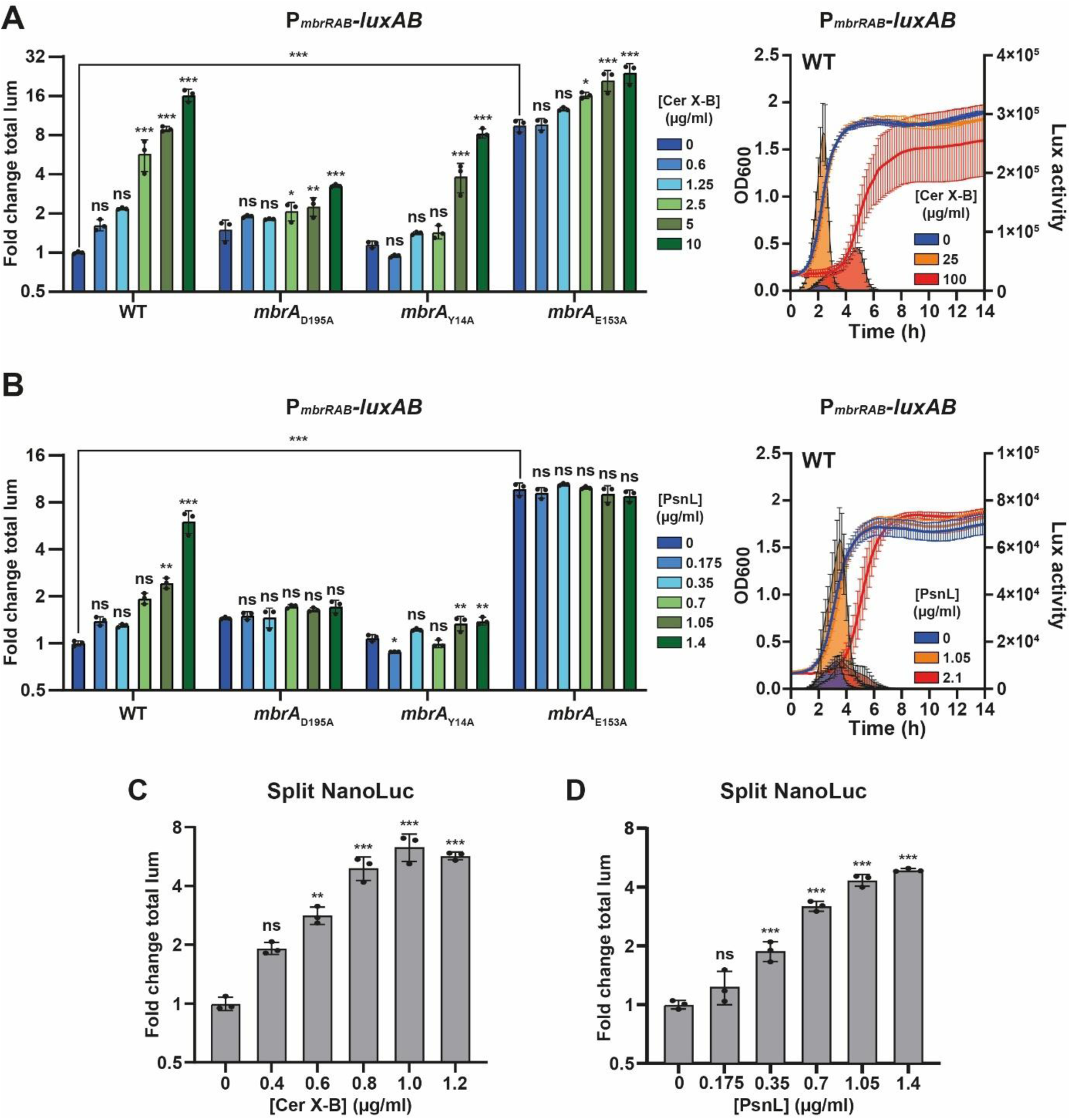
Induction of *mbrRAB* expression by bacteriocin addition. **A.** P*_mbrRAB_* activation in response to increasing concentration of cereucin X-B (Cer X-B). (left) Fold change in total luminescence (Fold change total lum) produced by the P*_mbrRAB_*-*luxAB* fusion with 0 to 10 μg/ml of Cer X-B in WT, *mbrA_D195A_*, *mbrA_Y14A_* or *mbrA_E153A_* background. (right) Growth (OD_600_) and luminescence (Lux activity, RLU/OD_600_) at subinhibitory and inhibitory concentrations of Cer X-B (20 and 100 μg/ml, respectively) in WT. **B.** P*_mbrRAB_* activation in response to increasing concentration of PsnL. (left) Genetic backgrounds as in panel A and tested PsnL concentrations from 0 to 1.4 μg/ml. (right) Setup as in panel A with 1.05 and 2.1 μg/ml of PsnL. **C.** and **D.** MbrR-MbrA interaction in response to increasing concentration of Cer X-B (panel C, 0 to 1.2 μg/ml) and PsnL (panel D, 0 to 1.4 μg/ml). Fold change in total luminescence produced by Sb-MbrR × MbrA-Lb in WT. Each replicate was compared to the mean of the WT without bacteriocin (negative control used as reference). Experimental values represent the average fold change (± SD) of three independent replicates. One-way ANOVA with Dunnett’s test were performed (ns, non-significant; *, *P* < 0.05; **, *P* < 0.01; ***, *P* < 0.001).

To directly test whether bacteriocin sensing modulates MbrR recruitment, we quantified MbrR-MbrA interaction using split NanoLuc assays. Addition of cereucin X-B or PsnL induced a dose-dependent increase in luminescence, demonstrating that bacteriocin exposure enhances MbrR**·**MbrA complex formation (Fig. 6C and D).

Together, these results demonstrate that MbrAB functions not only as an efflux transporter but also as a bacteriocin-responsive sensor that physically recruits MbrR in response to subinhibitory concentrations of antimicrobial peptides. This dual function reveals a direct coupling between antimicrobial detection and transcriptional regulation.

### ATP binding and hydrolysis by MbrA dynamically controls MbrR

The above results suggest that the membrane component MbrB directly engages bacteriocins and transmits a conformational change to the cytoplasmic ATPase MbrA, which in turn recruits MbrR. To dissect the dynamics of this process, we tested whether ATP binding and hydrolysis by MbrA are required for MbrR sequestration and/or release.

Structural modeling using AlphaFold3 [36] indicated a strong influence of ATP on the MbrR_2_**·**MbrA_2_ tetrameric complex. Inclusion of two ATP molecules increased the confidence scores (ipTM-pTM: 0.88-0.90 vs 0.52-0.59 without ATP) and promoted compaction of the MbrA dimer, suggesting that ATP stabilizes the MbrR-MbrA interface and strengthens overall complex formation (Fig. S4).

To validate this experimentally, we generated MbrA mutants targeting conserved and key residues for ATP binding (MbrA-Y14A) and ATP hydrolysis (MbrA-E153A) based on prior studies of ABC transporters [41–43] (Fig. 5B). These mutations were introduced into the wild-type background carrying the P*_mbrRAB_*-*luxAB* reporter fusion. Upon exposure to cereucin X-B or PsnL, the MbrA-Y14A mutant displayed a lower reporter induction compared to wild type, indicating that ATP binding is required for MbrR recruitment. In contrast, the MbrA-E153A mutant exhibited strong derepression (∼10-fold) even in the absence of bacteriocins, suggesting that ATP hydrolysis is necessary for MbrR release (Fig. 6A and B).

Together, these findings show that MbrA dynamically controls MbrR through its ATPase cycle: ATP binding promotes MbrR recruitment, while ATP hydrolysis facilitates its subsequent release. This mechanistic insight reveals an unprecedented, energy-dependent regulatory layer in ABC transporter-mediated transcriptional control.

## DISCUSSION

Resistance to AMPs in bacteria frequently arises through mutations in regulatory circuits that control efflux systems [28, 29, 44] or cell envelope biosynthesis pathways [12]. Although upregulation of these defense mechanisms enhances survival under antimicrobial stress, their constitutive activation typically incurs substantial fitness costs [28, 29], necessitating regulatory strategies that ensure rapid but conditional deployment [27].

In this study, we describe an unconventional regulatory paradigm in which a membrane-associated ABC transporter, MbrAB, functions as both sensor and effector by mediating sequestration of the YtrA-like transcriptional repressor MbrR. This mechanism promotes derepression of the Mbr multi-bacteriocin resistance system upon AMP exposure (Fig. 7A). A notable feature of this defense module is the coordinated activation of three mechanistically distinct effectors that may function independently or synergistically depending on the biochemical nature of the antimicrobial peptide encountered. While ABC transporters and CaaX proteases are frequently associated with bacteriocin resistance systems, [9, 16] the involvement of a flotillin protein was unexpected. Flotillins are integral components of membrane microdomains and may modulate bacteriocin susceptibility indirectly, for instance by chaperoning membrane-associated proteases [45] or promoting membrane repair processes [46].

**Fig. 7.**
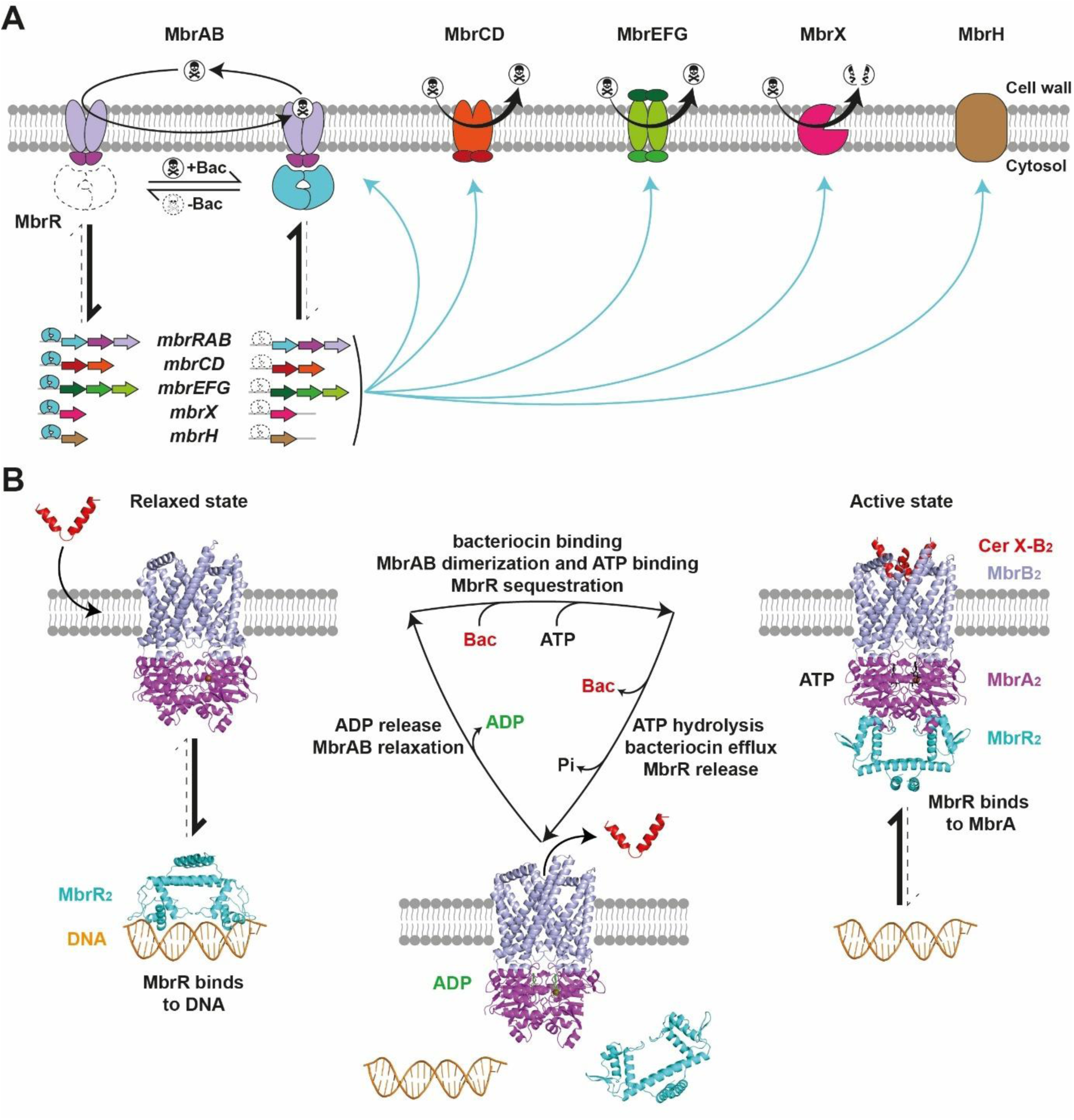
MbrR mode of action. **A.** Global scheme of the regulation of the Mbr system in *S. salivarius.* Black arrows represent movement or conformational change. Blue arrows represent protein production. The dead’s-head symbol represents bacteriocin (Bac). **B.** Model of the MbrRAB transduction system using AlphaFold3 modelled complexes. Bacteriocin interaction with MbrB induces its conformational change that is transmitted to MbrA. This favors MbrA dimerization, ATP binding, and MbrR recruitment. Bacteriocin efflux and ATP hydrolysis will lead to MbrR release and MbrAB relaxation to its basal state for the next cycle. The AlphaFold3 structure prediction of cereucin X-B was used to represent bacteriocin. In panels A and B, genes, proteins, and molecules are color-coded: *mbrRAB*/MbrRAB (cyan to light purple), *mbrCD*/MbrCD (red and orange), *mbrEFG*/MbrEFG (dark to light green), *mbrH*/MbrH (brown), mbrX/MbrX (pink arrow), orange for DNA, red for bacteriocin, black for ATP, green for ADP, and brown dot for Mg^2+^.

Our data support a model in which MbrRAB represents a previously unrecognized signal transduction architecture, whereby a membrane-bound ABC transporter acts as a proxy environmental sensor and transduces antimicrobial signals through its ATP-binding and hydrolysis cycle to regulate transcription. Integrating our experimental findings with established mechanistic knowledge of ABC transporter function [47, 48], we propose that translocation of bacteriocins into the membrane cavity of MbrB induces conformational rearrangements that are transmitted to the MbrA ATPase, thereby promoting ATP binding and recruitment of MbrR to the membrane. Subsequent bacteriocin extrusion coupled to ATP hydrolysis would facilitate MbrR release, restoring basal repression and resetting the regulatory cycle (Fig. 7B). This regulatory strategy bypasses classical activation mechanisms involving direct ligand binding or covalent modification of transcription factors. Instead, it enables a rapid, reversible, and energetically efficient means of signal transmission, as ATP hydrolysis simultaneously drives antimicrobial efflux and transcriptional regulation. Interestingly, the MbrRAB system shares several conceptual features with peptide-mediated quorum-sensing pathways in Gram-positive bacteria [49], including sensitivity to extracellular peptide concentration, dose-dependent activation, and autoregulatory feedback through control of *mbrRAB* expression. However, in contrast to canonical quorum sensing systems, MbrRAB does not rely on a dedicated autoinducing peptide. Rather, it appears to function as a quorum-responsive defensive module that senses the accumulation of AMPs produced by competing microbial populations.

Comparative genomic analyses revealed that the MbrRAB system is broadly distributed among streptococci, with the notable exception of species within the sanguinis group and subsets of the mitis group (Fig. S8A). Homologs of MbrR-regulated genes, accompanied by conserved MbrR-binding motifs, were consistently identified across most streptococcal species, indicating a conserved defense strategy (Fig. S8B and Table S4). In contrast, the flotillin MbrH and the CaaX protease MbrX appear less conserved (Fig. S8B), suggesting lineage-specific adaptation or functional redundancy. Intriguingly, MbrR-binding motifs were also identified upstream of genes encoding bacteriocin immunity peptides in some species, suggesting that MbrR may coordinate multiple layers of bacteriocin resistance (Table S4). Consistent with this hypothesis, a related YtrA-like regulator has previously been implicated in resistance to host-derived antimicrobial peptides and cell envelope-targeting antibiotics in the pneumococcus [24]. Collectively, these observations support the hypothesis that the MbrRAB system functions as a general AMP-responsive regulatory module across streptococci.

MbrR belongs to the YtrA subfamily within the GntR superfamily of transcriptional regulators [50]. Members of this superfamily are typically modulated by small-molecule ligands that induce conformational changes affecting DNA binding [50]. Although effector molecules have been identified for several GntR subfamilies [51–53], no ligand has yet been conclusively assigned to YtrA regulators. Structural studies suggest that YtrA proteins possess a reduced effector-binding domain, potentially precluding classical ligand-mediated regulation [50, 54, 55]. The founding YtrA regulator was characterized in *Bacillus subtilis*, where it participates in envelope stress responses triggered by cell wall-targeting antibiotics such as vancomycin [56]. However, vancomycin is unlikely to directly interact with cytosolic YtrA, implying the existence of an indirect sensing mechanism. Our structural modeling predicts a direct interaction between YtrA and the ATPase component YtrB of its cognate ABC transporter (Fig. 8), supporting a sequestration-based regulatory model analogous to that described here for MbrR.

**Figure 8.**
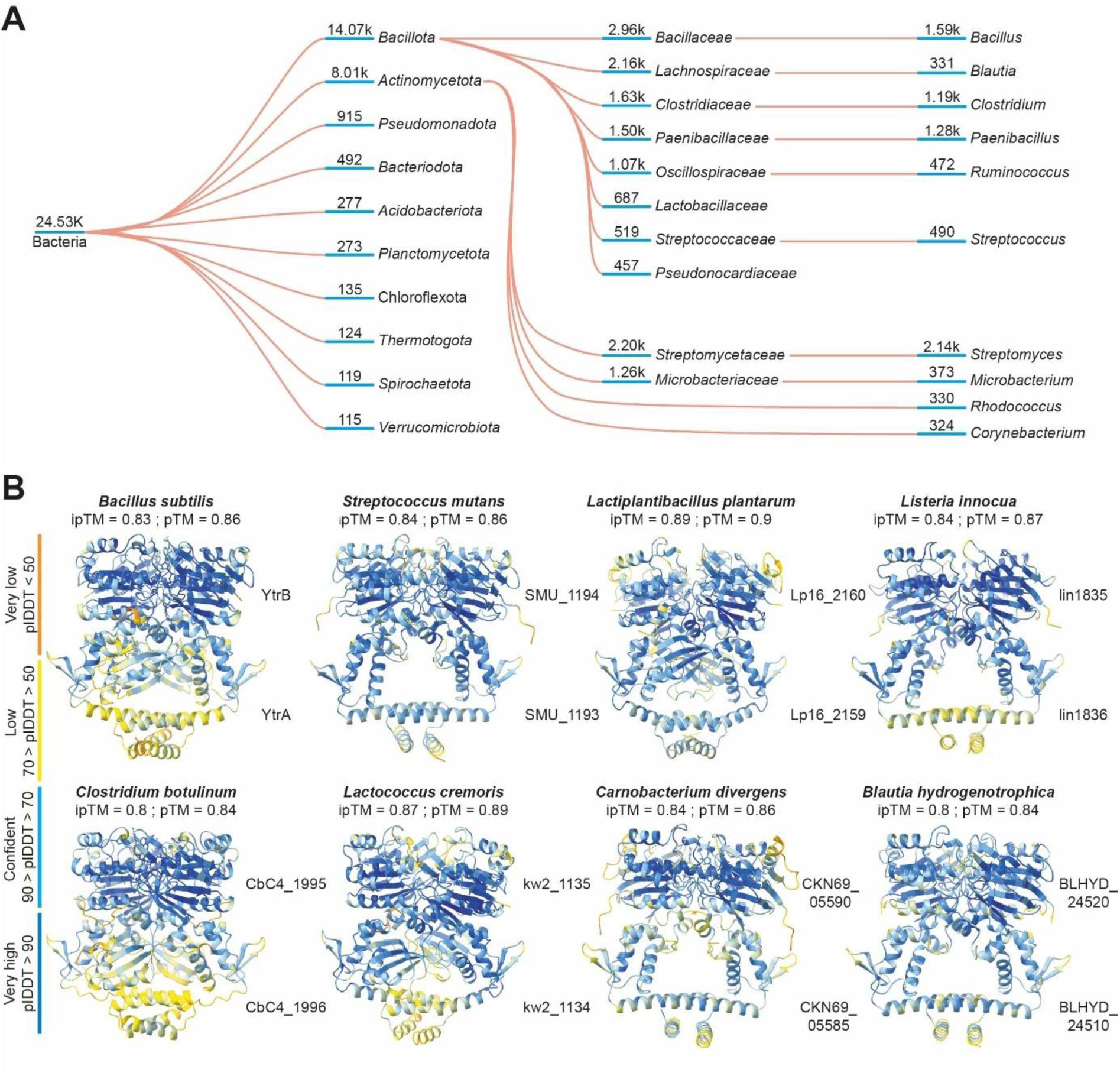
YtrA across bacteria. **A.** Clustering of YtrA in bacteria. Numbers underlined in blue indicate how many YtrA regulators were identified in each phylogenic group. Large-scale structural analysis was performed with the software AFDB clusters. **B.** AlphaFold3 structure predictions of YtrA-like regulators in interaction with their associated ABC transporter ATPase component from various *Bacillota* species. The interface Predicted Template Modeling (iPTM) and Predicted Template Modeling (PTM) scores are indicated on top of each structure. The structure color code is based on the pLDDT confidence score indicated on the left side.

To further explore the conservation of this regulatory paradigm, large-scale structural and genomic analyses suggest that this regulatory principle may be widely conserved. Mining structural databases identified tens of thousands of YtrA-like proteins across bacterial phyla, predominantly within *Bacillota* and *Actinomycetota* (Fig. 8A). Genomic context analyses revealed strong syntenic associations between YtrA-like regulators and adjacent ABC transporter genes (Fig. S9). Furthermore, structural modeling of multiple YtrA-ABC transporter ATPase complexes consistently predicted high-confidence protein-protein interactions (Fig. 8B), supporting the hypothesis that transporter-mediated sequestration represents a general regulatory strategy for YtrA-controlled regulons.

Together, our findings uncover a previously unrecognized mechanism of AMP resistance regulation in Gram-positive bacteria, whereby YtrA-family repressors are controlled through direct interaction with their cognate ABC transporters rather than classical ligand-mediated or post-translational mechanisms. This transporter-mediated sequestration enables rapid and selective transcriptional reprogramming in response to antimicrobial stress, providing an efficient membrane-to-transcription signaling axis. The widespread conservation of this regulatory architecture suggests that it represents a fundamental strategy by which bacteria coordinate resistance responses during interbacterial competition. Importantly, targeting transporter-repressor interactions may offer new opportunities to sensitize pathogens to host-derived or probiotic antimicrobial peptides, highlighting a potential avenue for antimicrobial intervention.

## MATERIAL AND METHODS

### Bacterial strains and DNA material

Bacterial strains, plasmids, synthetic DNA fragments, oligonucleotides, and PCR fragments used in this study are listed in Supplementary Tables 5, 6, 7, 8, and 9, respectively.

### Strain and growth conditions

*Streptococcus salivarius* HSISS4 and its derivatives were grown without shaking at 37°C in M17G (M17 medium with 1% (w/v) glucose; Oxoid) or in CDMG (chemically defined medium with 1% (w/v) glucose) [57]. *Escherichia coli* TOP10 (Invitrogen) was cultivated with shaking at 37°C in Lysogenic Broth (LB) medium. Electrotransformation of *E. coli* was performed as described previously [58]. Solid media inoculated with *S. salivarius* were incubated anaerobically using AnaeroGen packs (Thermo Fisher Scientific). When required, media were supplemented with antibiotics at the following concentrations: ampicillin (250 µg/ml), spectinomycin (200 µg/ml), chloramphenicol (5 µg/ml) or erythromycin (10 µg/ml).

### Strain construction by natural transformation

*S. salivarius* HSISS4 and its derivatives were transformed with Gibson assembly or PCR products for the generation of mutant strains. PCR with the Q5 polymerase (NEB #M0491L) was used to amplify all DNA fragments, following the manufacturer’s recommended protocol. To induce natural transformation, an overnight CDMG preculture was diluted in 500 μl of fresh CDMG at a final OD_600_ of 0.05 and incubated for 135 min at 37°C. Then, 1 µM of XIP and linear DNA were added to the culture, and cells were further incubated for 3 h at 37°C before plating on M17G agar supplemented with appropriate antibiotics. For the back-transformation of SNP/InDel, a co-transformation procedure of two DNA fragments was performed to avoid inserting a resistance cassette close to the mutation. The first donor DNA was a fragment containing the specific mutation. The second DNA acted as a tracer of transformation events as it contains a spectinomycin resistance cassette for insertion at an ectopic locus (*tRNAser*). The transformants were selected on spectinomycin and co-transformation events were screened by PCR. All constructions were verified by DNA sequencing.

### Peptide solubilization

Synthetic pheromone peptide XIP (LPYFAGCL; purity of 95 %) and sBI7 (LPFWLILG; purity of 95%) were resuspended in water at 500 µM and used at a concentration of 1 µM. Synthetic bacteriocins from the PARAGEN collection [35] were provided by Syngulon (Seraing, Belgium) and resuspended in water at 1 mg/ml. Salivaricins not supplied by Syngulon were resuspended in DMSO at 100 µM. All peptides were ordered at Peptide2.0 Inc. (Chantilly, VA, USA).

### Isolation of resistance mutants to sBlpK

To isolate resistant mutants from *S. salivarius* HSISS4 (Δ*blpK–blpI*), 300 µl of an overnight preculture was spread onto M17G agar plates containing a gradient of synthetic BlpK (sBlpK) ranging from 0 to 10 µg/ml. Plates were incubated overnight at 37°C. Colonies growing within the sBlpK gradient were subsequently isolated by streaking onto fresh M17G agar plates. Individual colonies were picked using sterile toothpicks, inoculated into 1 ml of liquid M17G medium, and incubated overnight at 37 °C. Cultures were then frozen at –80°C for long-term preservation.

### Luciferase expression assays

Overnight cultures were diluted in CDMG to reach a final OD_600_ of 0.05. Subsequently, 300 µl of these culture samples were placed in the wells of a sterile covered white microplate with a transparent bottom (Greiner, Alphen a/d Rijn, The Netherlands). When required, cultures were supplemented with synthetic peptides. Growth (OD_600_) and luciferase activity (Relative Light Units, RLU) were measured every 10 minutes for 24 hours using a multi-well plate reader (Hidex Sense, Hidex, Turku, Finland), following the methodology previously described [59].

### Split NanoLuc assays

Overnight cultures were diluted 2:100 in CDMG and incubated at 37°C for 4 hours. Then, cultures were diluted to reach a final OD_600_ of 0.02. Subsequently, 285 µl of these culture samples were placed in the wells of a sterile covered white microplate with a transparent bottom (Greiner) and 15 µl of Nano-Glo Live Cell Substrate (5% of substrate mixed with 95% Nano-Glo LCS Dilution Buffer; Promega, Walldorf, Germany) were added to each well. When required, cultures were supplemented with synthetic peptides. Growth and luciferase activity were measured as reported above.

### Whole Genome Sequencing and data processing

*S. salivarius* HSISS4 (Δ*blpK–blpI*) and its spontaneous resistant mutants were pre-cultured overnight in 2 ml of CDMG medium at 37 °C. Genomic DNA (gDNA) was extracted using the GenElute Bacterial Genomic DNA kit (NA2110, Sigma-Aldrich), following the manufacturer’s instructions. A total of 19 gDNA samples were submitted to Genewiz (Azenta Life Sciences, Leipzig, Germany) for short-read, non-human whole-genome sequencing. FASTQ files were provided for each sample. Sequencing data were uploaded to the Galaxy web platform [60], and genomic variants were identified using the snippy tool by aligning reads to the reference genome.

### RNAseq and data processing

*S. salivarius* HSISS4 (Δ*blpK–blpI*) and its Δ*mbrR* mutant were pre-cultured overnight in CDMG medium at 37°C. Cultures were then diluted to an OD_600_ of 0.05 in 50 ml of fresh CDMG and grown at 37 °C to mid-exponential phase (OD_600_ of 0.6). Cells were harvested by centrifugation at 2,200 × *g* for 5 minutes and resuspended in 6 ml of PBS. One milliliter of this suspension was centrifuged again (2,200 × *g*, 5 minutes), and total RNA was extracted using the RNeasy Mini Kit (QIAGEN), following the manufacturer’s instructions. RNA integrity was assessed using an RNA Nano chip on a Bioanalyzer (Agilent Technologies).

Two biological replicates of each strain were provided to Genewiz (Azenta Life Sciences, Leipzig, Germany) for strand-specific RNA sequencing. The resulting BAM files were analyzed using the RNA-seq quantitation pipeline in SeqMonk (version 1.48.1; Babraham Bioinformatics, UK) to quantify read counts per coding sequence (CDS). Differential expression analysis was performed using the EdgeR statistical filter. Genes were considered differentially expressed if they exhibited an adjusted *P* value < 0.05 and an absolute log₂ fold change ≥ 1.5 (upregulated) or ≤ –1.5 (downregulated).

### Spot-on lawn assays for bacteriocin activity

To assess bacteriocin sensitivity, a two-layer agar assay was performed. A first 30-ml feeding layer of M17G-1.5% agar (w/v) was poured into a Petri dish. Once solidified, a second 15-ml layer of M17G-0.3% agar (w/v) supplemented with 800 µl of an overnight culture of the indicator strain was poured on the top. After solidification, 2 µl of chemically synthesized bacteriocins were spotted onto the surface of the top layer. Plates were incubated overnight at 37°C under anaerobic conditions. Except when specifically indicated, strain HSISS4 Δ*blpK–blpI* was used as indicator strain in all experiments.

### Minimum inhibitory concentration (MIC) determination

MICs were determined using a two-fold serial dilution method across 11 concentrations of the tested antibiotic. Overnight cultures were diluted to a final OD_600_ of 0.05 in M17G medium containing the antibiotic at the defined concentrations. Aliquots of 300 µl were transferred into the wells of a transparent-bottom 96-well microplate (Greiner). Plates were incubated at 37°C for 20 hours. Following incubation, OD_600_ was measured using a multi-well plate reader (Hidex Sense, Hidex, Turku, Finland). MIC values were determined based on three technical replicates per antibiotic concentration.

### MbrR purification

The PCR fragment encoding MbrR-StrepTag (StrepTag at C-ter) was cloned into the pGIR311 vector (pBAD derivative) [61]. This plasmid was electroporated into *E. coli* TOP10. A 25-ml preculture of the recombinant strain was inoculated into 1 L of pre-warmed LB containing ampicillin and incubated at 42°C with continuous shaking. When the culture reached an OD_600_ of ∼ 0.4, it was incubated on ice for 10 minutes. Then, protein expression was induced by adding L-arabinose to a final concentration of 0.1% (w/v). After 4 hours of induction at 30°C with continuous shaking, bacteria were harvested by centrifugation at 2,200 × *g* for 10 minutes, and the pellet was washed in 40 ml of cold buffer W (100 mM Tris pH 8.0, 150 mM NaCl, 1 mM EDTA, pH 7.5). The pellet was then resuspended in 20 ml of cold buffer W supplemented with 1mg/ml of lysozyme and incubated on ice for 30 minutes. The cells were sonicated at 4°C (Vibra cell, Bioblock Scientific), followed by the addition of DNase I (2.5 µg/ml) and MgCl_2_ (1 mM), and further incubation on ice for 30 minutes. The soluble fraction was collected by centrifugation at 23,300 × *g* for 20 min at 4°C. Finally, the recombinant Strep-tagged MbrR protein was then purified on a 1 ml Strep-Tactin XT 4Flow column (IBA BioTAGnology, Göttingen, Germany) according to the manufacturer’s instructions. Elution fractions containing MbrR were pooled and applied to a Superdex Peptide 10/300 GL size-exclusion chromatography column (Cytiva) equilibrated with buffer W.

### Electrophoretic Mobility Shift Assay

Double-stranded DNA fragments (30 bp) were generated by annealing single-stranded oligonucleotides (Table S8), with Cy3 labeling at the 5’ end of one strand and the other strand remaining unlabeled. The binding reaction (20 µl) included 8 µl of binding buffer (20 mM Tris-HCl pH 7.5, 150 mM NaCl, 1 mM EDTA, 1 mM DTT, 10% glycerol, 1 mg/ml BSA), 40 ng of a labeled probe, 1 µg of poly-dIdC, and serial dilutions 2:2 or 1:4 from 2 or 4 µM of purified Strep-Tagged MbrR, respectively. The reaction was incubated at 37°C for 10 minutes before loading the samples onto a native 4–20% gradient gel (iD PAGE Gel; Eurogentec, Belgium). Gel electrophoresis was performed at 120 V for approximately 90 minutes in MOPS buffer (Tris-base 50 mM pH 7.7, MOPS 50 mM, EDTA 1 mM). DNA complexes were visualized on a fluorescent image analyzer (Amersham Typhoon; GE Healthcare Bio-Sciences AB; Sweden).

### Bioinformatic tools

Structure predictions were obtained with AlphaFold3 [36] and corresponding figures were generated using PyMol [62] or ChimeraX [63].

MbrR binding site logo was created by aligning sequences with the MEME suite online software [64].

For phylogenic studies in streptococci, alignment of MbrR sequences was generated using Clustal OMEGA [65]. In parallel, tBLASTn homology searches with default settings were performed on nine selected species of streptococci to identify homologous proteins of the MbrR regulon. Then, upstream promoter regions of identified genes were manually examined to uncover MbrR-binding sites.

Large-scale structural analyses of the YtrA family was performed with software AFDB clusters [66]. The synteny of the *mbrR* locus across various streptococci and members of *Bacillota* was analyzed using the online tool SyntTax [67].

## DATA AVAILABILITY STATEMENT

All data generated or analyzed during this study are included in this published article and supplementary files. All WGS data have been stored in the ENA database under accession number PRJEB106241. RNA-seq data are accessible from the GEO database under accession number GSE318898. The source data underlying the figures and supplementary figures can be found in Supplementary Dataset 1 and 2.

## ACKNOWLEDGMENTS

We warmly thank Philippe Gabant from the Syngulon company who shared synthetic bacteriocins with us from the PARAGEN Collection. The work of PH was supported by the Belgian National Fund for Scientific Research (FNRS, grants PDR T.0110.18/T.0111.22 and CDR J.0090.21), the Concerted Research Actions (ARC, grants 17/22-084 and 22/27-120) from Federation Wallonia-Brussels. JD held a doctoral fellowship from FNRS (FRIA fellowship). LP held a doctoral fellowship from FNRS (FNRS-ASP). JM received funding from the European Union’s Horizon 2020 research and innovation program (Marie Skłodowska-Curie grant N° 101018461). PH is Research Director at the FNRS.

## AUTHOR’S CONTRIBUTION

J.D., L.P., J.M., and P.H.; conception and design. J.D., L.P., C.C.; acquisition of data. J.D., J.M., and P.H.; analysis and interpretation of data. J.D., J.M., and P.H.; draft or revising the manuscript. All authors read and approved the final manuscript.

## SUPPLEMENTAL INFORMATION

Supplemental information includes 9 figures, 9 tables, and 2 datasets.

