## Supplementary material for "A streamlined ABC extruder-repressor module drives multi-bacteriocin resistance in streptococci": Fig. S1, S2, S3, S4, S5, S6, S7, S8, S9 and Table S1, S2, S3, S4, S5, S6, S7, S8, S9

#### 1. SUPPLEMENTAL FIGURES

**Figure S1. Impact of sBlpK on the growth of various simple mutants.**

**Figure S2. MbrR binding site specificity.**

**Figure S3. Comparison of the bacteriocin resistance profile between WT, Δ*mbrR*, and Δ*mbrB*$_{R14P}$ mutant strains.**

**Figure S4. AlphaFold benchmarking of MbrR interaction with MbrAB and MbrCD.**

**Figure S5. Response of P$_{mbrRAB}$-*luxAB* fusion to bacteriocins in wild-type (WT) and Δ*mbrR* backgrounds.**

**Figure S6. Induction of *mbr* genes in response to cereucin X and PsnL.**

**Figure S7. AlphaFold3 structure prediction of dimeric complexes MbrB$_2$-WT and MbrB$_2$-R14P.**

**Figure S8. MbrR distribution and diversity across streptococci.**

**Figure S9. Synteny analysis of YtrA in streptococci and *Bacillota*.**

#### 2. SUPPLEMENTAL TABLES

**Table S1. List of SNPs/InDels detected in spontaneous resistant mutants.**

**Table S2. MIC values for various antibiotics.**

**Table S3: RNA-seq analysis comparing Δ*mbrR* and WT strains.**

**Table S4. MbrR -binding sites identified in various streptococcal species.**

**Table S5. List of bacterial strains used in this study.**

**Table S6. List of plasmids used in this study.**

**Table S7. List of synthetic DNA fragments used in this study.**

**Table S8. List of oligonucleotides used in this study.**

**Table S9. List of PCR fragments used to generate mutants in this study.**

#### 3. SUPPLEMENTAL DATASET

**Dataset S1. All original images used in figures**

**Dataset S2. All numerical data and statistical analyses used in this work**

This file includes Figures S1-S9 and Tables S1-S9.

#### 1. SUPPLEMENTAL FIGURES

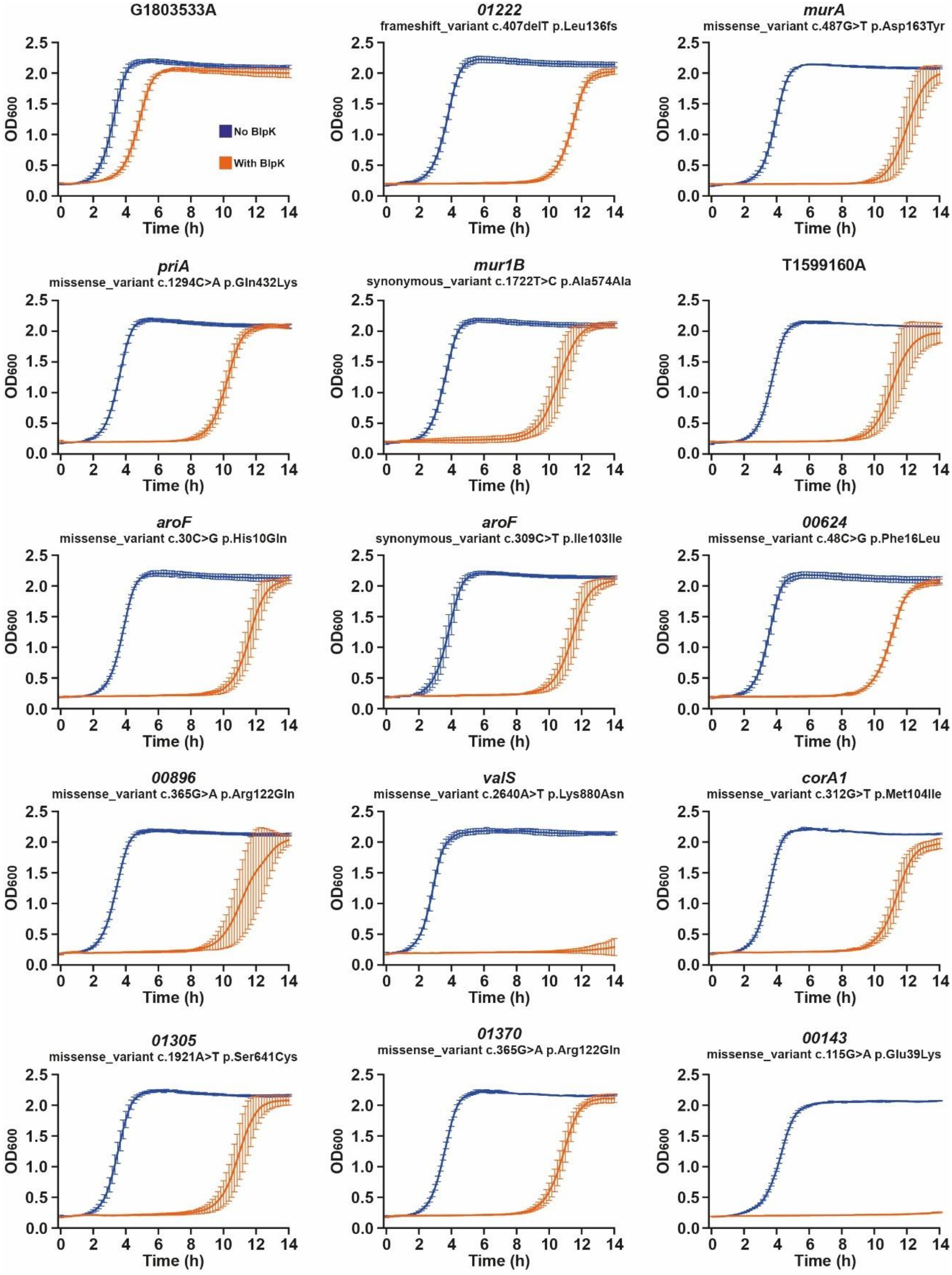

**Figure S1. Impact of sBlpK on the growth of various simple mutants.** Impact of sBlpK on the growth of other SNP/InDel identified by WGS. Growth was monitored in M17G liquid with 2µM of sBlpK (orange) or without (green). Each curve is a mean (± SD) of three independent biological replicates. Each mutation was backcrossed into HSISS4 Δ*blpK–blpI* that was used for the initial selection of spontaneous mutants.

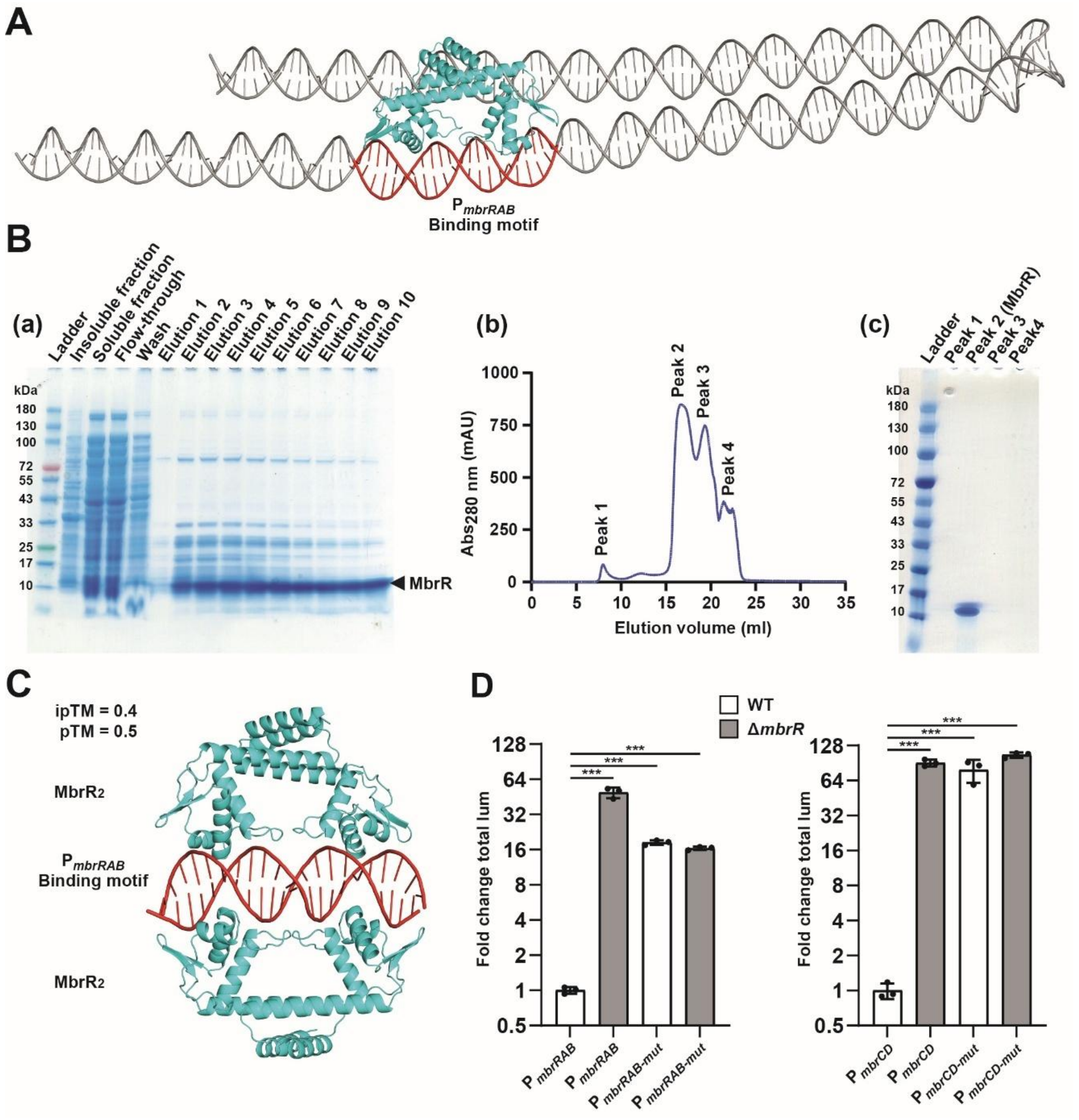

**Figure S2. MbrR binding site specificity. A.** *in silico* binding of MbrR dimer to its operator in $P_{mbrRAB}$. Modelling was performed with AlphaFold3. DNA, binding motif, and MbrR are colored in grey, red, and cyan, respectively. **B**. MbrR purification **(a)** Analysis of purified MbrR by SDS-PAGE prior to size-exclusion chromatography. **(b)** Elution profile of size-exclusion chromatography. **(c)** Analysis of purified MbrR (peak 2) by SDS-PAGE after size-exclusion chromatography. **C.** Structural prediction by AlphaFold3 of interaction between two dimers of MbrR (cyan) and MbrR operator from $P_{mbrRAB}$ (red). Interface Predicted Template Modeling (ipTM) and Predicted Template Modeling (pTM) scores are indicated on the top-left side of the structure. **D.** Regulation of mutated binding sites. Fold change in total luminescence (Fold change total lum) produced by the activation of $P_{mbrRAB}$ (control), $P_{mbrRAB\text{-}mut}$, $P_{mbrCD}$ (control), and $P_{mbrCD\text{-}mut}$ fused to *luxAB* genes in the wild type (WT) and Δ*mbrR* background. Each replicate was compared to the mean of the wild type. Experimental values represent the averages fold change (± SD) of three independent replicates. One-way ANOVA with Dunnett's test were performed (***, $P < 0.001$).

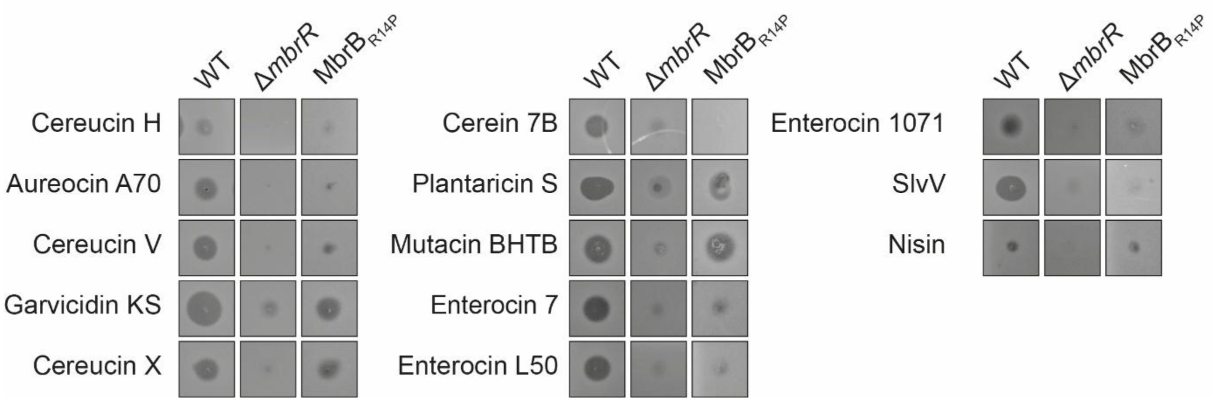

**Figure S3. Comparison of the bacteriocin resistance profile between WT, Δ*mbrR*, and Δ*mbrB*$_{R14P}$ mutant strains.** 2 µl of each synthetic bacteriocin was spotted on plates. For clarity, bacteriocin cocktails composed of multiple peptides are referred to the generic name of the corresponding bacteriocin. Additional details on concentration and solubilization are available in the Methods section.

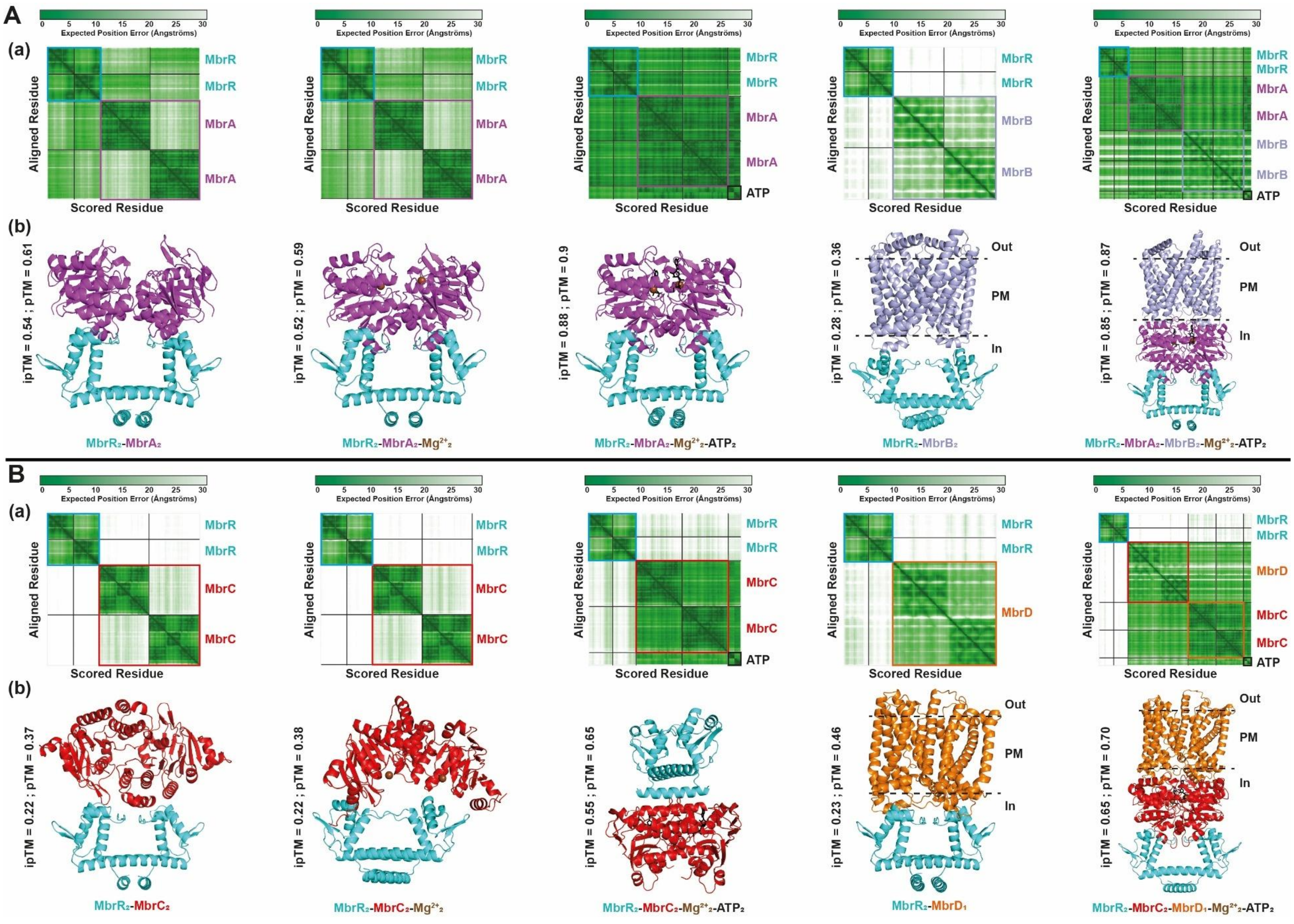
A
(a)
Expected Position Error (Ångströms)
Aligned Residue
Scored Residue
MbrR
MbrA
MbrB
ATP
(b)
ipTM = 0.54 ; pTM = 0.61
ipTM = 0.52 ; pTM = 0.59
ipTM = 0.88 ; pTM = 0.9
ipTM = 0.28 ; pTM = 0.36
ipTM = 0.85 ; pTM = 0.87
Out
PM
In
MbrR₂-MbrA₂
MbrR₂-MbrA₂-Mg²⁺₂
MbrR₂-MbrA₂-Mg²⁺₂-ATP₂
MbrR₂-MbrB₂
MbrR₂-MbrA₂-MbrB₂-Mg²⁺₂-ATP₂
B
(a)
MbrC
MbrD
(b)
ipTM = 0.22 ; pTM = 0.37
ipTM = 0.22 ; pTM = 0.38
ipTM = 0.55 ; pTM = 0.65
ipTM = 0.23 ; pTM = 0.46
ipTM = 0.65 ; pTM = 0.70
MbrR₂-MbrC₂
MbrR₂-MbrC₂-Mg²⁺₂
MbrR₂-MbrC₂-Mg²⁺₂-ATP₂
MbrR₂-MbrD₁
MbrR₂-MbrC₂-MbrD₁-Mg²⁺₂-ATP₂

**Figure S4. AlphaFold benchmarking of MbrR interaction with MbrAB (A) and MbrCD (B).** (**a**) Predicted Aligned Error (PAE) heatmap of AlphaFold-Multimer predictions for the different complexes. (**b**) AlphaFold3 structural prediction. Interface Predicted Template Modeling (ipTM) and Predicted Template Modeling (pTM) scores are indicated to the left side of the structure. Proteins and molecules are color-coded as follows: MbrR (cyan), MbrA (purple), MbrB (light blue), MbrC (red), MbrD (orange), ATP (black), and $Mg^{2+}$ (brown). Subscript numbers refer to the number of proteins or molecules included in the model. The dotted line represents the plasma membrane (PM).

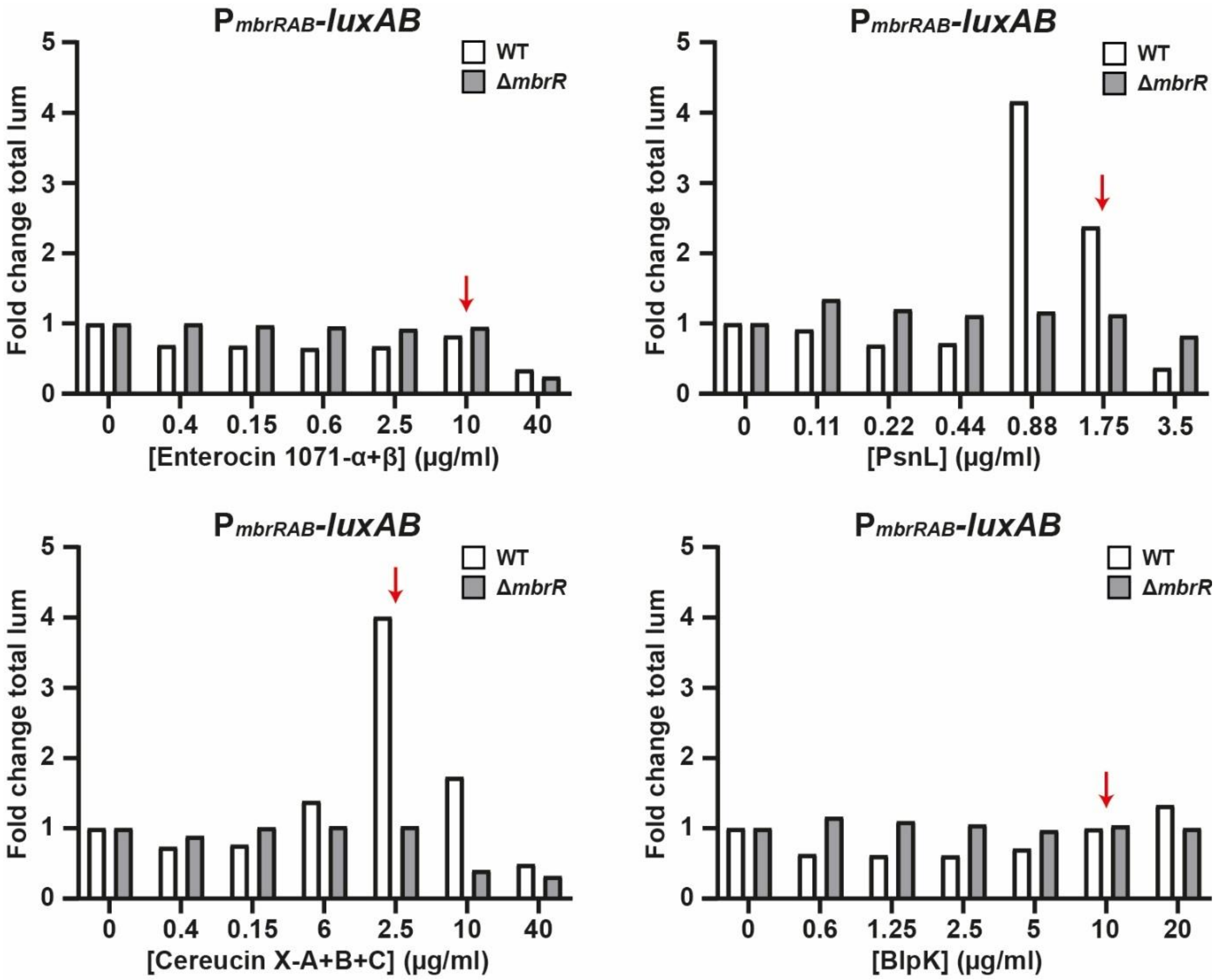

**Figure S5. Response of P*mbrRAB*-*luxAB* fusion to bacteriocins in wild-type (WT) and Δ*mbrR* backgrounds.** Fold change in total luminescence (Fold change total lum) was obtained by comparing each condition to the one without bacteriocin. Red arrow indicates the lowest concentration at which bacteriocin addition impacts growth. This experiment was a prescreen performed without replicates.

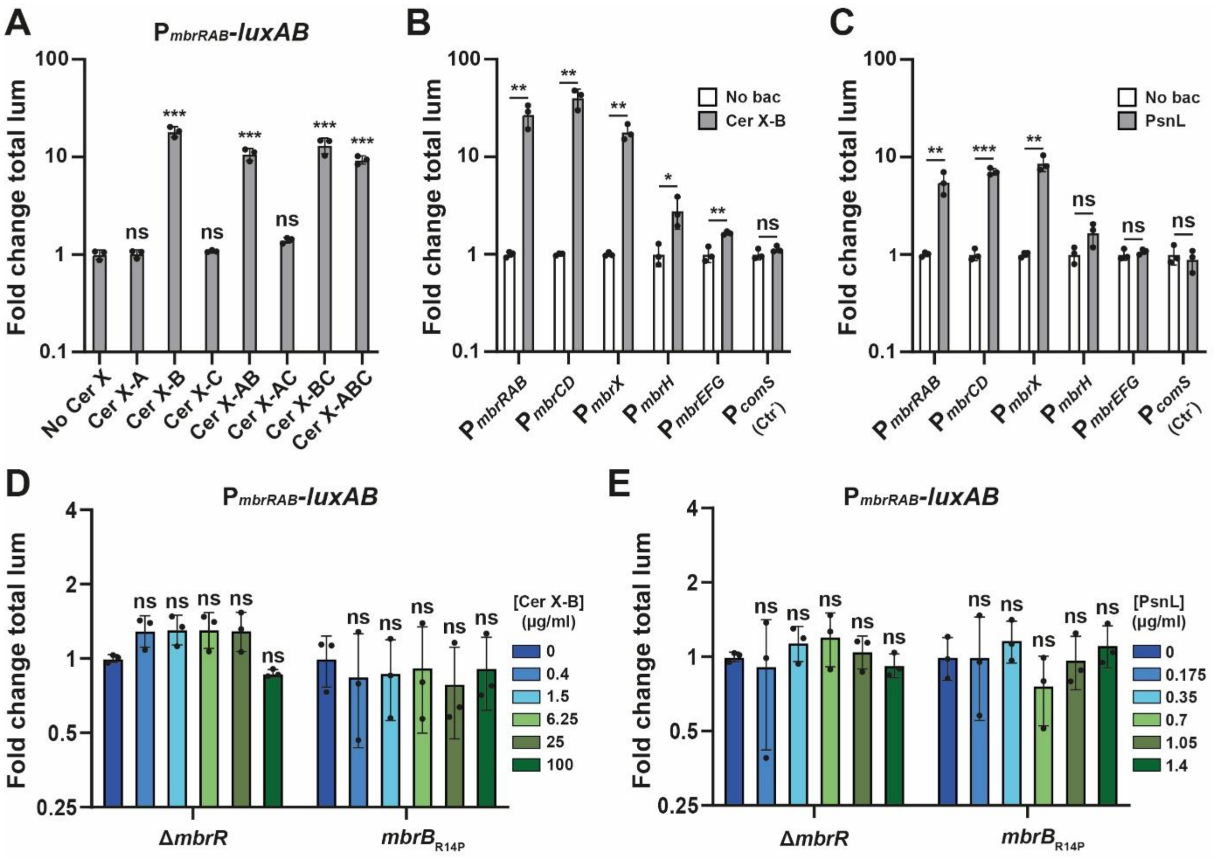

**Figure S6. Induction of *mbr* genes in response to cereucin X and PsnL. A.** $P_{mbrRAB}$ activation in response to cereucin (Cer) X-A, X-B, X-C, and various peptide combinations (AB, AC, BC, and ABC). Fold change in total luminescence (Fold change total lum) produced by the $P_{mbrRAB}$-*luxAB* fusion in WT background with each Cer X peptide alone or combination (10 µg/ml of peptide in all cases). **B.** and **C.** Cer X-B- or PsnL-dependent activation of *mbr* promoters. Fold change in total luminescence produced by each promoter fused to the *luxAB* genes in WT background with 10 µg/ml of Cer X-B (panel B) or 0.7 µg/ml of PsnL (panel C). The promoter of *comS* ($P_{comS}$) was used as negative control ($Ctr^-$). **D.** and **E.** $P_{mbrRAB}$ activation in response to increasing concentration of Cer X-B (panel D) or PsnL (panel E). Fold change in total luminescence produced by the $P_{mbrRAB}$-*luxAB* fusion with 0 to 10 µg/ml of Cer X-B (panel D) or 0 to 1.4 µg/ml of PsnL (panel E) in Δ*mbrR* or $mbrB_{R14P}$ background. For all panels, fold change in total luminescence were obtained by comparing each replicate to the mean of the condition without bacteriocin addition. Experimental values represent the average fold change (± SD) of three independent replicates. One-way ANOVA with Dunnett's test and *t* test were performed for panels A/D/E and panels B/C, respectively (ns, non-significant; *, $P < 0.05$; **, $P < 0.01$; ***, $P < 0.001$).

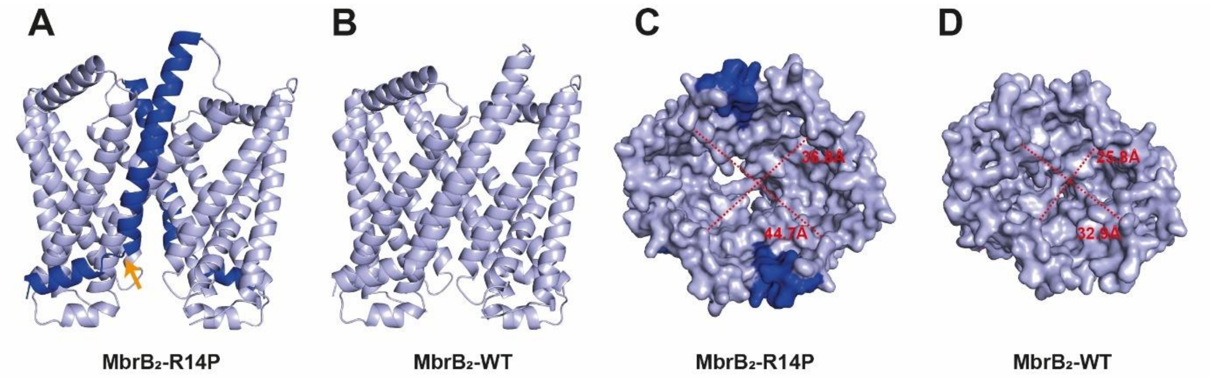

**Figure S7. AlphaFold3 structure prediction of dimeric complexes $MbrB_2$-WT·and $MbrB_2$-R14P.** **A**. and **B**. Lateral view of $MbrB_2$-R14P (panel A) and $MbrB_2$-WT·(panel B) complexes. The first mutated alpha-helix is colored in dark blue in the $MbrB_2$-R14P complex. The orange arrow indicates the position of the R14P mutation. **C**. and **D**. Top view of $MbrB_2$-R14P (panel C) and $MbrB_2$-WT·(panel D) complexes with surface representation. The predicted bacteriocin-binding cavity is around 10 ångström larger at the top in the $MbrB_2$-R14P complex compared to the WT.

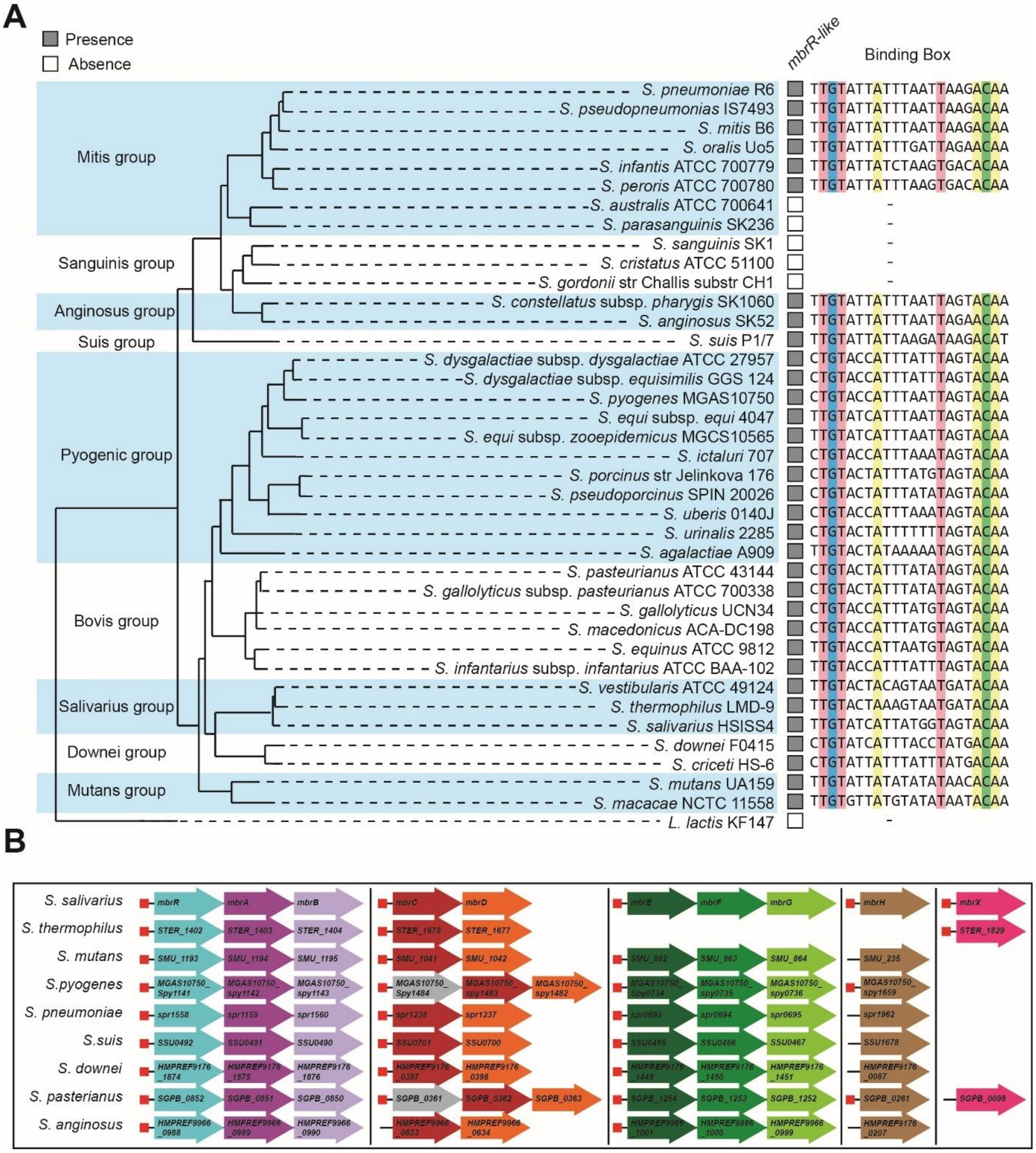

**Figure S8. MbrR distribution and diversity across streptococci. A.** Conservation of the MbrR system across *Streptococcus* species. The phylogenetic tree was adapted from Z. Q. Shao *et al.,* 2013 [1]. A gray box indicates the presence of an *mbrR-like* gene, whereas an empty box denotes the absence of an ortholog in the corresponding genome. Conserved nucleotides within the MbrR binding motif are color-coded by base: red for thymine, blue for guanine, yellow for adenine, and green for cytosine. **B.** Scheme of genomic organization of MbrR regulon in streptococci. Red squares represent MbrR regulation motifs, and arrows represent genes. Arrows are colored according to their putative function, *mbrRAB-like* (blue to purple arrow), *mbrCD-like* (red and orange arrow), *mbrEFG-like* (dark to light green arrow), *mbrH-like* (brown arrow), *mbrX-like* (pink arrow), and bacteriocin immunity gene (gray arrow).

**A.**

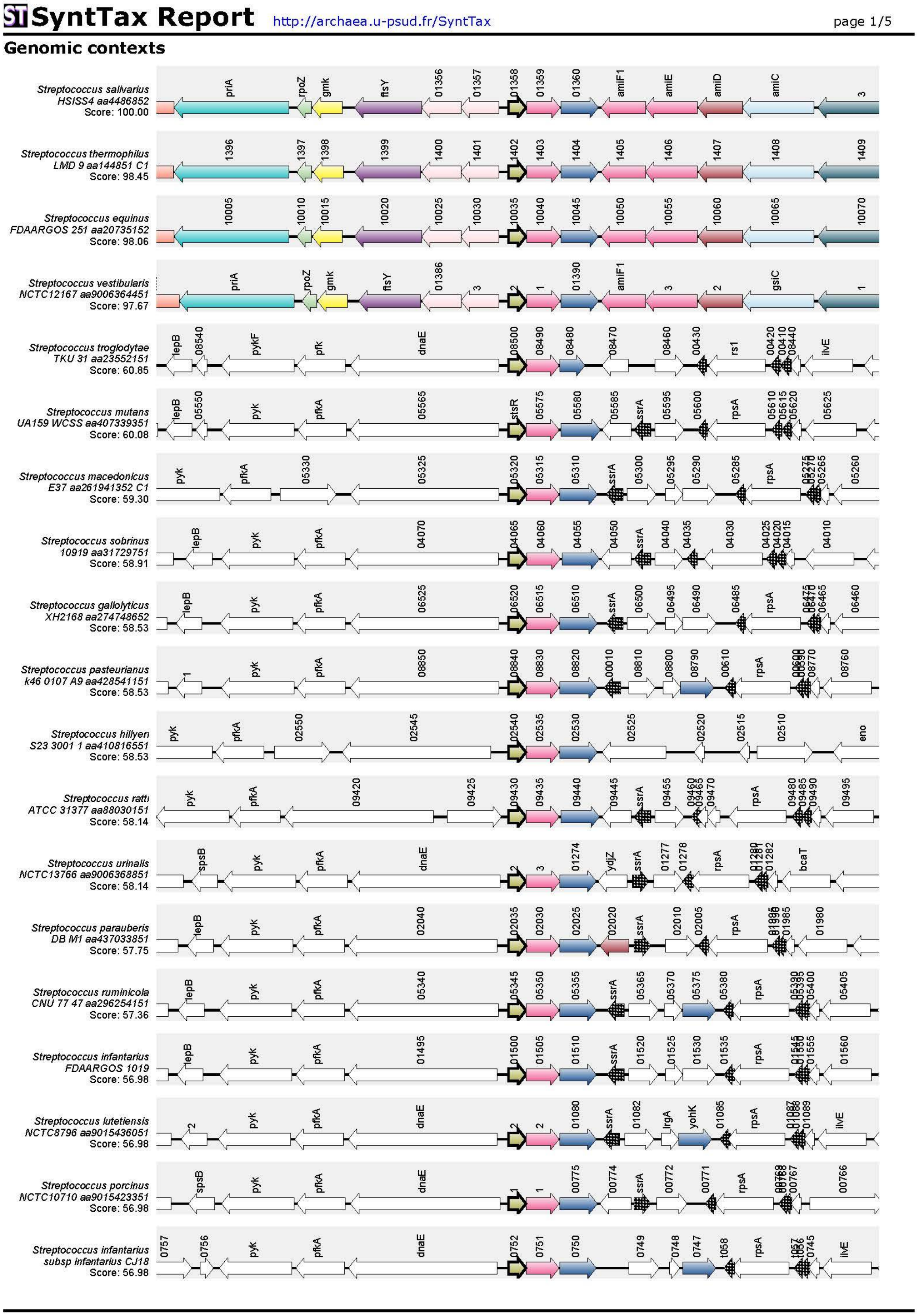
SyntTax Report
http://archaea.u-psud.fr/SyntTax
page 1/5
Genomic contexts
Streptococcus salivarius HSISS4 aa4486852 Score: 100.00
Streptococcus thermophilus LMD 9 aa144851 C1 Score: 98.45
Streptococcus equinus FDAARGOS 251 aa20735152 Score: 98.06
Streptococcus vestibularis NCTC12167 aa9006364451 Score: 97.67
Streptococcus troglodytae TKU 31 aa23552151 Score: 60.85
Streptococcus mutans UA159 WCSS aa407339351 Score: 60.08
Streptococcus macedonicus E37 aa261941352 C1 Score: 59.30
Streptococcus sobrinus 10919 aa31729751 Score: 58.91
Streptococcus gallolyticus XH2168 aa274748652 Score: 58.53
Streptococcus pasteurianus k46 0107 A9 aa428541151 Score: 58.53
Streptococcus hillyeri S23 3001 1 aa410816551 Score: 58.53
Streptococcus ratti ATCC 31377 aa88030151 Score: 58.14
Streptococcus urinalis NCTC13766 aa9006368851 Score: 58.14
Streptococcus parauberis DB M1 aa437033851 Score: 57.75
Streptococcus ruminicola CNU 77 47 aa296254151 Score: 57.36
Streptococcus infantarius FDAARGOS 1019 Score: 56.98
Streptococcus lutetiensis NCTC8796 aa9015436051 Score: 56.98
Streptococcus porcinus NCTC10710 aa9015423351 Score: 56.98
Streptococcus infantarius subsp infantarius CJ18 Score: 56.98

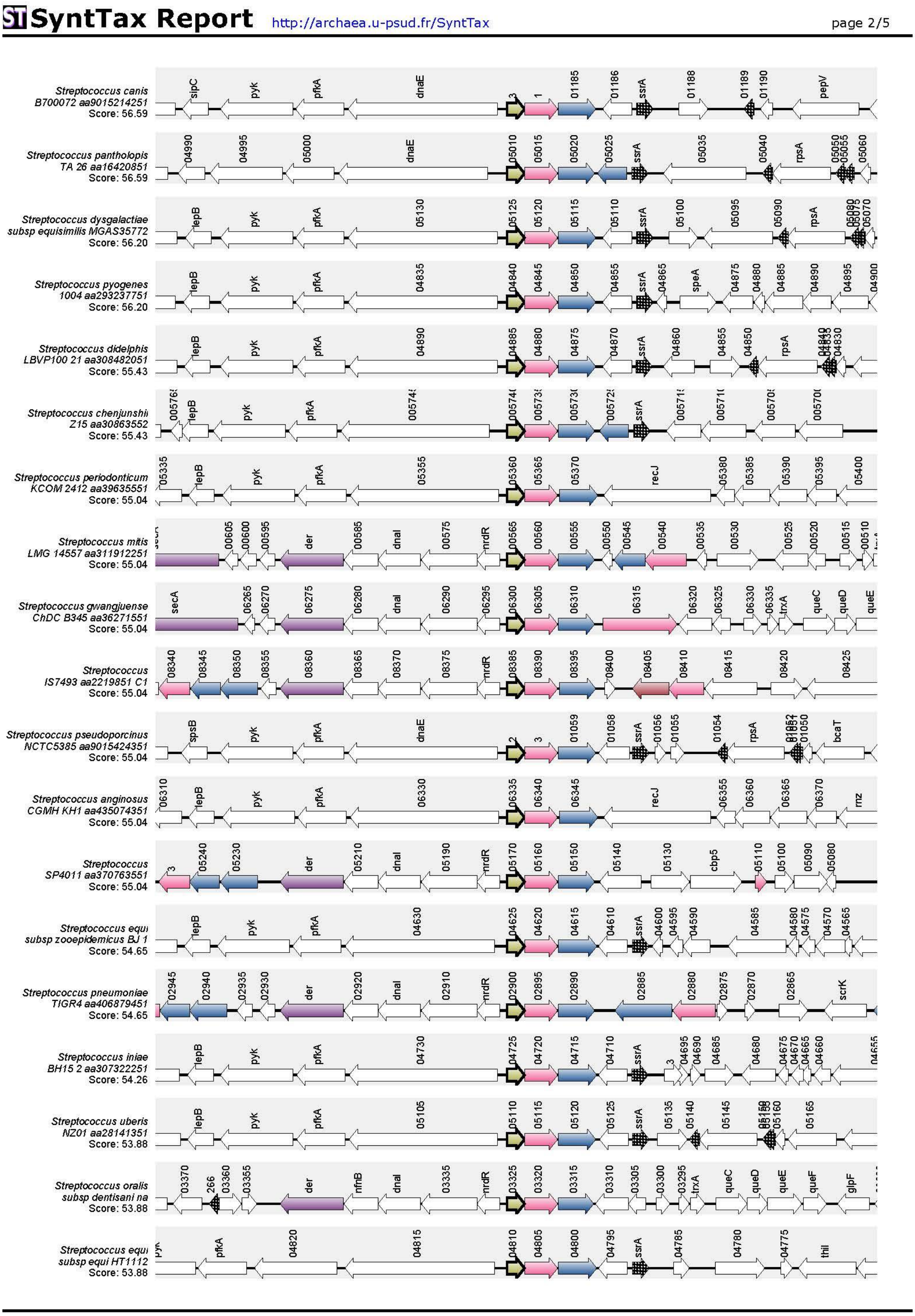
SyntTax Report
http://archaea.u-psud.fr/SyntTax
page 2/5
Streptococcus canis B700072 aa9015214251 Score: 56.59
Streptococcus pantholopis TA 26 aa16420851 Score: 56.59
Streptococcus dysgalactiae subsp equisimilis MGAS35772 Score: 56.20
Streptococcus pyogenes 1004 aa293237751 Score: 56.20
Streptococcus didelphis LBVP100 21 aa308482051 Score: 55.43
Streptococcus chenjunshii Z15 aa30863552 Score: 55.43
Streptococcus periodonticum KCOM 2412 aa39635551 Score: 55.04
Streptococcus mitis LMG 14557 aa311912251 Score: 55.04
Streptococcus gwangjuense ChDC B345 aa36271551 Score: 55.04
Streptococcus IS7493 aa2219851 C1 Score: 55.04
Streptococcus pseudoporcinus NCTC5385 aa9015424351 Score: 55.04
Streptococcus anginosus CGMH KH1 aa435074351 Score: 55.04
Streptococcus SP4011 aa370763551 Score: 55.04
Streptococcus equi subsp zooepidemicus BJ 1 Score: 54.65
Streptococcus pneumoniae TIGR4 aa406879451 Score: 54.65
Streptococcus iniae BH15 2 aa307322251 Score: 54.26
Streptococcus uberis NZ01 aa28141351 Score: 53.88
Streptococcus oralis subsp dentisani na Score: 53.88
Streptococcus equi subsp equi HT1112 Score: 53.88

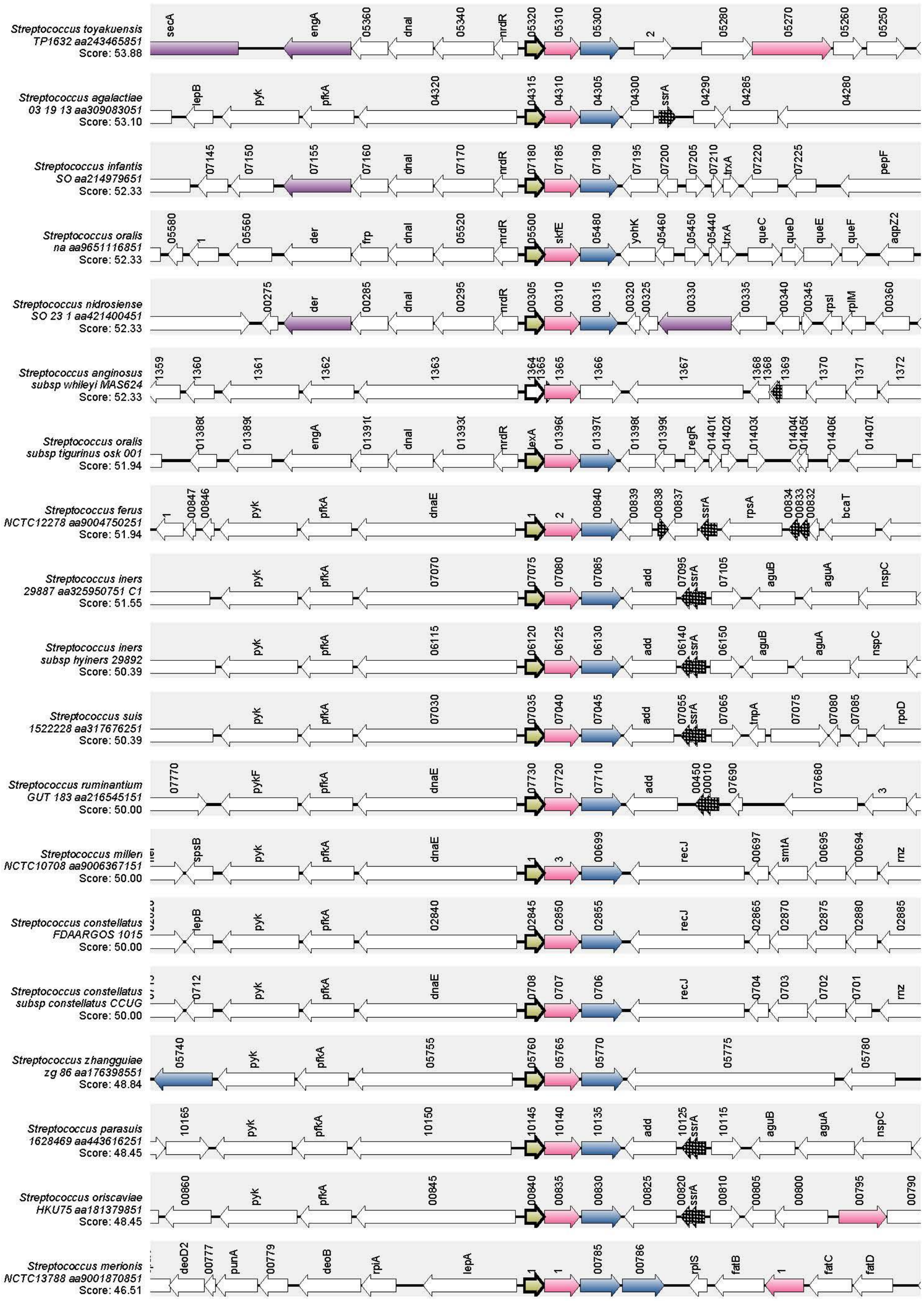
Streptococcus toyakuensis TP1632 aa243465851 Score: 53.88
Streptococcus agalactiae 03 19 13 aa309083051 Score: 53.10
Streptococcus infantis SO aa214979651 Score: 52.33
Streptococcus oralis na aa9651116851 Score: 52.33
Streptococcus nidrosiense SO 23 1 aa421400451 Score: 52.33
Streptococcus anginosus subsp whileyi MAS624 Score: 52.33
Streptococcus oralis subsp tigurinus osk 001 Score: 51.94
Streptococcus ferus NCTC12278 aa9004750251 Score: 51.94
Streptococcus iners 29887 aa325950751 C1 Score: 51.55
Streptococcus iners subsp hyiners 29892 Score: 50.39
Streptococcus suis 1522228 aa317676251 Score: 50.39
Streptococcus ruminantium GUT 183 aa216545151 Score: 50.00
Streptococcus milleri NCTC10708 aa9006367151 Score: 50.00
Streptococcus constellatus FDAARGOS 1015 Score: 50.00
Streptococcus constellatus subsp constellatus CCUG Score: 50.00
Streptococcus zhangguiae zg 86 aa176398551 Score: 48.84
Streptococcus parasuis 1628469 aa443616251 Score: 48.45
Streptococcus oriscaviae HKU75 aa181379851 Score: 48.45
Streptococcus merionis NCTC13788 aa9001870851 Score: 46.51

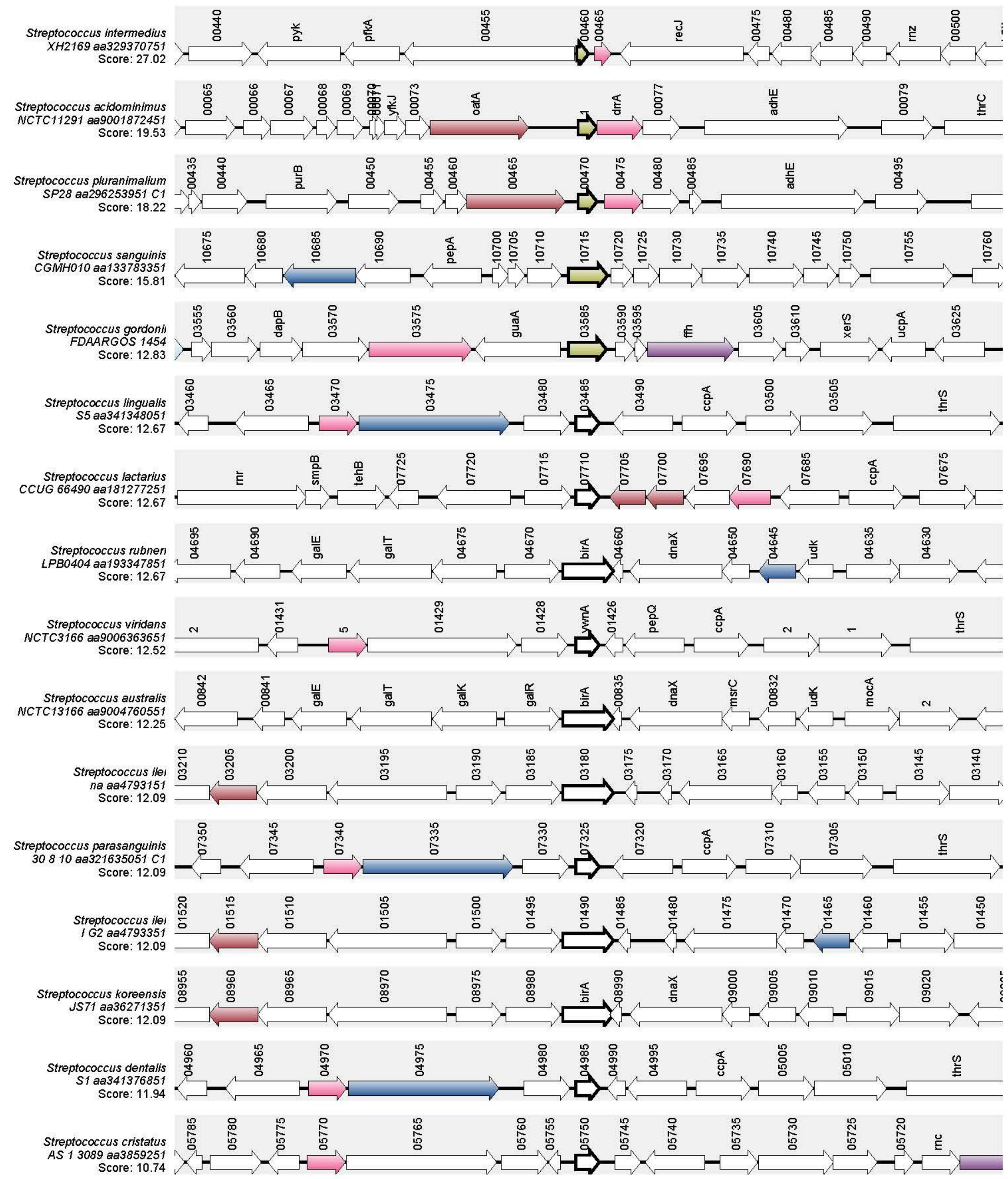
Streptococcus intermedius XH2169 aa329370751 Score: 27.02
Streptococcus acidominimus NCTC11291 aa9001872451 Score: 19.53
Streptococcus pluranimalium SP28 aa296253951 C1 Score: 18.22
Streptococcus sanguinis CGMH010 aa133783351 Score: 15.81
Streptococcus gordonii FDAARGOS 1454 Score: 12.83
Streptococcus lingualis S5 aa341348051 Score: 12.67
Streptococcus lactarius CCUG 66490 aa181277251 Score: 12.67
Streptococcus rubneri LPB0404 aa193347851 Score: 12.67
Streptococcus viridans NCTC3166 aa9006363651 Score: 12.52
Streptococcus australis NCTC13166 aa9004760551 Score: 12.25
Streptococcus ilei na aa4793151 Score: 12.09
Streptococcus parasanguinis 30 8 10 aa321635051 C1 Score: 12.09
Streptococcus ilei I G2 aa4793351 Score: 12.09
Streptococcus koreensis JS71 aa36271351 Score: 12.09
Streptococcus dentalis S1 aa341376851 Score: 11.94
Streptococcus cristatus AS 1 3089 aa3859251 Score: 10.74

**B.**

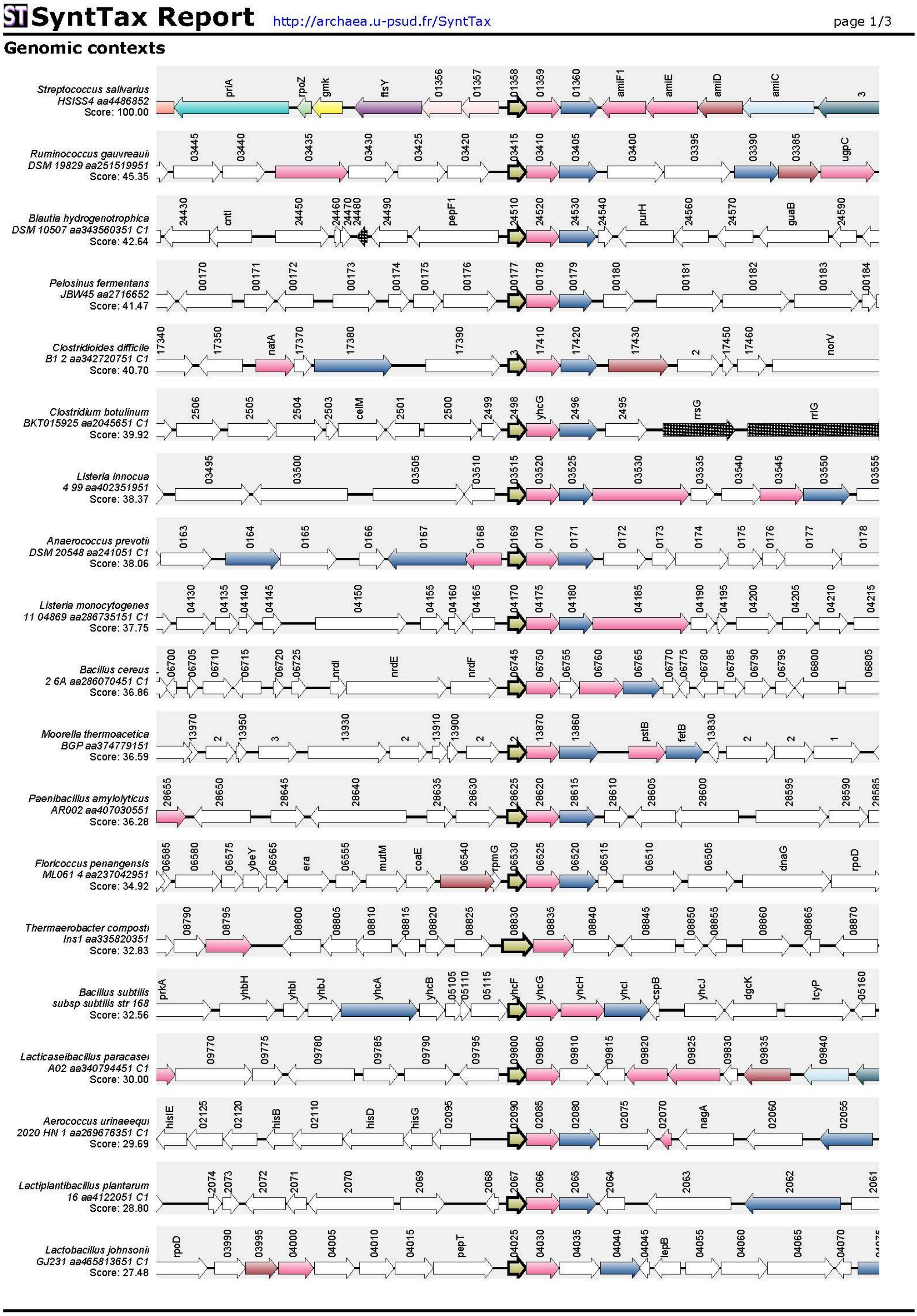
SyntTax Report
http://archaea.u-psud.fr/SyntTax
page 1/3
Genomic contexts
Streptococcus salivarius HSISS4 aa4486852 Score: 100.00
Ruminococcus gauvreauii DSM 19829 aa251519951 Score: 45.35
Blautia hydrogenotrophica DSM 10507 aa343560351 C1 Score: 42.64
Pelosinus fermentans JBW45 aa2716652 Score: 41.47
Clostridioides difficile B1 2 aa342720751 C1 Score: 40.70
Clostridium botulinum BKT015925 aa2045651 C1 Score: 39.92
Listeria innocua 4 99 aa402351951 Score: 38.37
Anaerococcus prevotii DSM 20548 aa241051 C1 Score: 38.06
Listeria monocytogenes 11 04869 aa286735151 C1 Score: 37.75
Bacillus cereus 2 6A aa286070451 C1 Score: 36.86
Moorella thermoacetica BGP aa374779151 Score: 36.59
Paenibacillus amylolyticus AR002 aa407030551 Score: 36.28
Floricoccus penangensis ML061 4 aa237042951 Score: 34.92
Thermaerobacter composti Ins1 aa335820351 Score: 32.83
Bacillus subtilis subsp subtilis str 168 Score: 32.56
Lacticaseibacillus paracasei A02 aa340794451 C1 Score: 30.00
Aerococcus urinaeequi 2020 HN 1 aa269676351 C1 Score: 29.69
Lactiplantibacillus plantarum 16 aa4122051 C1 Score: 28.80
Lactobacillus johnsonii GJ231 aa465813651 C1 Score: 27.48

SyntTax Report http://archaea.u-psud.fr/SyntTax page 2/3

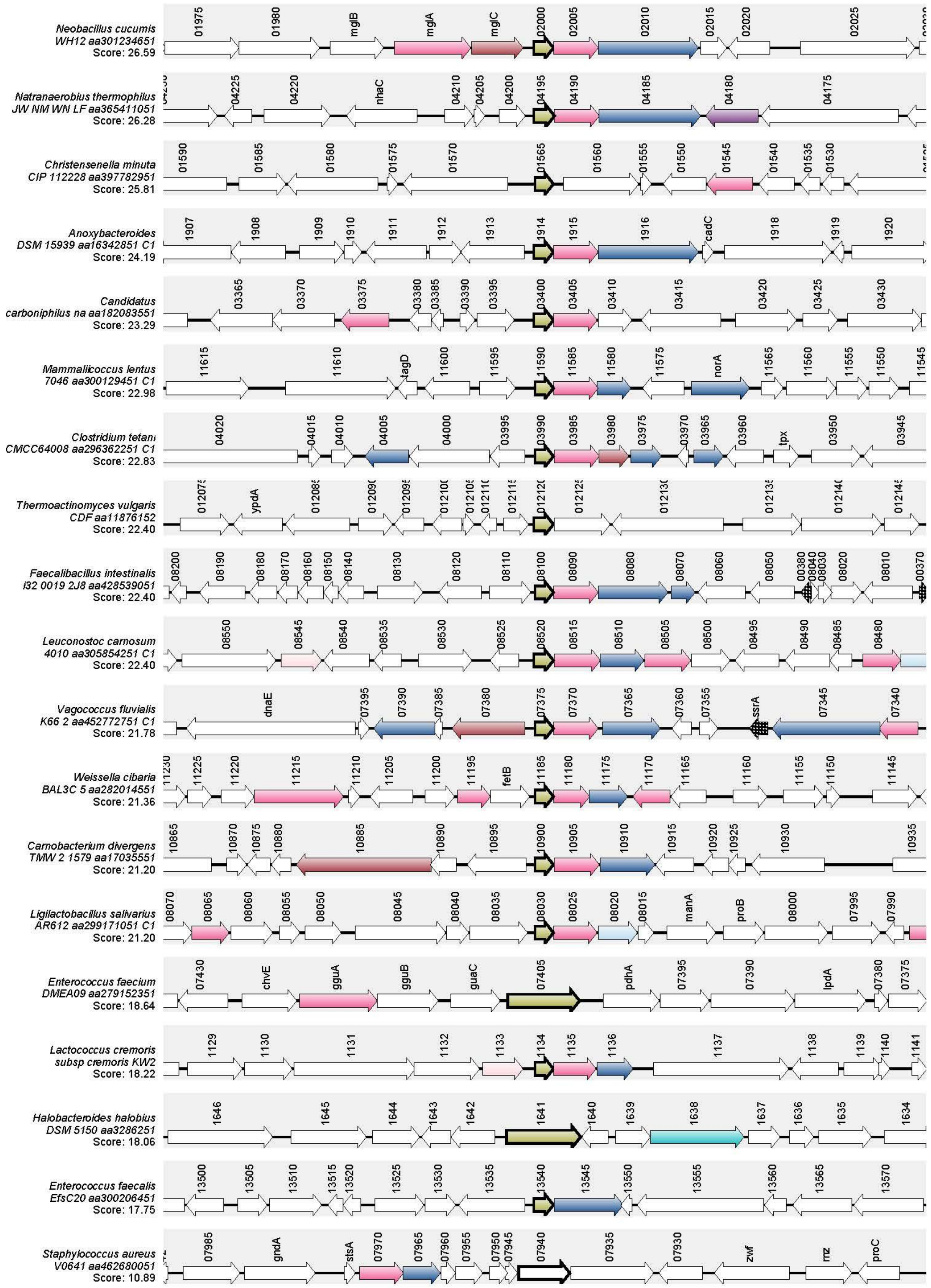

**Figure S9. Synteny analysis of YtrA in streptococci (A) and *Bacillota* (B).** Data were generated with the SyntTax server using MbrR as query.

#### 2. SUPPLEMENTAL TABLES

#### Table S1. List of SNPs/InDels detected in spontaneous resistant mutants.

| N[a] | CHROM[b] | POS[c] | REF[d] | ALT[e] | NT_POS[f] | AA_POS[g] | EFFECT[h] | LOCUS_TAG[i] | PRODUCT[j] |
|---|---|---|---|---|---|---|---|---|---|
| **SNPs/Indels identified by WGS in 19 resistant mutants** | | | | | | | | | |
| 1 | NZ_CP013216 | 1548888 | G | T | 1294/2397 | 432/798 | missense_variant c.1294C>A p.Gln432Lys | HSISS4_01352 (*priA*) | primosomal protein N’ |
| 1 | NZ_CP013216 | 1555851 | G | C | 41/786 | 14/261 | missense_variant c.41G>C p.Arg14Pro | HSISS4_01360 (*mbrB*) | hypothetical protein |
| 2 | NZ_CP013216 | 673331 | C | G | 48/399 | 16/132 | missense_variant c.48C>G p.Phe16Leu | HSISS4_00624 | DUF948 domain-containing protein |
| 2 | NZ_CP013216 | 1555850 | C | CGTA | 44/786 | 15/261 | disruptive_inframe_insertion c.41_43dupGTA p.Arg14_Lys15insSer | HSISS4_01360 (*mbrB*) | hypothetical protein |
| 3 | NZ_CP013216 | 173299 | G | A | 115/816 | 39/271 | missense_variant c.115G>A p.Glu39Lys | HSISS4_00143 | endonuclease/exonuclease/phosphatase family protein |
| 3 | NZ_CP013216 | 1554842 | C | T | 124/378 | 42/125 | stop_gained c.124C>T p.Gln42* | HSISS4_01358 (*mbrR*) | GntR family transcriptional regulator |
| 3 | NZ_CP013216 | 1670328 | C | A | 312/906 | 104/301 | missense_variant c.312G>T p.Met104Ile | HSISS4_01473 (*corA1*) | magnesium transporter CorA family protein |
| 4 | NZ_CP013216 | 1455373 | T | A | 1921/5247 | 641/1748 | missense_variant c.1921A>T p.Ser641Cys | HSISS4_01305 | accessory Sec-dependent serine-rich glycoprotein adhesin |
| 4 | NZ_CP013216 | 1554992 | CT | C | 275/378 | 92/125 | frameshift_variant c.275delT p.Leu92fs | HSISS4_01358 (*mbrR*) | GntR family transcriptional regulator |
| 5 | NZ_CP013216 | 1326901 | CA | C | 407/603 | 136/200 | frameshift_variant c.407delT p.Leu136fs | HSISS4_01222 | nitroreductase family protein |
| 5 | NZ_CP013216 | 1554864 | A | T | 146/378 | 49/125 | missense_variant c.146A>T p.Asn49Ile | HSISS4_01358 (*mbrR*) | GntR family transcriptional regulator |
| 5 | NZ_CP013216 | 1568801 | G | T | 75/675 | 25/224 | synonymous_variant c.75C>A p.Gly25Gly | HSISS4_01370 | GTP pyrophosphokinase family protein |
| 6 | NZ_CP013216 | 928128 | T | C | 1722/2127 | 574/708 | synonymous_variant c.1722T>C p.Ala574Ala | HSISS4_00855 (*mur1B*) | N-acetylmuramoyl-L-alanine amidase |
| 7 | NZ_CP013216 | 1554912 | AT | A | 195/378 | 65/125 | frameshift_variant c.195delT p.Ser66fs | HSISS4_01358 (*mbrR*) | GntR family transcriptional regulator |
| 8 | NZ_CP013216 | 928128 | T | C | 1722/2127 | 574/708 | synonymous_variant c.1722T>C p.Ala574Ala | HSISS4_00855 (*mur1B*) | N-acetylmuramoyl-L-alanine amidase |
| 8 | NZ_CP013216 | 970488 | G | A | 365/609 | 122/202 | missense_variant c.365G>A p.Arg122Gln | HSISS4_00896 | histidine phosphatase family protein |
| 8 | NZ_CP013216 | 1548888 | G | T | 1294/2397 | 432/798 | missense_variant c.1294C>A p.Gln432Lys | HSISS4_01352 (*priA)* | primosomal protein N' |
| 8 | NZ_CP013216 | 1555851 | G | C | 41/786 | 14/261 | missense_variant c.41G>C p.Arg14Pro | HSISS4_01360 | hypothetical protein |
| 9 | NZ_CP013216 | 1548888 | G | T | 1294/2397 | 432/798 | missense_variant c.1294C>A p.Gln432Lys | HSISS4_01352 (*priA)* | primosomal protein N' |
| 9 | NZ_CP013216 | 1555851 | G | C | 41/786 | 14/261 | missense_variant c.41G>C p.Arg14Pro | HSISS4_01360 | hypothetical protein |
| 10 | NZ_CP013216 | 1554992 | CT | C | 275/378 | 92/125 | frameshift_variant c.275delT p.Leu92fs | HSISS4_01358 (*mbrR*) | GntR family transcriptional regulator |
| 11 | NZ_CP013216 | 1554992 | CT | C | 275/378 | 92/125 | frameshift_variant c.275delT p.Leu92fs | HSISS4_01358 (*mbrR*) | GntR family transcriptional regulator |
| 12 | NZ_CP013216 | 1554933 | G | C | 215/378 | 72/125 | missense_variant c.215G>C p.Arg72Pro | HSISS4_01358 (*mbrR*) | GntR family transcriptional regulator |
| 13 | NZ_CP013216 | 1548888 | G | T | 1294/2397 | 432/798 | missense_variant c.1294C>A p.Gln432Lys | HSISS4_01352 (*priA)* | primosomal protein N' |
| 13 | NZ_CP013216 | 1555851 | G | C | 41/786 | 14/261 | missense_variant c.41G>C p.Arg14Pro | HSISS4_01360 | hypothetical protein |
| 14 | NZ_CP013216 | 928128 | T | C | 1722/2127 | 574/708 | synonymous_variant c.1722T>C p.Ala574Ala | HSISS4_00855 (*mur1B*) | N-acetylmuramoyl-L-alanine amidase |
| 14 | NZ_CP013216 | 1599160 | T | A | - | - | - | - | - |
| 14 | NZ_CP013216 | 1803533 | G | T | - | - | - | - | - |
| 14 | NZ_CP013216 | 447098 | A | T | 2640/2652 | 880/883 | missense_variant c.2640A>T p.Lys880Asn | HSISS4_00397 (*valS*) | valine--tRNA ligase |
| 15 | NZ_CP013216 | 615583 | C | T | 309/1167 | 103/388 | synonymous_variant c.309C>T p.Ile103Ile | HSISS4_00558 (*aroF*) | chorismate synthase |
| 15 | NZ_CP013216 | 1803533 | G | A | - | - | - | - | - |
| 16 | NZ_CP013216 | 615304 | C | G | 30/1167 | 10/388 | missense_variant c.30C>G p.His10Gln | HSISS4_00558 (*aroF*) | chorismate synthase |
| 16 | NZ_CP013216 | 897040 | G | T | 487/1272 | 163/423 | missense_variant c.487G>T p.Asp163Tyr | HSISS4_00824 (*murA*) | UDP-N-acetylglucosamine 1-carboxyvinyltransferase |
| 16 | NZ_CP013216 | 1554720 | TG | T | 3/378 | 1/125 | frameshift_variant c.3delG p.Ala2fs | HSISS4_01358 (*mbrR*) | GntR family transcriptional regulator |
| 17 | NZ_CP013216 | 673331 | C | G | 48/399 | 16/132 | missense_variant c.48C>G p.Phe16Leu | HSISS4_00624 | DUF948 domain-containing protein |
| 17 | NZ_CP013216 | 928128 | T | C | 1722/2127 | 574/708 | synonymous_variant c.1722T>C p.Ala574Ala | HSISS4_00855 (*mur1B*) | N-acetylmuramoyl-L-alanine amidase |
| 17 | NZ_CP013216 | 1555850 | C | CGTA | 44/786 | 15/261 | disruptive_inframe_insertion c.41_43dupGTA p.Arg14_Lys15insSer | HSISS4_01360 | hypothetical protein |
| 18 | NZ_CP013216 | 1599160 | T | A | - | - | - | - | - |
| 18 | NZ_CP013216 | 1803533 | G | T | - | - | - | - | - |
| 19 | NZ_CP013216 | 1803533 | G | A | - | - | - | - | - |
| **SNPs/Indels identified by *mbrR* sequencing in 21 resistant mutants** | | | | | | | | | |
| 20 | NZ_CP013216 | 1554720 | TG | T | 3/378 | 1/125 | frameshift_variant c.3delG p.Ala2fs | HSISS4_01358 (*mbrR*) | GntR family transcriptional regulator |
| 21 | NZ_CP013216 | 1554780 | TG | T | 63/378 | 21/125 | frameshift_variant c.63TG>T p.Lys22fs | HSISS4_01358 (*mbrR*) | GntR family transcriptional regulator |
| 22 | NZ_CP013216 | 1554780 | TG | T | 63/378 | 21/125 | frameshift_variant c.63TG>T p.Lys22fs | HSISS4_01358 (*mbrR*) | GntR family transcriptional regulator |

| | | | | | | | | | |
|---|---|---|---|---|---|---|---|---|---|
| 23 | NZ_CP013216 | 1554792 | C | T | 74/378 | 25/125 | missense_variant c.74C>T p.Ser25Phe | HSISS4_01358 (*mbrR*) | GntR family transcriptional regulator |
| 24 | NZ_CP013216 | 1554815 | C | T | 97/378 | 33/125 | stop_gained c.97C>T p.Gln33* | HSISS4_01358 (*mbrR*) | GntR family transcriptional regulator |
| 25 | NZ_CP013216 | 1554815 | C | T | 97/378 | 33/125 | stop_gained c.97C>T p.Gln33* | HSISS4_01358 (*mbrR*) | GntR family transcriptional regulator |
| 26 | NZ_CP013216 | 1554842 | C | T | 124/378 | 42/125 | stop_gained c.124C>T p.Gln42* | HSISS4_01358 (*mbrR*) | GntR family transcriptional regulator |
| 27 | NZ_CP013216 | 1554864 | A | T | 146/378 | 49/125 | missense_variant c.146A>T p.Asn49Ile | HSISS4_01358 (*mbrR*) | GntR family transcriptional regulator |
| 28 | NZ_CP013216 | 1554879 | C | T | 161/378 | 54/125 | missense_variant c.161C>T p.Ala54Val | HSISS4_01358 (*mbrR*) | GntR family transcriptional regulator |
| 29 | NZ_CP013216 | 1554893 | G | T | 175/378 | 59/125 | stop_gained c.175G>T p.Glu59* | HSISS4_01358 (*mbrR*) | GntR family transcriptional regulator |
| 30 | NZ_CP013216 | 1554912 | AT | A | 195/378 | 65/125 | frameshift_variant c.195delT p.Ser66fs | HSISS4_01358 (*mbrR*) | GntR family transcriptional regulator |
| 31 | NZ_CP013216 | 1554924 | C | T | 206/378 | 69/125 | missense_variant c.206C>T p.Thr69Ile | HSISS4_01358 (*mbrR*) | GntR family transcriptional regulator |
| 32 | NZ_CP013216 | 1554971 | C | T | 253/378 | 85/125 | stop_gained c.253C>T p.Arg85* | HSISS4_01358 (*mbrR*) | GntR family transcriptional regulator |
| 33 | NZ_CP013216 | 1554992 | CT | C | 275/378 | 92/125 | frameshift_variant c.275delT p.Leu92fs | HSISS4_01358 (*mbrR*) | GntR family transcriptional regulator |
| 34 | NZ_CP013216 | 1554992 | CT | C | 275/378 | 92/125 | frameshift_variant c.275delT p.Leu92fs | HSISS4_01358 (*mbrR*) | GntR family transcriptional regulator |
| 35 | NZ_CP013216 | 1554992 | CT | C | 275/378 | 92/125 | frameshift_variant c.275delT p.Leu92fs | HSISS4_01358 (*mbrR*) | GntR family transcriptional regulator |
| 36 | NZ_CP013216 | 1554992 | CT | C | 275/378 | 92/125 | frameshift_variant c.275delT p.Leu92fs | HSISS4_01358 (*mbrR*) | GntR family transcriptional regulator |
| 37 | NZ_CP013216 | 1554992 | CT | C | 275/378 | 92/125 | frameshift_variant c.275delT p.Leu92fs | HSISS4_01358 (*mbrR*) | GntR family transcriptional regulator |
| 38 | NZ_CP013216 | 1554992 | CT | C | 275/378 | 92/125 | frameshift_variant c.275delT p.Leu92fs | HSISS4_01358 (*mbrR*) | GntR family transcriptional regulator |
| 39 | NZ_CP013216 | 1554992 | CT | C | 275/378 | 92/125 | frameshift_variant c.275delT p.Leu92fs | HSISS4_01358 (*mbrR*) | GntR family transcriptional regulator |
| 40 | NZ_CP013216 | 1554992 | CT | C | 275/378 | 92/125 | frameshift_variant c.275delT p.Leu92fs | HSISS4_01358 (*mbrR*) | GntR family transcriptional regulator |

[a] N, spontaneous mutant number
[b] CHROM, reference genome used
[c] POS, position of the mutation in the genome
[d] REF, nucleotide(s) identified in the reference genome
[e] ALT, mutated nucleotide(s) identified in the spontaneous mutant
[f] NT_POS, position of the modified nucleotide in the gene
[g] AA_POS, position of the modified amino acid in the encoded protein
[h] EFFECT, nucleotide modification regarding position in CDS and protein
[i] LOCUS TAG, name of the gene
[j] PRODUCT, annotated function.

#### Table S2. MIC values for various antibiotics.

| Class | Target | Antibiotic | WT | Δ*mbrR* |
|---|---|---|---|---|
| | | | MIC (μg/ml)[a] | |
| Macrolide | 50S | Erythromycin | 2 | 2 |
| Glycopeptide & lipoglycopeptide | Cell wall synthesis | Vancomycin | 1 | 1 |
| Glycopeptide & lipoglycopeptide | Disrupts bacterial cell membrane | Daptomycin | 128 | 128 |
| Glycopeptide & lipoglycopeptide | Cell wall synthesis | Ramoplanin | 0.5 | 0.5 |
| Penicillin | Cell wall synthesis | Penicillin G | 0.12 | 0.12 |
| Penicillin | Cell wall synthesis | Ampicillin | 1 | 1 |
| Cephalosporin | Cell wall synthesis | Cefotaxime | 0.03 | 0.03 |
| Chloramphenicol | 50S | Chloramphenicol | 2 | 2 |
| Tetracycline | 30S | Tetracycline | 0.5 | 0.5 |
| Aminoglycoside | 30S | Streptomycin | 512 | 512 |
| Polypeptide | Cell wall synthesis | Bacitracin | 8 | 8 |
| Phosphonic | Cell wall synthesis | Fosfomycin | 256 | 256 |
| Carboxylic acid | Inhibition protein synthesis | Mupirocin | 0.06 | 0.06 |

[a] Mean values of MIC were determined from three technical replicates performed in the same conditions.

**Table S3. RNA-seq analysis comparing Δ*mbrR* and WT strains.**

| Locus tag (HSISS4_x) | Product | Start | End | *P* value | False Discovery Rate | Log2 Fold Change[a] |
|---|---|---|---|---|---|---|
| 01608 | ABC transporter, ATP-binding protein | 1802643 | 1803374 | 1.6E-16 | 1.3E-13 | 5.04 |
| 01607 | ABC transporter permease protein | 1801051 | 1802640 | 5.1E-17 | 8.3E-14 | 4.99 |
| 01755 | Membrane-bound protease, CAAX family | 1957526 | 1958188 | 5.3E-12 | 2.9E-09 | 4.08 |
| 00448 | ABC transporter permease protein | 503293 | 504501 | 6.8E-07 | 2.8E-04 | 2.86 |
| 00447 | ABC transporter ATP-binding protein | 502597 | 503283 | 1.5E-06 | 5.0E-04 | 2.60 |
| 00446 | Periplasmic component of efflux system | 501333 | 502586 | 1.5E-05 | 3.3E-03 | 2.54 |
| r00078 | 5S ribosomal RNA | 1908627 | 1908742 | 1.0E+00 | 1.0E+00 | 2.38 |
| 01360 | ABC transporter, ATP-binding protein | 1555811 | 1556596 | 1.6E-05 | 3.3E-03 | 2.37 |
| r00043 | tRNA-Asn | 127033 | 127106 | 1.0E+00 | 1.0E+00 | 2.29 |
| *tnp657A* | Mobile element protein | 528780 | 529253 | 1.0E+00 | 1.0E+00 | 2.29 |
| 01359 | ABC-type multidrug transport system, ATPase component | 1555101 | 1555799 | 3.4E-05 | 6.2E-03 | 2.29 |
| 00528 | hypothetical protein | 584990 | 585130 | 3.2E-01 | 1.0E+00 | 2.06 |
| 01430 | Putative stomatin/prohibitin-family membrane protease subunit YbbK | 1624618 | 1625517 | 2.1E-04 | 3.5E-02 | 2.02 |
| r00025 | 5S ribosomal RNA | 68768 | 68883 | 4.7E-01 | 1.0E+00 | 1.80 |
| r00064 | tRNA-Asn | 1902745 | 1902818 | 3.1E-01 | 1.0E+00 | 1.72 |
| r00031 | tRNA-Met | 69330 | 69403 | 5.8E-02 | 1.0E+00 | 1.56 |
| 01400 | Amino acid ABC transporter. amino acid-binding/permease protein | 1597234 | 1598046 | 4.1E-02 | 1.0E+00 | 1.56 |
| 00462 | hypothetical protein | 517145 | 517828 | 1.9E-02 | 1.0E+00 | 1.55 |
| r00036 | tRNA-Gln | 69743 | 69814 | 3.7E-02 | 1.0E+00 | 1.52 |
| r00037 | tRNA-Leu | 69824 | 69907 | 5.1E-02 | 1.0E+00 | 1.51 |
| r00068 | 16S ribosomal RNA | 1906196 | 1907743 | 2.7E-01 | 1.0E+00 | -1.80 |
| *tnpSth1A* | Mobile element protein | 422562 | 424201 | 4.8E-01 | 1.0E+00 | -1.97 |
| r00054 | tRNA-Thr | 462761 | 462833 | 1.0E+00 | 1.0E+00 | -2.28 |
| *tnpSth1G* | ISSth1, transposase (orf1), IS3 family | 1354071 | 1355710 | 1.0E+00 | 1.0E+00 | -2.28 |
| 01358 | Transcriptional regulator, GntR family | 1554719 | 1555096 | 2.0E-06 | 5.6E-04 | -4.89 |

[a] Genes showing a $\log_2$ fold change ≥ 1.5 (upregulated, red) or ≤ –1.5 (downregulated, green)

#### Table S4. MbrR -binding sites identified in various streptococcal species.

| Species | Strain | Putative regulation box | Gene regulated by MbrR |
|---|---|---|---|
| *S. salivarius* | HSISS4 | TTGTTTTATTTATGTGATACAA | *HSISS4_00446* to *00448* |
| | | TTGTATCATTATGGTAGTACAA | *HSISS4_01358* to *01360* |
| | | TTGTATCATTTCAAAGGTACAA | *HSISS4_01608* & *01607* |
| | | TTGTATCTTTATTATAATACAA | *HSISS4_01755* |
| | | TTGTATCACTGTGCTATAATAA | *HSISS4_01430* |
| *S. thermophilus* | LMD9 | TTGTATCATTACTTTAGTACAA | *STER_1402* to *1404* |
| | | TTGTATCATTTCAAAGGTACAA | *STER_1678* & *1677* |
| | | TTGTACTTTTATTATAATACAA | *STER_1829* |
| *S. pneumoniae* | R6 | GTGTATTATATATCTAGTACAA | *spr0693* to *0695* |
| | | TTGTATTATTTAATTAAGACAA | *spr1558* to *1560* |
| | | TTGTATCAAAGTGCTAGTATAA | *spr1238* & *1237* |
| *S. mutans* | UA159 | TTGTATTATAGCCTTAAGACAA | *SMU_862* to *864* |
| | | TTGTATGATTTCAGTAGTATAA | *SMU_1041* & *1042* |
| | | TTGTATTATATATATAACACAA | *SMU_1193* to *1195* |
| *S. suis* | P1/7 | TTGTATTATTAAGATAAGACAT | *SSU0492* to *0490* |
| | | TTATACTGCTGAATTAGTACAA | *SSU0701* & *0700* |
| | | TTGTGTTGTCTATATAATACAC | *SSU0465* to *0467* |
| *S. pyogenes* | MGAS17050 | CTGTACCATTTAATTAGTACAA | *MGAS10750_Spy1141* to *1143* |
| | | TTGTACTATTTGTATAAGACAA | *MGAS10750_Spy0734* to *0736* |
| | | TTGTACTTTTACGCTAATACAA | *MGAS10750_Spy1484* to *1482* |
| | | TTGACTTATTGTTATAATACAA | *MGAS10750_spy1659* |
| *S. pasteurianus* | ATCC 43144 | TTGTATCACTTTAATAGTACAA | *SGPB_0361* to *0363* |
| | | CTGTACTATTTATATAGTACAA | *SGPB_0852* to *0850* |
| | | TTGTACCCATCAATTAATACAA | *SGPB_0261* |
| | | TTGTATTGTTGCGATAAGACAA | *SGPB_1254* to *1252* |
| *S. anginosus* | SK52 | TTGTATTATTTAATTAGAACAA | *HMPREF9966_0988* to *0990* |
| | | TTGTATTATATATTTAACACAA | *HMPREF9966_1001* to *0999* |
| *S. downei* | F0415 | CTATACTATGAAAGTAATACAA | *HMPREF9176_1449* to *1451* |
| | | CTGTATTATTTATTTATGACAA | *HMPREF9176_1874* to *1876* |
| | | TTGTATTAGTTTTACGTAACAA | *HMPREF9176_0397* & *0398* |

#### Table S5. List of bacterial strains used in this study.

| Names | Characteristics | Reference/source |
|---|---|---|
| *Streptococcus salivarius* | | |
| HSISS4 | Wild-type gastro-intestinal tract isolate | [2] |
| JM1006 | HSISS4 Δ*blpK-I::lox72* | [3] |
| JD0026 | JM1006 Δ*mbrR::spec* | This work |
| JD0027 | JM1006 *mbrB*$_{R14P}$, *tRNA*$_{ser}$*::spec* | This work |
| JD0028 | JM1006 *mbrB*$_{insS15}$, *tRNA*$_{ser}$*::spec* | This work |
| JD0029 | JM1006 G1803533A, *tRNA*$_{ser}$*::spec* | This work |
| JD0030 | JM1006 *priA* missense_variant c.1294C>A p.Gln432Lys, *tRNA*$_{ser}$*::spec* | This work |
| JD0031 | JM1006 *mur1B* synonymous_variant c.1722T>C p.Ala574Ala, *tRNA*$_{ser}$*::spec* | This work |
| JD0032 | JM1006 T1599160A, *tRNA*$_{ser}$*::spec* | This work |
| JD0033 | JM1006 *aroF* missense_variant c.30C>G p.His10Gln, *tRNA*$_{ser}$*::spec* | This work |
| JD0034 | JM1006 *aroF* synonymous_variant c.309C>T p.Ile103Ile, *tRNA*$_{ser}$*::spec* | This work |
| JD0035 | JM1006 *00624* missense_variant c.48C>G p.Phe16Leu, *tRNA*$_{ser}$*::spec* | This work |
| JD0036 | JM1006 *00896* missense_variant c.365G>A p.Arg122Gln, *tRNA*$_{ser}$*::spec* | This work |
| JD0037 | JM1006 *valS* missense_variant c.2640A>T p.Lys880Asn, *tRNA*$_{ser}$*::spec* | This work |
| JD0038 | JM1006 *corA1* missense_variant c.312G>T p.Met104Ile, *tRNA*$_{ser}$*::spec* | This work |
| JD0039 | JM1006 *00143* missense_variant c.115G>A p.Glu39Lys, *tRNA*$_{ser}$*::spec* | This work |
| JD0040 | JM1006 *01305* missense_variant c.1921A>T p.Ser641Cys, *tRNA*$_{ser}$*::spec* | This work |
| JD0041 | JM1006 *01370* synonymous_variant c.75C>A p.Gly25Gly, *tRNA*$_{ser}$*::spec* | This work |
| JD0042 | JM1006 *01222* frameshift_variant c.407delT p.Leu136fs, *tRNA*$_{ser}$*::spec* | This work |
| JD0043 | JM1006 *murA* missense_variant c.487G>T p.Asp163Tyr, *tRNA*$_{ser}$*::spec* | This work |
| JD0044 | JM1006 *tRNA*$_{thr}$::P$_{mbrRAB}$*-luxAB-cat* | This work |
| JD0045 | JD0026 *tRNA*$_{thr}$::P$_{mbrRAB}$*-luxAB-cat* | This work |
| JD0046 | JM1006 *tRNA*$_{thr}$::P$_{mbrCD}$*-luxAB-cat* | This work |
| JD0047 | JD0026 *tRNA*$_{thr}$::P$_{mbrCD}$*-luxAB-cat* | This work |
| JD0048 | JM1006 *tRNA*$_{thr}$::P$_{mbrX}$*-luxAB-cat* | This work |
| JD0049 | JD0026 *tRNA*$_{thr}$::P$_{mbrX}$*-luxAB-cat* | This work |
| JD0050 | JM1006 *tRNA*$_{thr}$::P$_{mbrH}$*-luxAB-cat* | This work |
| JD0051 | JD0026 *tRNA*$_{thr}$::P$_{mbrH}$*-luxAB-cat* | This work |
| JD0052 | JM1006 *tRNA*$_{thr}$::P$_{mbrEFG}$*-luxAB-cat* | This work |
| JD0053 | JD0026 *tRNA*$_{thr}$::P$_{mbrEFG}$*-luxAB-cat* | This work |
| JD0054 | JM1006 *tRNA*$_{thr}$::P$_{comS}$*-luxAB-cat* | This work |
| JD0055 | JD0026 *tRNA*$_{thr}$::P$_{comS}$*-luxAB-cat* | This work |
| JD0056 | JD0026 Δ*mbrX::cat* | This work |
| JD0057 | JD0026 Δ*mbrH::cat* | This work |
| JD0058 | JD0026 Δ*mbrCD::cat* | This work |
| JD0059 | JD0026 Δ*mbrEFG::cat* | This work |
| JD0060 | JM1006 Δ*mbrRAB::spec* | This work |
| JD0061 | JM1006 *tRNA*$_{thr}$::P$_{mbrRAB\ mut}$*-luxAB-cat* | This work |
| JD0062 | JD0026 *tRNA*$_{thr}$::P$_{mbrRAB\ mut}$*-luxAB-cat* | This work |
| JD0063 | JM1006 *tRNA*$_{thr}$::P$_{mbrCD\ mut}$*-luxAB-cat* | This work |
| JD0064 | JD0026 *tRNA*$_{thr}$::P$_{mbrCD\ mut}$*-luxAB-cat* | This work |
| JD0065 | JD0060 *tRNA*$_{thr}$::P$_{mbrRAB}$*-luxAB-cat* | This work |
| JD0066 | JM1006 *mbrB*$_{R14P}$, *tRNA*$_{thr}$::P$_{mbrRAB}$*-luxAB-cat* | This work |
| JD0067 | JD0066, Δ*mbrA::lox72* | This work |
| JD0068 | JD0044 *mbrA*$_{Y14A}$*-spec* | This work |
| JD0069 | JD0044 *mbrA*$_{E153A}$*-spec* | This work |
| JD0070 | JD0044 *mbrA*$_{D195A}$*-spec* | This work |
| JD0071 | JD0045 *mbrA*$_{Y14A}$*-spec* | This work |
| JD0072 | JD0045 *mbrA*$_{E153A}$*-spec* | This work |
| JD0073 | JD0045 *mbrA*$_{D195A}$*-spec* | This work |
| JD0074 | JD0066 *mbrA*$_{D195A}$*-spec* | This work |
| JD0075 | JD0060 *P*$_{mbrRAB}$*::Sb-mbrR-mbrA-Lb-mbrB*$_{R14P}$-cat | This work |
| JD0076 | JD0060 *P*$_{mbrRAB}$*::Sb-mbrR-mbrA-Lb-mbrB*-cat | This work |
| JD0077 | JD0060 *P*$_{mbrRAB}$*::Sb-mbrR-mbrA*$_{D195A}$*-Lb-mbrB*$_{R14P}$-cat | This work |
| JD0078 | JD0060 *P*$_{mbrRAB}$*::Sb-mbrR-mbrA*$_{D195A}$*-Lb-mbrB*-cat | This work |
| JD0079 | JD0060 *P*$_{mbrRAB}$*::Sb-mbrR-mbrA-mbrB*$_{R14P}$-cat, *tRNA*$_{thr}$::P$_{sptA}$*-Lb-spec* | This work |

| | | |
|---|---|---|
| JD0080 | JD0060 *$P_{mbrRAB}$::mbrR-mbrA-mbrB$_{R14P}$-Lb-cat*, *tRNA$_{thr}$::*$P_{sptA}$*-Sb-spec* | This work |
| JD0081 | JD0060 Δ*mbrCD::cat* | This work |
| JD0082 | JD0081 Δ*mbrRAB ::lox72,* Δ*mbrCD ::lox72* | This work |
| JD0083 | JD0081 *$P_{mbrRAB}$::Sb-mbrR-mbrA-mbrB$_{R14P}$-cat*, $P_{mbrCD}$*::mbrC-Lb-MbrD-spec* | This work |
| JD0084 | JD0060 *$P_{mbrRAB}$::Sb-mbrR-mbrA-Lb (ΔmbrB::erm)* | This work |
| *Escherichia coli* | | |
| Top10 | *mcrA*, Δ(*mrr-hsd*RMS-*mcrBC*), Phi80*lacZ(del)M15*, Δ*lacX74*, *deoR*, *recA1*, *araD139*, Δ(*ara-leu*)7697, *galU*, *galK*, *rpsL(SmR)*, *endA1*, *nupG* | Invitrogen |

### Table S6. List of plasmids used in this study.

| Names | Characteristics | Reference/source |
|---|---|---|
| pGIR311 | $Amp^r$; pBADHisA encoding LarAH3 | [4] |
| pGhostcre | Thermosensitive replication vector in *S. salivarius*, encoding the Cre recombinase; $ery^R$ | [5] |
| pGILF*spec* | pG+host9 derivative containing the spectinomycin resistance cassette $P_{spec}$-*spec* downstream of *luxAB* | [5] |
| pNZ5319 | pACYC184 derivative containing the cat gene under the control of the P32 constitutive promoter from *Lactococcus lactis* | [6] |
| pGIUD0855ery | pUC18 derivative containing the *erm* gene | [5] |
| pBAD-*mbrR_strep* | pGIR311 derivative encoding MbrR fused to a C-terminal StrepTag | This work |

### Table S7. List of synthetic DNA fragments used in this study.

| Names | Characteristics | Reference/source |
|---|---|---|
| Construct 1 | *$P_{mbrRAB}$::Sb-MbrR-mbrA-Lb-mbrB$_{R14P}$-cat* | This work |
| Construct 2 | *$P_{mbrRAB}$::Sb-MbrR-mbrA-mbrB$_{R14P}$-cat* | This work |
| Construct 3 | *$P_{mbrCD}$::-MbrC-Lb-mbrD-spec* | This work |

The sequences of the small fragment of NanoLuc luciferase (Sb) with a linker for N-terminal fusion, and the large fragment (Lb) with a linker for C-terminal fusion, were obtained from the JW Veening lab [7]. These NanoLuc luciferase sequences are optimized for streptococci and are derived from Dixon *et al.,* 2016 [8].

**Table S8. List of oligonucleotides used in this study.**

| Names | Sequences |
|---|---|
| JD382_UF_gntR | 5'-TCAAGGTCCGTTGTCATAAC-'3 |
| JD383_DR_gntR | 5'-AAGACAACTTCATCTAGGAC-'3 |
| JD384_UF_aroC | 5'-TGATGGGGTTTTGATTGATC-'3 |
| JD385_DR_aroC | 5'-TGACTGATTTGACTGGTTAC-'3 |
| JD387_UF_murA | 5'-AAGGCCAAAGAAGTCCTTGG-'3 |
| JD388_DR_murA | 5'-TAGGGTTTGCTAGATACCTT-'3 |
| JD396_DR_01360 | 5'-ATCATCAAAGAAGTGGGACT-'3 |
| JD445_F_gntR (lox66) | 5'-ATCCTTATGGGATTTATCTTCCTTAGGAAAGGACTTGTAGACCG-'3 |
| JD446_R_gntR (lox71) | 5'-ATTACATTCCCTTTAGTAACGTGAATGTCCAAGCCATAAGAGAC-'3 |
| JD449_R_01359 (lox71) | 5'-TACATTCCCTTTAGTAACGTGAAAATATCAGTCATCGGTCTAC-'3 |
| JD450_F_01359 (lox66) | 5'-CCTTATGGGATTTATCTTCCTTATTAAATTTAGACTAAAGAAA-'3 |
| JD476_F_01607 | 5'-CTAGGCAAAAAACATGGCTC-'3 |
| JD478_R_01803533 | 5'-AGATTTTCCTTGAAATCGTC-'3 |
| JD482_R_PgntR (ATG_luxAB) | 5'-TAAGCAAAAAGTTTCCAAATTTCATAAGAGACTCCTTTCTAAAAG-'3 |
| JD488_DR_01607 | 5'-AGCTGCTGTTGATAATTTAG-'3 |
| JD491_UF_01609 | 5'-CTTAACCAACTTTATGAACC-'3 |
| JD498_R_priA | 5'-TGTCGTATCAACATCCATAC-'3 |
| JD499_F_priA | 5'-GTGAATTGGCACGTTTGAAA-'3 |
| JD500_F_mur1B | 5'-GAATCAGCAGAAATTGTAAG-'3 |
| JD501_R_mur1B | 5'-CATAAGCACTCCTTTTGTTG-'3 |
| JD502_F_1599160 | 5'-ATATTCCCTTGACCATCTTC-'3 |
| JD503_R_1599160 | 5'-ACAAGTTGCTCAATGACCTA-'3 |
| JD504_F_DUF948 | 5'-AATTTATTCGCCAGCAGATG-'3 |
| JD505_R_DUF948 | 5'-TCTGCAATCTTTCGCATCTC-'3 |
| JD514_F_00143 | 5'-TAAAATCGTTTCACCAGTAG-'3 |
| JD515_R_00143 | 5'-TTTGATATCAGCACCTGAAG-'3 |
| JD516_F_00896 | 5'-GGGATTAATCATTTTCATGG-'3 |
| JD517_R_00896 | 5'-AGACGTCTACCAAACGATTG-'3 |
| JD518_R_01222 | 5'-TTTAATGTATCTCTCCTTCG-'3 |
| JD519_F_01222 | 5'-ATCGAAAGTTATTCGGTTTC-'3 |
| JD520_R_01305 | 5'-TAATCTTACTCTTATCTGCG-'3 |
| JD521_F_01305 | 5'-TCTTTGAGTGAGTCTTTATC-'3 |
| JD522_R_1370 | 5'-AACTTAATCTTGGCCAGAAG-'3 |
| JD523_F_01370 | 5'-GTTAGGTCCTCTAGTTTAAC-'3 |
| JD524_R_corA1 | 5'-GTATCGAACTTCTCTTGTTC-'3 |
| JD525_F_corA1 | 5'-TTTAGTCGATAAGCAAGGTG-'3 |
| JD526_F_valS | 5'-TAAGGTTGATTTCTACCCAC-'3 |
| JD527_R_valS | 5'-CATACATACTTAAAATGGCG-'3 |
| JD532_F_pgntR2 (tRNAthr) | 5'-TGAAAAATTGAGAGGTATAAATCAATTGGACTCAAACAGGACTTC-'3 |
| JD557_F_01359 (spec) | 5'-ATTACATTCCCTTTAGTAACGTGAAATGTTCCACAACTAGGAGG-'3 |
| JD558_R_01360 (spec) | 5'-ATCCTTATGGGATTTATCTTCCTTATTTCCAAAACATGCTCATAC-'3 |
| JD559_F_P01608 (tRNAthr) | 5'-TGAAAAATTGAGAGGTATAAATCAACAGTGATTTTCCTCCTATTTATTC-'3 |
| JD560_R_P01608 (ATG_luxAB) | 5'-TAAGCAAAAAGTTTCCAAATTTCATCTTTTATCCTTCTTGAGAATC-'3 |
| JD575_R_P01430 (luxAB) | 5'-TAAGCAAAAAGTTTCCAAATTTCATGTTAAGTCCTCCAAAATAAAC-'3 |
| JD576_F_P01430 (up tRNAthr) | 5'-TGAAAAATTGAGAGGTATAAATCAAGTCTTTTGAGATTTGAGTTTTG-'3 |
| JD577_R_P01755 (luxAB) | 5'-TAAGCAAAAAGTTTCCAAATTTCATGAGAATCCACCTCTTTTCAT-'3 |
| JD578_F_P01755 (up tRNAthr) | 5'-TGAAAAATTGAGAGGTATAAATCAAGAGCCTATATTATCTTGAAA-'3 |
| JD579_R_P00448 (luxAB) | 5'-TAAGCAAAAAGTTTCCAAATTTCATGGAAGCCAATTCCTCCAATG-'3 |
| JD580_F_P00448 (tRNAthr) | 5'-TGAAAAATTGAGAGGTATAAATCAAAACGTCAGATTCGTCCCAATC-'3 |
| JD582_UR_01608 (cat) | 5'-TCTACATTCCCTTTAGTAACGTGAAAAAACGAATCATCTTTTATC-'3 |
| JD583_UF_01755 | 5'-TCTAGGGTATTTTATTGGTC-'3 |
| JD584_DR_01755 | 5'-CAAATACCAATACTACTGTG-'3 |
| JD587_DF_01755 (cat) | 5'-TCTACATTCCCTTTAGTAACGTGAACCATATATCAAATAAAGCTC-'3 |
| JD588_UF_00446 | 5'-CTGAAACTAATCGTTTCAGA-'3 |
| JD589_DR_00448 | 5'-TCTTCGATAGCTTTTACAAG-'3 |
| JD593_UF_01430 | 5'-CAGATTGGTTCAAGCTTTAA-'3 |
| JD594_DR_01430 | 5'-GCTTCCTTGAAAAAGTTTAC-'3 |
| JD599_R_01755 (cat) | 5'-GCCCTTATGGGATTTATCTTCCTTATGACCCTTTGTTCATGAGAA-'3 |
| JD600_R_01607-8 (cat) | 5'-GCCCTTATGGGATTTATCTTCCTTAATGTTTTTTGCCTAGAACAA-'3 |
| JD601_Probe_EMSA_F_Cy3_PgntR | 5'-Cy3-TTAGTTGTATCATTATGGTAGTACAAAAAT-'3 |

| | |
|---|---|
| JD602_Probe_EMSA_R_PgntR | 5'-ATTTTTGTACTACCATAATGATACAACTAA-'3 |
| JD603_P_EMSA_R_P01607-8 | 5'-TTTATTGTATCATTTCAAAGGTACAATTTC-'3 |
| JD604_P_EMSA_F_Cy3_P01607-8 | 5'-Cy3-GAAATTGTACCTTTGAAATGATACAATAAA-'3 |
| JD605_P_EMSA_F_Cy3_P01755 | 5'-Cy3-TTCGTTGTATTATAATAAAGATACAAATAA-'3 |
| JD606_P_EMSA_R_P01755 | 5'-TTATTTGTATCTTTATTATAATACAACGAA-'3 |
| JD607_P_EMSA_R_P00446 | 5'-TCTTTTGTATCACATAAATAAAACAATTGT-'3 |
| JD608_P_EMSA_F_Cy3_P00446 | 5'-Cy3-ACAATTGTTTTATTTATGTGATACAAAAGA-'3 |
| JD609_P_EMSA_F_Cy3_P01430 | 5'-Cy3-AAAATTGTATCACTGTGCTATAATAAAGCA-'3 |
| JD610_P_EMSA_R_P01430 | 5'-TGCTTTATTATAGCACAGTGATACAATTTT-'3 |
| JD611_P_EMSA_F_Cy3_P01607-8_mut | 5'-Cy3-GAAATTGTACCTTTGAAATGATATAATAAA-'3 |
| JD612_P_EMSA_R_P01607-8_mut | 5'-TTTATTATATCATTTCAAAGGTACAATTTC-'3 |
| JD616_R_01430 (cat) | 5'-GCCCTTATGGGATTTATCTTCCTTAAAGTGCTGCTTGCATGTTAA-'3 |
| JD621_EMSA_F_CY3_PgntR_mut | 5'-Cy3-TTAGTTATATCATTATGGTAGTAAAAAAAT-'3 |
| JD622_EMSA_R_PgntR_mut | 5'-ATTTTTTTACTACCATAATGATATAACTAA-'3 |
| JD623_R_00446_2 (cat) | 5'-CCCTTATGGGATTTATCTTCCTTATTTCTTTCGCATAAGTCTC-'3 |
| JD632_F_PgntR_mut | 5'-GTATTTTAGTTTTATCATTATGGTAGTAAAAAAATAATAC-'3 |
| JD633_R_PgntR_mut | 5'-GTATTATTTTTTTACTACCATAATGATAAAACTAAAATAC-'3 |
| Fw.UptRNAser-uplox66 | 5'-CTGTTAAAACCATCTTCTTTTTAATTACTTAAGGAAGATAAATCCCATAAG-'3 |
| UF_tRNAthr | 5'-TGTCAAAGGATTAGGAAAAC-'3 |
| UR_tRNAthr | 5'-TTGATTTATACCTCTCAATTT-'3 |
| F_luxAB_ATG | 5'-ATGAAATTTGGAAACTTTTTGC-'3 |
| R_tRNAthr | 5'-AAGGAGAAAATTATGTACAC-'3 |
| UF_tRNAser | 5'-CAAGATTAACCATGACCTTC-'3 |
| DR_tRNAser2 | 5'-TTGGATAAGGTCTTGACTTC-'3 |
| Rv.lox71 | 5'-TTCACGTTACTAAAGGGAATGTA-'3 |
| Fw.lox66 | 5'-TAAGGAAGATAAATCCCATAAGG-'3 |
| F_PcomS_tRNAthr | 5'-AAATTGAGAGGTATAAATCAACTGCAGAAAATTACAATAAG-‘3 |
| R_PcomS_luxAB_TTG | 5'-GCAAAAAGTTTCCAAATTTCAAAATAAAACTCCTTTTAAC-‘3 |
| LP93_F_up_gntR | 5’-AAATCGTGTTGAGGTAGTC-‘3 |
| LP94_R_Dn_01360 | 5’-TCCAAGACCCTGCGGCTAG-‘3 |
| LP96_R_dn_01607 | 5’-CTGCTGTTGATAATTTAGC-‘3 |
| LP97_F_01359_D195A | 5’-GCAGATATTGAATCCGTCCTAGCTGAAGTTGTCTTCTTAAAATAC-‘3 |
| LP98_R_01359_D195A | 5’-GTATTTTAAGAAGACAACTTCAGCTAGGACGGATTCAATATCTGC-‘3 |
| LP101_F_01360_P14R | 5’-GAATTACAATCGGTCCGTAAATGGTATCTTGGCATC-‘3 |
| LP102_R_01360_P14R | 5’-GATGCCAAGATACCATTTACGGACCGATTGTAATTC-‘3 |
| LP138_R_01608_LgBit | 5’-CTGAGCCGCCGCCGCCTGATGAGCCTACCTCCTCAATCTTACGTC-‘3 |
| LP145_F_01359 | 5’-ATGACTGATATTGTAACTCTAAC-‘3 |
| LP159_F_01359_Y14A | 5’-CTAACAAATCTCACCAAGACCGCTAATGGTATTCCAGCCCTTAG-‘3 |
| LP160_R_01359_Y14A | 5’-CTAAGGGCTGGAATACCATTAGCGGTCTTGGTGAGATTTGTTAG-‘3 |
| LP161_F_01359_E153A | 5’-CTCGATGCTCCTATCGGTGGAGTCGATCC-‘3 |
| LP162_R_01359_E153A | 5’-GATAGGAGCATCGAGGAGATAGAGTTTAGC-‘3 |
| LP210_LgBit_ery | 5’-ATGAAAATTCCTCCGGGTTACTATGAGTTGATTGTTACAC-‘3 |
| LP211_dnMbrB_ery | 5’-TATTTAACGGGAGGAAATAGAGAAAAACCGTGTTCCTTC-‘3 |
| F_PpotA2_tRNAthr | 5’-AAATTGAGAGGTATAAATCAATCATTTTGGAAGCAAAATAC-3’ |
| GH80_Rv_PsptA | 5’-ATCTTGATTTCTCCAATTTGAT-3’ |
| GH148_F_Smbit_Nter | 5’-TTAAGGAGGCAAATATGGTA-‘3 |
| GH66_R_ery_SplitLuc | 5’-CTATTTCCTCCCGTTAAATA-3’ |

#### Table S9. List of PCR fragments used to generate mutants in this study.

| PCR fragment | Primer 1 | Primer 2 |
|---|---|---|
| Upstream region of *mbrR* | JD382_UF_gntR | JD446_R_gntR (lox71) |
| *Spec*$^R$ cassette | FW.lox66 | RV.lox71 |
| *Cat* cassette | FW.lox66 | RV.lox71 |
| Downstream region of *mbrR* | JD445_F_gntR (lox66) | JD396_DR_01360 |
| Upstream region of *mbrA* | JD382_UF_gntR | JD449_R_01359 (lox71) |
| Downstream region of *mbrB* | JD450_F_01359 (lox66) | JD396_DR_01360 |
| *mbrAB* amplification | JD382_UF_gntR | JD396_DR_01360 |
| Upstream region of $tRNA_{ser}$ | UF_tRNAser | Fw.UptRNAser-uplox66 |
| Spec downstream region of $tRNA_{ser}$ | FW.lox66 | DR_tRNAser |
| Spec cassette at $tRNA_{ser}$ fragment used for co-transformation | UF_tRNAser | DR_tRNAser |
| *1803533* amplification | JD476_F_01607 | JD478_R_01803533 |
| *priA* amplification | JD498_R_priA | JD499_F_priA |
| *mur1B* amplification | JD500_F_mur1B | JD501_R_mur1B |
| *1599160* amplification | JD502_F_1599160 | JD503_R_1599160 |
| *aroF* amplification | JD384_UF_aroC | JD385_DR_aroC |
| *aroF* amplification | JD384_UF_aroC | JD385_DR_aroC |
| *00624* amplification | JD504_F_DUF948 | JD505_R_DUF948 |
| *00896* amplification | JD516_F_00896 | JD517_R_00896 |
| *valS* amplification | JD526_F_valS | JD527_R_valS |
| *corA1* amplification | JD525_R_corA1 | JD525_F_corA1 |
| *00143* amplification | JD514_F_00143 | JD515_R_00143 |
| *01305* amplification | JD520_R_01305 | JD521_F_01305 |
| *01370* amplification | JD522_R_1370 | JD523_F_01370 |
| *01222* amplification | JD518_R_01222 | JD519_F_01222 |
| *murA* amplification | JD387_UF_murA | JD388_DR_murA |
| Upstream region of $tRNA_{thr}$ | UF_tRNAthr | UR_tRNAthr |
| Promoter of *mbrR* for *luxAB* fusion | JD482_R_PgntR (ATG_luxAB) | JD532_F_pgntR2 (tRNAthr) |
| *luxAB*- downstream region of $tRNA_{thr}$ | F_luxAB_ATG | DR_tRNAthr |
| Promoter of *mbrCD* for *luxAB* fusion | JD559_F_P01608 (tRNAthr) | JD560_R_P01608 (ATG_luxAB) |
| Promoter of *mbrX* for *luxAB* fusion | JD578_F_P01755 (up tRNAthr) | JD577_R_P01755 (luxAB) |
| Promoter of *mbrH* for *luxAB* fusion | JD576_F_P01430 (up tRNAthr) | JD575_R_P01430 (luxAB) |
| Promoter of *mbrEFG* for *luxAB* fusion | JD579_R_P00448 (luxAB) | JD580_F_P00448 (tRNAthr) |
| Promoter of *comS* for *luxAB* fusion | F_PcomS_tRNAthr | R_PcomS_luxAB_TTG |
| Upstream region of *mbrX* | JD583_UF_01755 | JD599_R_01755 (cat) |
| $P_{32}$-*cat* cassette amplification | 1331 Fw.lox66 | 1330 Rv.lox71 |
| Downstream region of *mbrX* | JD587_DF_01755 (cat) | JD584_DR_01755 |
| Upstream region of *mbrH* | JD593_UF_01430 | JD616_R_01430 (cat) |
| Downstream region of *mbrH* | JD595_DF_01430 | JD594_DR_01430 |
| Upstream region of *mbrCD* | JD448_DR_01607 | JD600_R_01607-8 (cat) |
| Downstream region of *mbrCD* | JD582_UR_01608 (cat) | JD491_UF_01609 |
| Upstream region of *mbrEFG* | JD588_UF_00446 | JD623_R_00446_2 (cat) |
| Downstream region of *mbrEFG* | JD590_DF_00448 | JD589_DR_00448 |
| Upstream region of *mbrB* | JD382_UF_gntR | JD558_R_01360 (spec) |
| Downstream region of *mbrB* | JD448_F_01359 (lox 71) | JD396_DR_01360 |
| Upstream region of *mbrA* | JD382_UF_gntR | JD447_R_01359 (lox 66) |
| Downstream region of *mbrA* | JD557_F_01359 (spec) | JD396_DR_01360 |
| Upstream part P*mbrR-mut-*$tRNA_{thr}$ | 682 UF_tRNAthr | JD632_F_PgntR_mut |
| Downstream part P*mbrR-mut-luxAB* $tRNA_{thr}$ | JD633_R_PgntR_mut | 685 DR_tRNAthr |
| Promoter of *mbrR* annealing for EMSA | JD601_probe_EMSA_F_Cy3_PgntR | JD602_Probe_EMSA_R_PgntR |
| Promoter of *mbrR*-mut annealing for EMSA | JD621_EMSA_F_CY3_PgntR_mut | JD622_EMSA_R_PgntR_mut |
| Promoter of *mbrCD* annealing for EMSA | JD604_P_EMSA_F_Cy3_P01607-8 | JD603_P_EMSA_R_P01607-8 |
| Promoter of *mbrCD*-mut annealing for EMSA | JD611_P_EMSA_F_Cy3_P01607-8_mut | JD612_P_EMSA_R_P01607-8_mut |
| Promoter of *mbrX* annealing for EMSA | JD605_P_EMSA_F_Cy3_P01755 | JD606_P_EMSA_R_P01755 |
| Promoter of *mbrH* annealing for EMSA | JD609_P_EMSA_F_Cy3_P01430 | JD610_P_EMSA_R_P01430 |
| Promoter of *mbrEFG* annealing for EMSA | JD608_P_EMSA_F_Cy3_P00446 | JD607_P_EMSA_R_P00446 |
| Upstream $mbrA_{Y14A}$ | JD382_UF_gntR | LP160_R_01359_Y14A |
| Downstream $mbrA_{Y14A}$ | LP159_F_01359_Y14A | LP94_R_Dn_01360 |
| Upstream $mbrA_{D195A}$ | JD382_UF_gntR | LP98_R_01359_D159A |
| Downstream $mbrA_{D195A}$ | LP97_F_01359_D159A | LP94_R_Dn_01360 |
| Upstream $mbrA_{E153A}$ | JD382_UF_gntR | LP162_R_01359_E153A |
| Downstream $mbrA_{E153A}$ | LP161_F_01359_E153A | LP94_R_Dn_01360 |
| Promoter of *sptA* for sBI7 induce expression | F_PpotA2_tRNAthr | GH80_Rv_PsptA |
| *Erm* cassette | GH148_F_Smbit_Nter | GH66_R_ery_SplitLuc |
