## Supplementary material for "A streamlined ABC extruder-repressor module drives multi-bacteriocin resistance in streptococci": Dataset S1

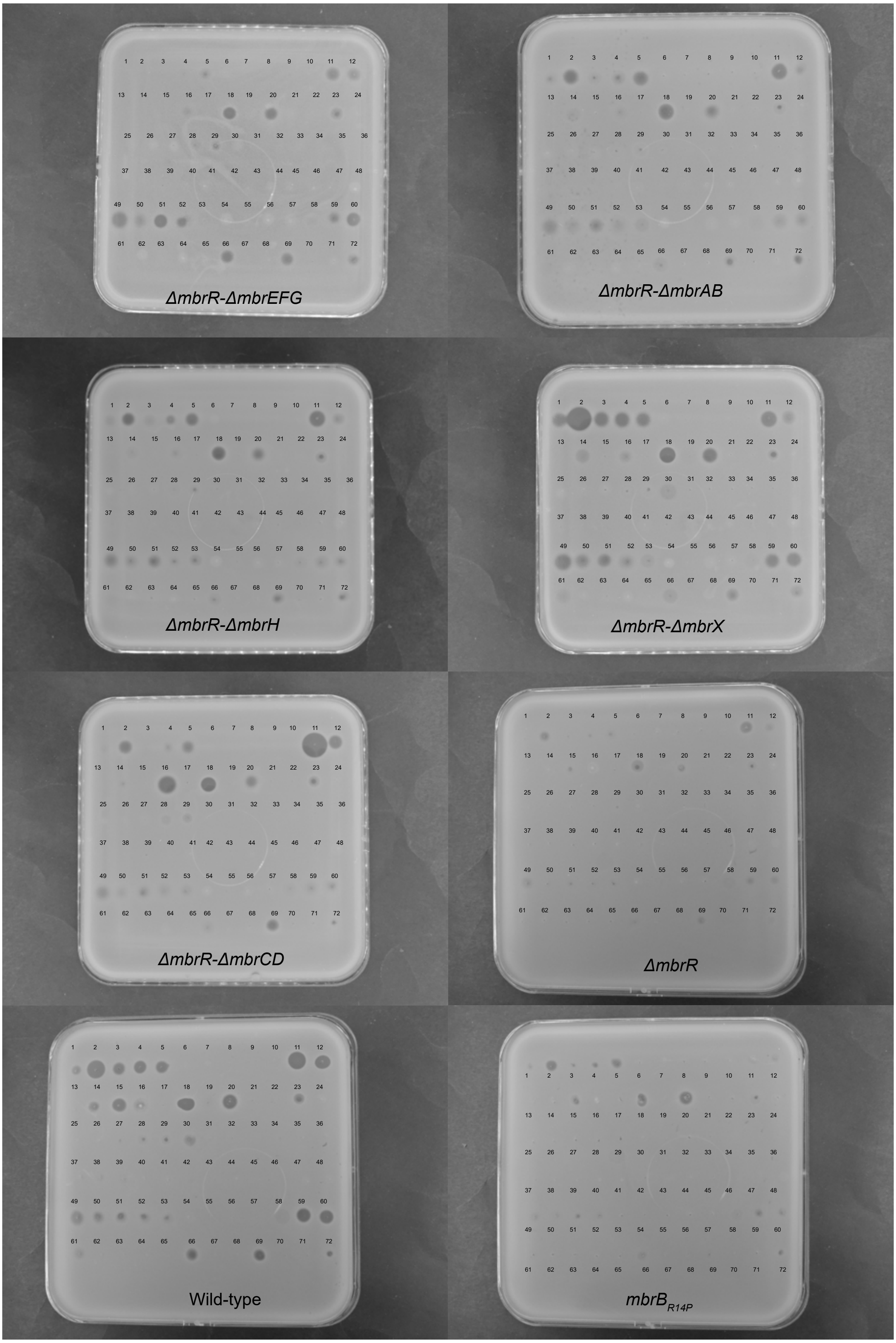


1 Cerein H- A+B+C
2 Garvicidin KS-A+B+C
3 Aureocin A70A+B+C+D
4 Cerein V-A+B+C
5 Cerein X-A+B+C
6 Enterocin E760
7 Brochocin C-Beta
8 -
9 -
10 BlpK_sp
11 BlpK
12 SlvV
13 -
14 Lacticin F-A + F-X
15 Acidocin LF221B
16 Sakacin Tα +Tβ
17 Microcin 24
18 Plantaricin Sα+Sβ
19 Lacticin Z_C*
20 Mutacin BHTB
21 Garvieacin Q
22 Lactococcin A
23 Plantaricin E + F
24 SlvY
25 Lactococcin A*
26 Lactococcin B
27 Lactococcin Qα+Qβ
28 Lactococcin Gα+Gβ
29 Plantaricin NC8alpha +beta
30 SlvZ
31 Mundticin L
32 Ent35
33 Avicin A
34 Sakacin X
35 Piscicolin 126
36 -
37 Leucocin C
38 Ubericin A
39 EntA
40 PedA1
41 Plantaricin 423
42 Leucocin A
43 Sakacin A
44 Enterocin P
45 Carnobacteriocin BM1
46 Hiracin JM79/Bac43
47 Ent50-52
48 OF 7
49 Lacticin Z
50 Lacticin Q
51 Epidermicin Ni01
52 Aureocin A53
53 Weisselin 110
54 LsbA
55 LsbB
56 Enterocin K1
57 Enterocin EJ97
58 Bactofencin A
59 Enterocin 7A+7B
60 EntL50A+B
61 Weisselin M
62 Weisselin Y
63 SlvW
64 Bacteriocin 32
65 Lacticin Z_B*
66 Ent1071A+B
67 -
68 L-1077
69 SlvV*
70 C7 SN
71 ColV SN
72 Nisin

Sensitivity assays of MbrR regulon to PARAGEN collection.
Fig. 1C, 4C, S3: full-size pictures

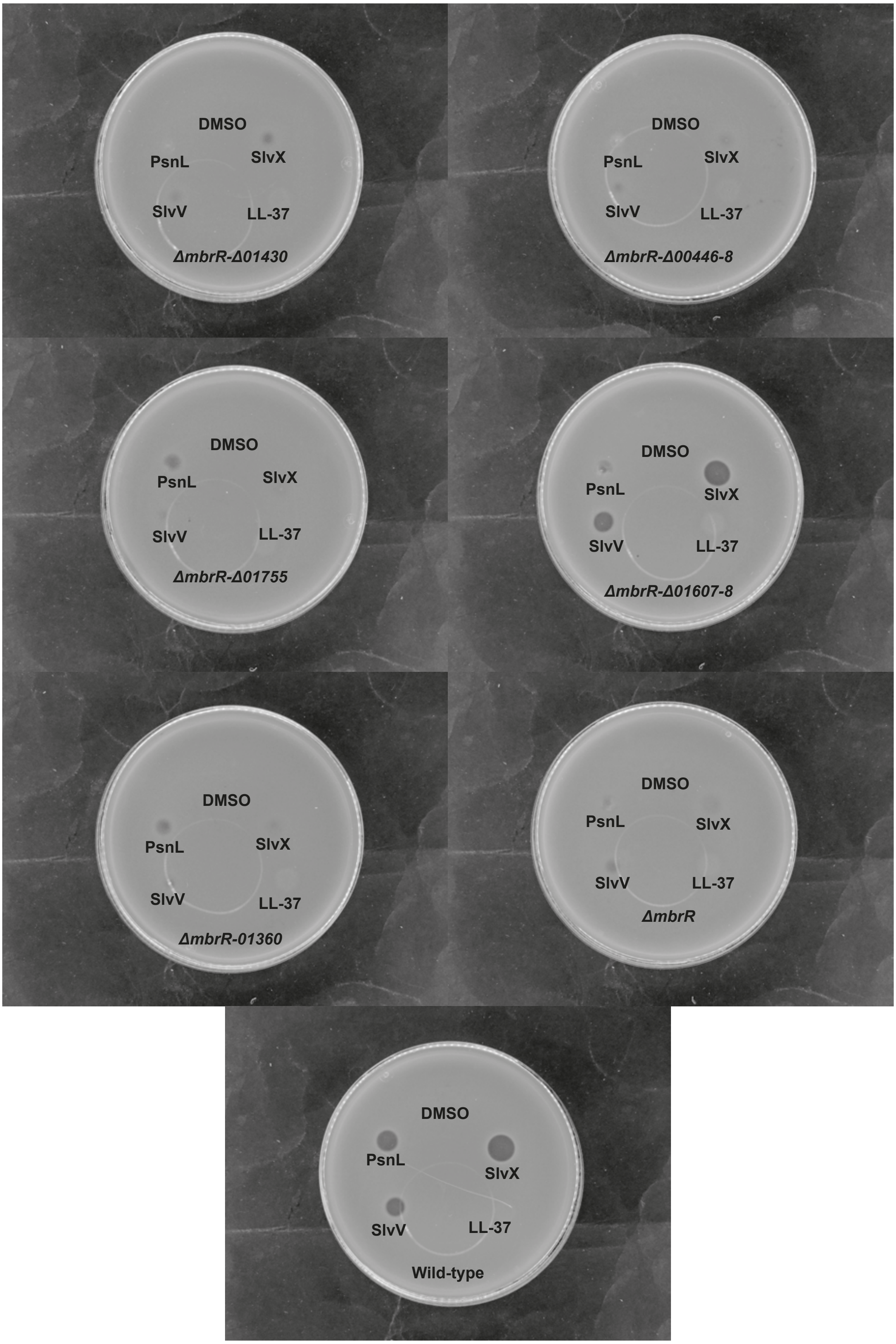


Sensitivity assays of MbrR regulon to salivaricins
Fig. 1C, 4C, S3: full-size pictures

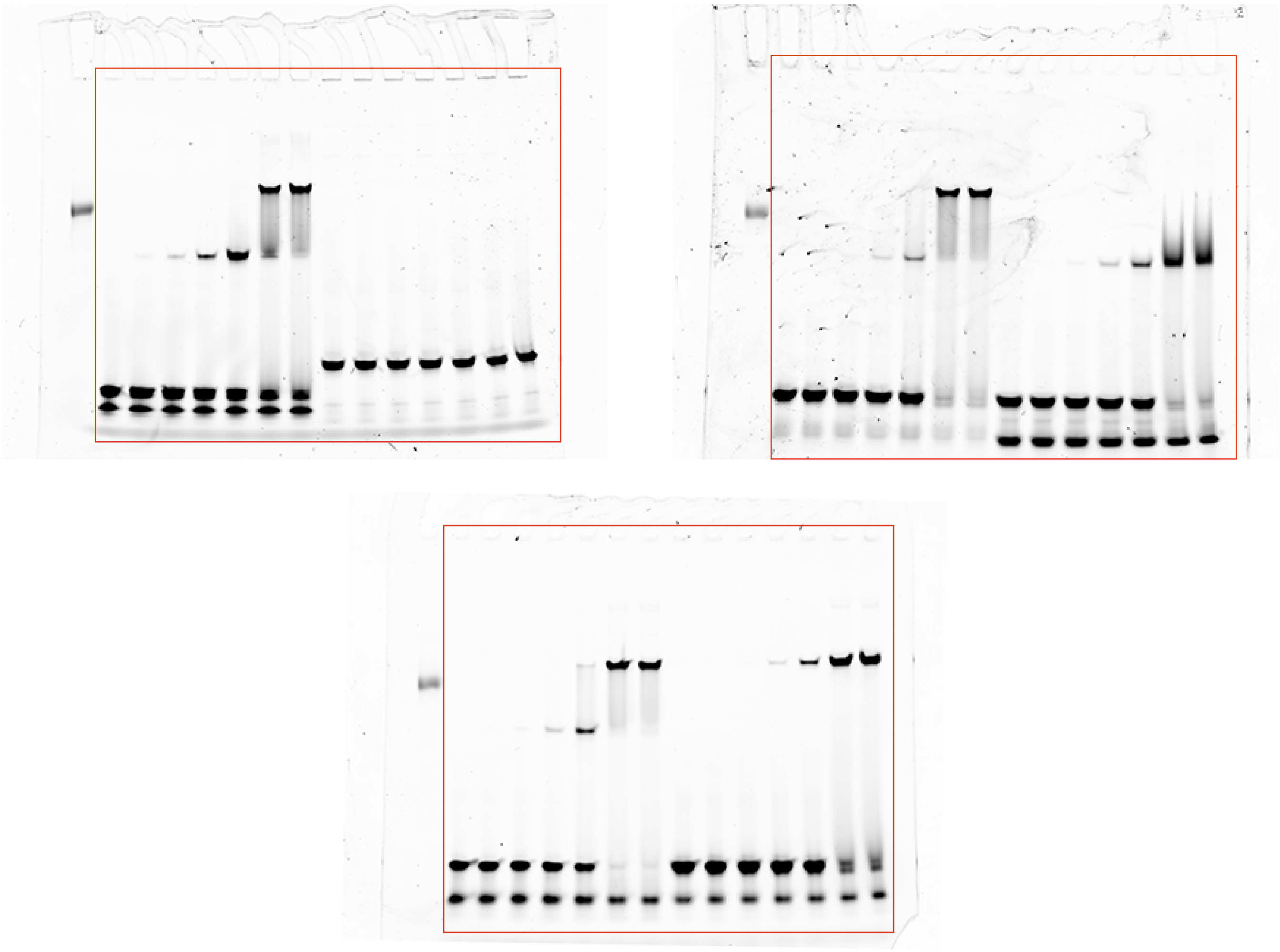

Fig. 3D: full-size pictures

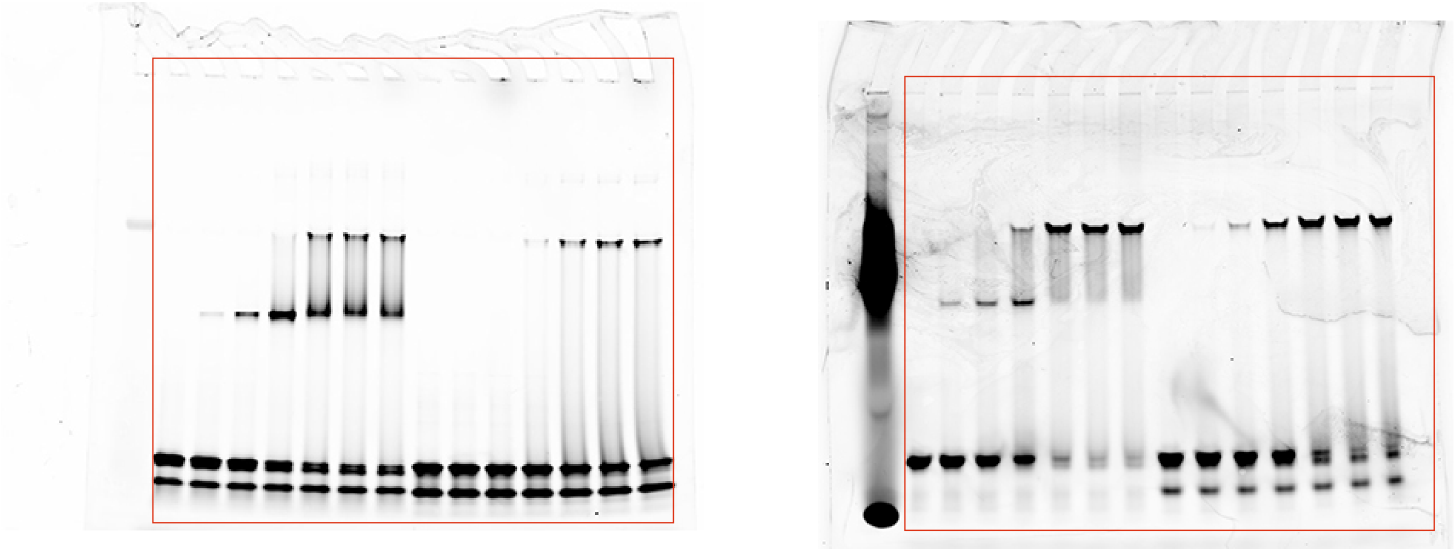

Fig. 3E: full-size pictures

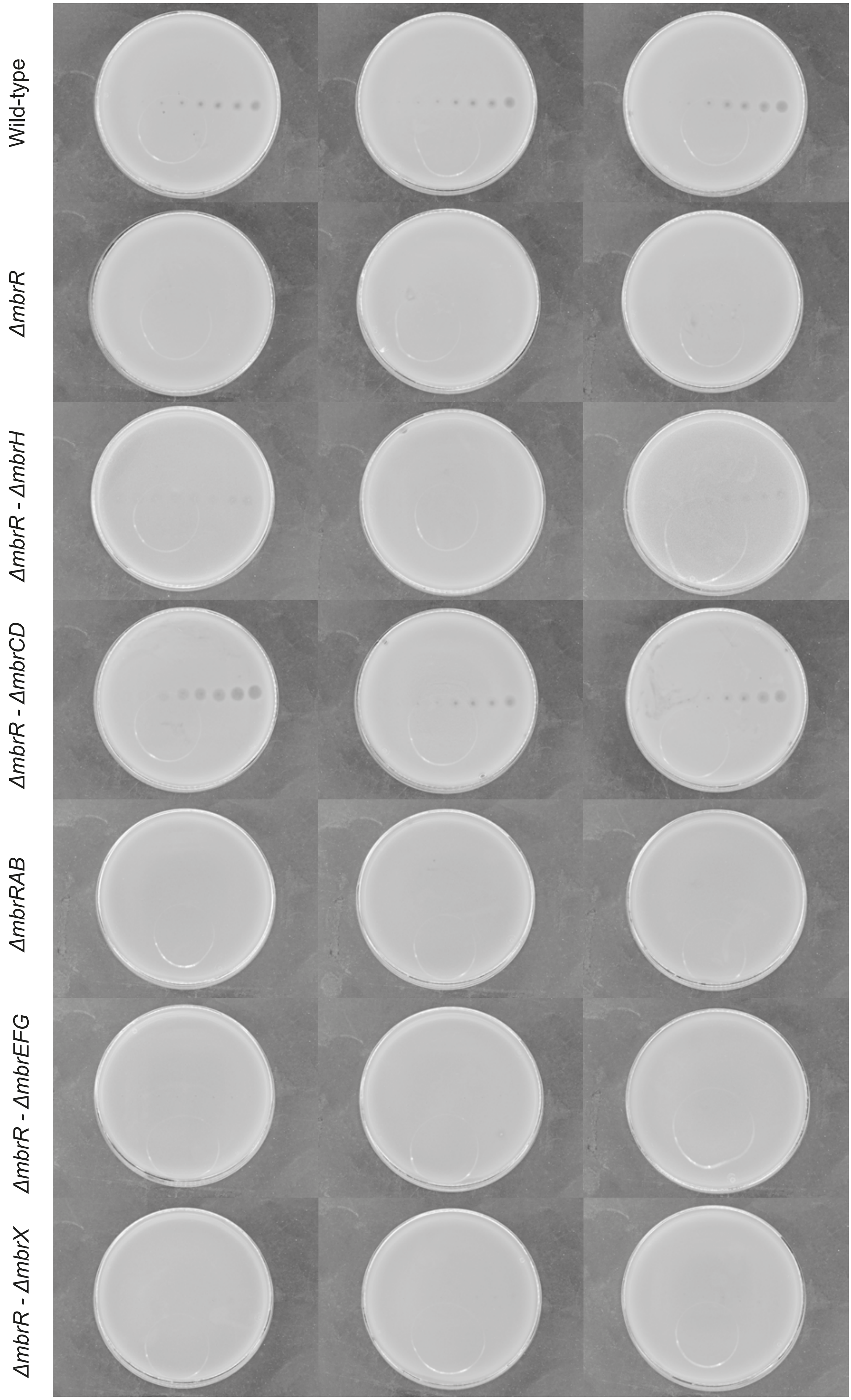


Triplicates of sensitivity assays of MbrR regulon to sBlpK.
Fig. 4A: full-size pictures
